# Atypical Protein Kinase C Promotes its own Asymmetric Localisation by Phosphorylating Cdc42 in the *C. elegans* zygote

**DOI:** 10.1101/2023.10.27.563985

**Authors:** John Packer, Alicia G. Gubieda, Aaron Brooks, Lars N. Deutz, Iolo Squires, Shona Ellison, Claudia Schneider, Sundar Ram Naganathan, Adam J.M. Wollman, Daniel J. Dickinson, Josana Rodriguez

## Abstract

Atypical protein kinase C (aPKC) is a major regulator of cell polarity. Acting in conjunction with Par6, Par3 and the small GTPase Cdc42, aPKC becomes asymmetrically localised and drives the polarisation of cells. aPKC activity is crucial for its own asymmetric localisation, suggesting a hitherto unknown feedback mechanism contributing to polarisation. Here we show in the *C. elegans* zygote that the feedback relies on aPKC phosphorylation of Cdc42 at serine 71. The turnover of CDC-42 phosphorylation ensures optimal aPKC asymmetry and activity throughout polarisation by tuning Par6/aPKC association with Par3 and Cdc42. Moreover, turnover of Cdc42 phosphorylation regulates actomyosin cortex dynamics that are known to drive aPKC asymmetry. Given the widespread role of aPKC and Cdc42 in cell polarity, this form of self-regulation of aPKC may be vital for the robust control of polarisation in many cell types.

**Key findings/graphical abstract:**

- Phosphorylation of CDC-42 by aPKC accelerates aPKC dissociation from CDC-42, limiting aPKC activity
- CDC-42/aPKC dissociation promotes aPKC association with PAR-3 and, thereby, aPKC asymmetry due to actomyosin flow
- Cycling of CDC-42 phosphorylation fuels the exchange of aPKC between anteriorly transported PAR-3 and aPKC-active CDC-42 complexes
- Turnover of CDC-42 phosphorylation alternates its association with effectors, aPKC and MRCK-1, ensuring proper actomyosin dynamics

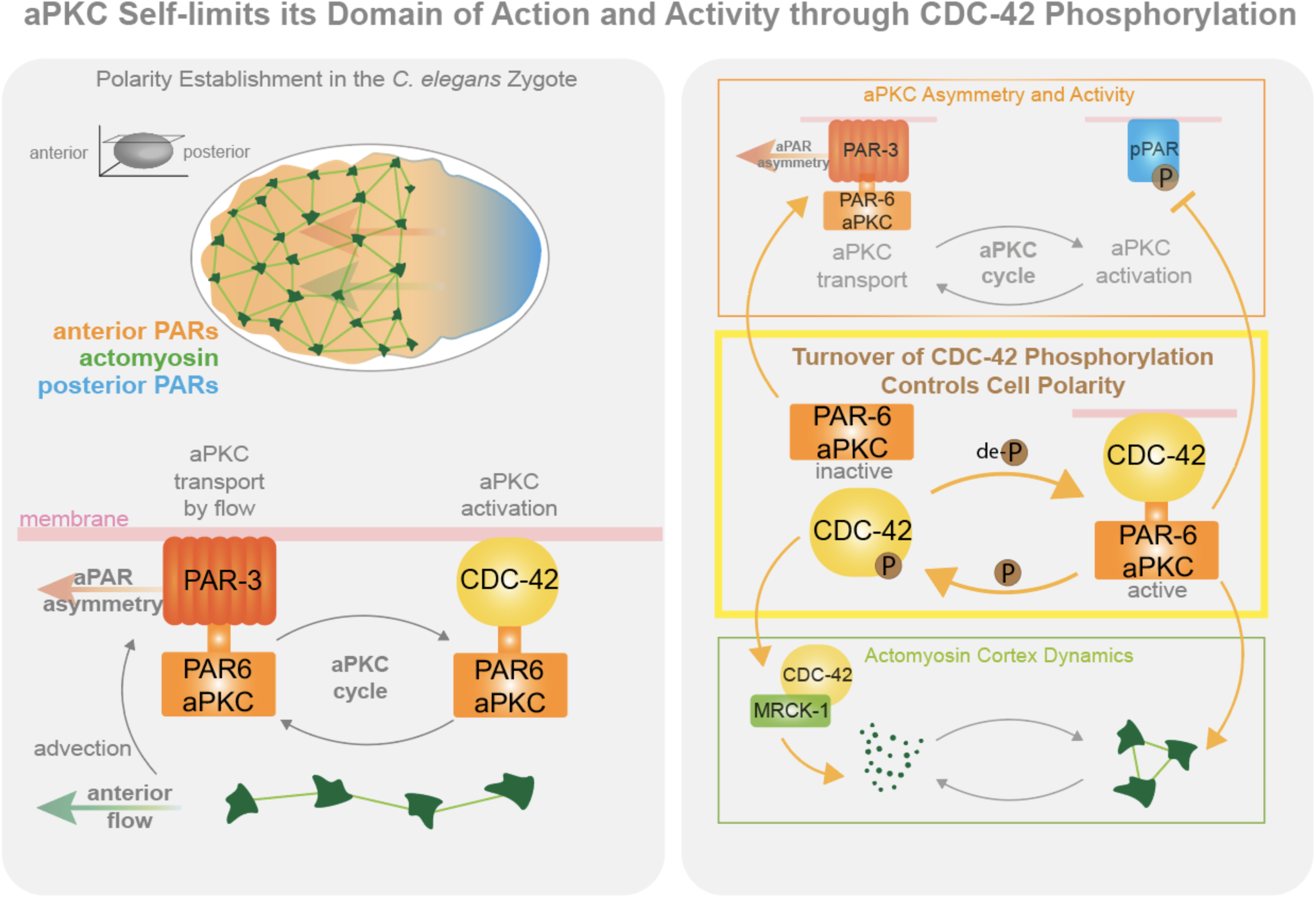

## INTRODUCTION

The polarisation of cells, or asymmetric distribution of cellular components and functions, is key to the generation of cellular diversity. Cell polarity depends on the asymmetric localisation of molecules within the cell.^1–3^ Prominent examples are the PAR proteins, conserved effectors of cell polarity, initially identified in *Caenorhabditis elegans.*^4–6^ In this system, atypical protein kinase C (aPKC), the small guanosine triphosphatase (GTPase) Cdc42, and the PDZ domain scaffold proteins Par3 and Par6 work together to polarise the zygote, leading to an asymmetric cell division. These same factors also play key roles in other polarizing cell types: for example, they specify the apical domain in epithelial cells,^7,8^ the front in migrating cells^9,10^ and the domain of an asymmetrically dividing cell that will be inherited by one of the daughter cells.^11,12^

aPKC is an essential component of this polarising network,^13^ its asymmetric membrane localisation and activity being controlled by the other network members. aPKC activity is regulated by both Par6 and Par3, which have been shown to enhance or suppress aPKC catalytic activity depending on the system studied.^14–24^ Meanwhile, a consensus has emerged that Cdc42, when in its active GTP-bound state, can recruit PAR-6/aPKC to the membrane and promote aPKC kinase activity.^21–23,25,26^ Par6 is considered an aPKC co-factor that brings aPKC to the membrane through its interaction with either Par3 or Cdc42, which can compete for PAR-6/aPKC membrane recruitment.^16,17,22,25,27–36^ Recent evidence suggests that aPKC can also be directly recruited to the membrane by binding to Par3^37,38^ and/or via a polybasic domain in the aPKC protein sequence that is capable of interacting with membrane lipids directly.^39,40^ How these various mechanisms of membrane association are regulated, influence each other, and impact overall aPKC asymmetry and activity remain unclear.

Tight spatial and temporal regulation of PAR protein interactions and their membrane localisation are key to the establishment of the aPKC membrane domain in polarising cells. For example, the cell cycle kinases Aurora A and Polo affect the membrane association of the PAR network, coordinating cell polarisation with the cell cycle.^19,41–43^ Moreover, mutual antagonism between PAR proteins is a prevailing mechanism that creates opposing PAR domains in the cell membrane. The kinase Par1 phosphorylates Par3, preventing Par3 oligomerisation and Par3 interaction with aPKC, in this way restricting aPKC from accumulating in the Par1 domain.^44^ Conversely, aPKC phosphorylates Par1, preventing Par1 build-up in the aPKC domain.^45–48^ Recent evidence indicates that aPKC activity, in addition to excluding proteins from opposing domains, is also necessary for restricting itself to its own domain.^21,49^ However, the mechanism by which aPKC controls its own localisation is unknown. Here, we have investigated this problem during polarisation of the *C. elegans* zygote.

The *C. elegans* zygote becomes polarised along the antero-posterior axis in an aPKC dependent manner.^6^ This polarisation drives the zygote’s asymmetric division, leading to anterior somatic and posterior germline precursors. Polarity establishment is triggered by fertilisation, and is characterized by cortical actomyosin flow that carries aPKC, Par6, Par3 and Cdc42 to the anterior cortex (in *C. elegans* these proteins are known as PKC-3, PAR-6, PAR-3 and CDC-42 and collectively named anterior PARs or aPARs).^50,51^ Cortical flow during establishment phase depends primarily on the small GTPase RhoA (RHO-1 in *C. elegans*), whose active GTP-bound state is promoted by the guanine nucleotide exchange factor (GEF), ECT-2, and inhibited by the GTPase-activating proteins (GAPs), RGA-3 and RGA-4.^52–58^ Subsequently, during the polarity maintenance phase, opposing posterior PARs (pPARs, which include the kinase PAR-1, the scaffolding protein PAR-2, LGL-1 and the CDC-42 GAP, CHIN-1) stabilize and maintain the asymmetric localization of aPKC through mutual antagonism between aPARs and pPARs.^47,59–64^ At this later phase, the actomyosin cortex, organised now as an anterior cap, helps refine the aPAR domain^62,65^ and depends on the small GTPase CDC-42 and its corresponding GAP and GEF, CHIN-1 and CGEF-1 respectively.^52,53,64^

In the *C. elegans* zygote, there are two separate membrane-associated pools of PAR-6/aPKC that depend on PAR-3 or CDC-42, respectively.^6,21,31,36,66,67^ . Previously, we proposed a dynamic exchange of PAR-6/aPKC between PAR-3 and CDC-42 (the “aPKC cycle”, Graphical Abstract) in the generation of the aPAR domain.^21^ PAR-6/aPKC segregate most efficiently to the anterior of the zygote when in complex with PAR-3 clusters, which depend on PAR-3 oligomerisation and are mechanically coupled to the actomyosin cortical flow of polarity establishment.^21,41,68–73^ On the other hand, binding to CDC-42 promotes aPKC catalytic activity at the membrane, but the CDC-42/PAR-6/aPKC complex is less efficiently transported by flow.^16,21,22^ Hence a constant cycle of PAR-6/aPKC between PAR-3 and CDC-42 seems necessary in the generation of an asymmetric active domain of aPKC. Previously, we showed that aPKC kinase activity is involved in this cycle and is needed to restrict the size of the aPAR domain to the anterior half of the cell.^21^ When aPKC is inhibited, aPKC and PAR-6 do not segregate to the anterior and remain localised throughout the zygote’s membrane even though PAR-3 still becomes anteriorly localised by the actomyosin flow.^21^ Similar observations have been reported in the *Drosophila* neuroblast, where inactivation of aPKC leads to its expansion towards the basal cell surface.^49^ In the *C. elegans* zygote, we found this aPKC domain expansion to be dependent on CDC-42.^21^ How aPKC mediates its own asymmetry, the identity of the phosphorylation targets, and the mechanisms involved, however remained unknown.

We previously found that kinase-inhibited aPKC remains associated with CDC-42 while uniformly occupying the entire zygote’s membrane.^21^ We therefore questioned if CDC-42 could be a substrate of aPKC and whether CDC-42 phosphorylation could mediate aPKC asymmetry. Here, we report that aPKC activity does lead to phosphorylation of CDC-42 on serine 71. This phosphorylation destabilises PAR-6/aPKC interaction with CDC-42 at the membrane. Once released from CDC-42, PAR-6/aPKC are free to be recruited into PAR-3 membrane clusters, promoting aPKC anterior-segregation by flow. Interestingly, CDC-42 phosphorylation by aPKC not only restricts aPKC localisation to the anterior but also limits aPKC activity, constituting a negative feedback mechanism. We propose that the phosphorylation/dephosphorylation cycle of CDC-42 (turnover of CDC-42 phosphorylation) ensures optimal aPKC distribution and activity levels by tuning the shuttling of PAR-6/aPKC between CDC-42 -active complex- and PAR-3 -anteriorly transported complex-. In addition to fuelling this PAR-6/aPKC cycle, we show that turnover of CDC-42 phosphorylation ensures proper dynamics of the actomyosin cortex by balancing the activities of two CDC-42 effectors, aPKC and MRCK-1. Our results reveal how aPKC can define its own domain of action by controlling the dynamic protein interactions of aPARs and contributing to the organisation of the actomyosin cortex - both through phosphorylation of CDC-42.

## RESULTS

### aPKC-dependent phosphorylation of CDC-42 reduces CDC-42 membrane localisation

To test our hypothesis that aPKC controls its own asymmetric localisation through phosphorylation of CDC-42, we first performed an *in vitro* kinase assay using purified recombinant human aPKC and Cdc42, which showed that aPKC can phosphorylate Cdc42 (**Figure 1A**). Using the group-based prediction tool GPS 5.0^74^, we identified three predicted aPKC phosphorylation sites in CDC-42: T3, S71 and S106. Among these, S71 is evolutionary conserved from worms to humans (**Figure 1B**) and has been reported to regulate Cdc42 function in mammalian tissue culture cells, impacting Cdc42 interaction with downstream effectors.^75–77^ To investigate if S71 could be an aPKC phosphosite, we compared phosphorylation levels of recombinant MBP-tagged CDC-42 vs. MBP-tagged non-phosphorylatable CDC-42 variant (serine 71 to alanine, S71A) in an *in vitro* kinase assay.

**Figure 1.**
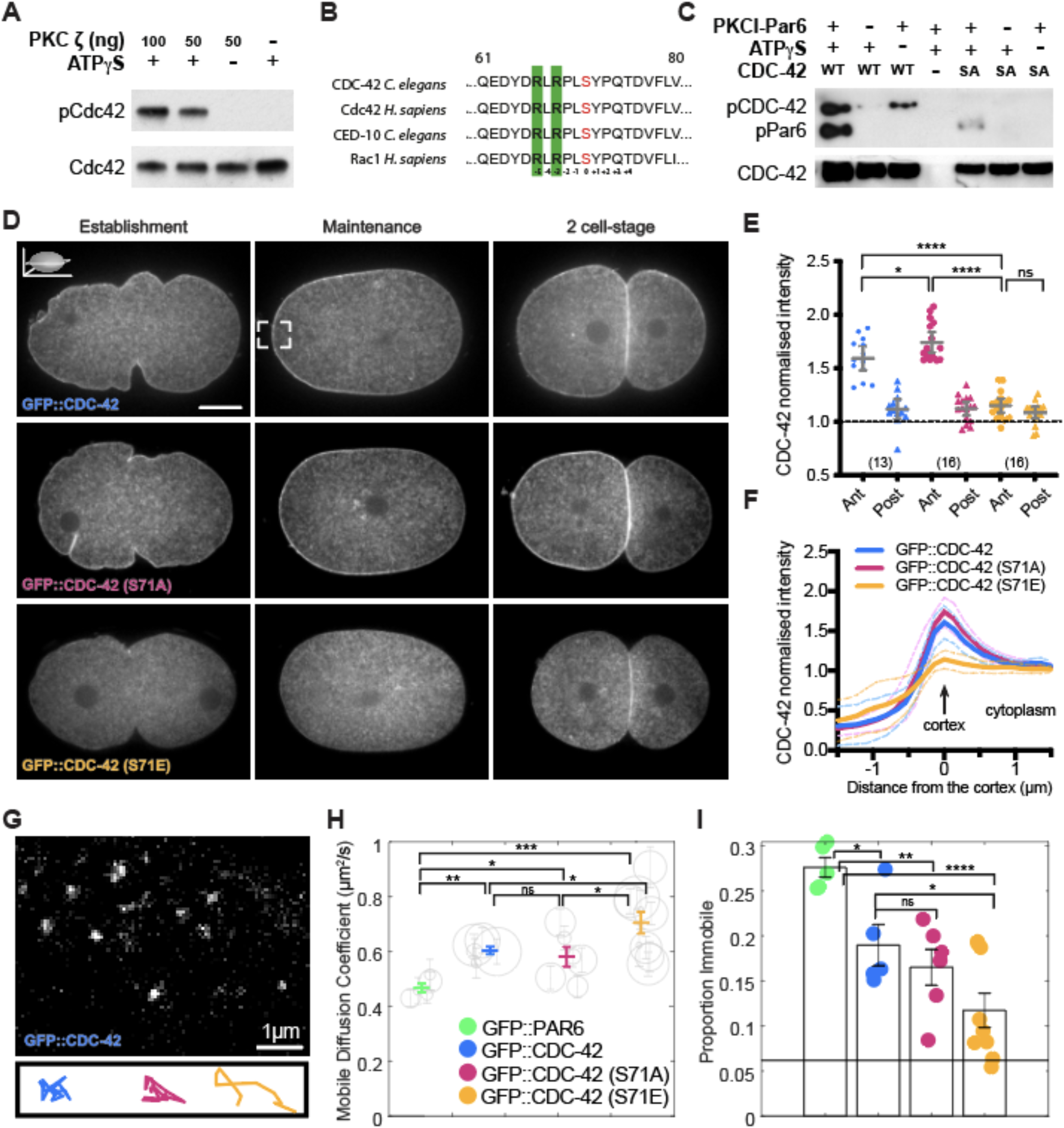
Analysis of CDC-42 phosphomutants indicates that phosphorylation of serine 71 reduces CDC-42 membrane localisation and asymmetry. **A.** Immunoblot showing that human Cdc42 can be phosphorylated by human aPKC (PKC ζ) *in vitro* using purified recombinant proteins incubated with ATP-gamma-S. After kinase reaction, the products were alkylated, and thiophosphorylated Cdc42 (pCdc42) was detected with an anti-thiophosphate ester antibody (see methods). Total Cdc42 detection is shown as loading control. **B.** Protein sequence conservation for Cdc42 and Rac1 (CED-10, *C. elegans* homologue) around serine 71 (red). Conserved arginines, which are part of aPKC consensus site are highlighted (green). **C.** Immunoblot showing the detection of phosphorylated CDC-42 and Par6 (positive control) from an *in vitro* kinase assay containing co-purified human PKCI - Par6 and wild-type *C. elegans* CDC-42 (WT) or CDC-42(S71A) (SA). Thiophosphorylation detection was done as in (A). We observed more phosphorylation of CDC-42 wild type compared to CDC-42(S71A) (after loading correction and background subtraction, i.e. subtraction of signal detected in the corresponding minus ATPγS lane). **D.** Representative midsection confocal images of live embryos at establishment, maintenance and 2-cell stage showing GFP::CDC-42, GFP::CDC-42(S71A) and GFP::CDC-42(S71E). Zygotes are always oriented anterior to the left. In these CDC-42 reporter lines we always deplete endogenous CDC-42 by RNAi targeting its 3’ UTR, in this way preventing possible phenotypic rescue due to endogenous CDC-42. **E.** CDC-42 anterior and posterior cortical intensities (values are normalised by its corresponding cytoplasmic CDC-42 levels) observed in CDC-42, CDC-42(S71A) and CDC-42(S71E) maintenance stage zygotes (graph shows mean ± CI 95%, number of embryos analysed are indicated between brackets). Note that CDC-42(S71E) presents a very weak cortical localisation with no asymmetry (no difference in intensity at anterior vs. posterior). **F.** CDC-42 intensity profiles spanning the anterior membrane of CDC-42, CDC-42(S71A) and CDC-42(S71E) maintenance stage zygotes, showing mean ± SD. Briefly, a 60x60 pixel area from a straightened anterior cortex (see inset in D. in GFP::CDC-42 maintenance) was projected in the y-axis to give a cross section profile spanning the cortex/membrane. The values are normalised so that the cytoplasmic levels are set to 1. **G.** Detection of wild-type CDC-42 particles at the membrane using HILO imaging. Below we show representative tracks obtained by following CDC-42 particles at the membrane in the different CDC-42 variants (blue: CDC-42, pink: CDC-42(S71A), orange: CDC-42(S71E)). **H.** Graph showing the diffusion coefficients (D) of the mobile fraction of CDC-42 variants and PAR-6, obtained from gamma fits to all mobile tracked data. Errors represent 95% confidence intervals on the fits. Diffusion coefficients of gamma fits for mobile data obtained from individual maintenance stage zygotes is shown by the position of the grey circles (circle size represents the number of tracks analysed per embryo and the lines represent 95% confidence intervals on the fits). **I.** Proportion of tracks found in the immobile fraction (D < 0.1 μm^2^/s) in each condition (graph shows all data points and mean ± SEM). Horizontal black line indicates the limit of detection of immobile fraction (baseline, See Figure S3F). Average number of tracks (± SEM) for the embryos analysed in H and I, CDC-42: 3295 ± 57 tracks/embryo in 5 embryos; CDC-42(S71A): 3047 ± 62 in 6 embryos; CDC-42(S71E): 3532 ± 47 in 9 embryos and PAR-6: 1380 ± 790 in 4 embryos. Unpaired, two-tail Student’s T test *p<0.05, **p<0.01, ***p<0.001, ****p<0.0001, ns not significant. Scale bar in establishment zygote: 10 µm. See also Figure S1A-B, S2 and S3.

Here we used human aPKC co-purified with Par6, which supports aPKC interaction with Cdc42, and confirmed that aPKC can phosphorylate wild-type CDC-42 *in vitro* (**Figure 1A and 1C**). Phosphorylation of CDC-42 strongly decreased when we mutated serine 71 to alanine (S71A) (**Figure 1C**), suggesting that this is the primary site on which aPKC phosphorylates CDC-42. We also detected the previously described aPKC phosphorylation of Par6 (**Figure 1C**).^78^ Phosphorylation of Par6 did not occur in the absence of CDC-42 (**Figure 1C**, middle lane), supporting the known role of CDC-42 in promoting aPKC kinase activity.^21–23,25,26^ Interestingly, phosphorylation of Par6 was also reduced in the presence of CDC-42(S71A) compared to that observed in the presence of wild-type CDC-42 (**Figure 1C**). Thus, the S71A point mutation could affect the ability of CDC-42 to activate aPKC, or alternatively, non-phosphorylatable CDC-42(S71A) could act as a competitive inhibitor at the concentrations used in this *in vitro* assay, blocking aPKC activity towards other sites.

Together, these results show that aPKC can phosphorylate CDC-42 *in vitro* and that a S71A mutation prevents CDC-42 phosphorylation by aPKC.

We next analysed the function of CDC-42 phosphorylation *in vivo*. For this, we constructed transgenic strains expressing GFP::CDC-42, GFP::CDC-42(S71A) (non-phosphorylatable) and GFP::CDC-42(S71E) (phosphomimetic). Wild-type CDC-42 is localised throughout the membrane of the zygote, but enriched at anterior versus posterior once polarity is established (maintenance stage) (**Figure 1D and 1E**). This led to a slight enrichment of CDC-42 at the anterior daughter cell after cell division (**Figure 1D**). The non-phosphorylatable (S71A) variant presented an asymmetry similar to wild-type CDC-42 (**Figure 1D**), but with slightly increased anterior membrane levels (**Figure 1E and 1F**). On the other hand, the phosphomimetic mutant’s membrane localisation was reduced throughout the zygote’s AP axis and did not exhibit anterior membrane enrichment (**Figure 1D-1F**). We also observed that CDC-42(S71E) causes strong embryonic lethality, unlike control or CDC-42(S71A) (**Figure S1A**). The weak membrane localisation observed for CDC-42(S71E) is not due to low expression levels, given that we observed similar CDC-42 protein levels in all three transgenic strains (**Figure S1B**). Therefore, we conclude that the membrane localisation of CDC-42(S71E) variant is reduced and has lost its asymmetry.

Given that the membrane localisation of CDC-42 might be stabilised by CDC-42 interaction with aPARs^22,64,72^, we investigated CDC-42 membrane localisation upon depletion of PAR-6 (**Figure S2**). PAR-6 partial depletion led to a strong reduction in wild-type CDC-42 membrane localisation (**Figure S2D-S2F**) just like that observed for the CDC-42(S71E) variant (**Figure S2A-S2C**), supporting a role for PAR-6 in stabilizing CDC-42 at the plasma membrane. In the CDC-42(S71A) mutant, a partial knockdown of PAR-6 led to loss of CDC-42 asymmetry but some CDC-42 still remained at the membrane (**Figure S2D-S2F**).

Together, these data show that PAR-6 is necessary for CDC-42 membrane localisation and asymmetry, and suggest that phosphorylation of CDC-42 might reduce CDC-42 membrane localization by disrupting its association with PAR-6.

We next analysed the impact of CDC-42 phosphorylation on CDC-42 dynamics at the membrane. Tracking the movement of individual molecules of CDC-42 in the plane of the membrane using near-TIRF microscopy (HILO^79^), we found that, in the zygote, CDC-42 diffuses more quickly than aPARs, consistent with previous work (**Figure 1G-1I**).^68,73,80–82^ Because of the slower dynamics of PAR-6 at the membrane compared to CDC-42 (**Figure 1G-1I**), we reasoned that if CDC-42 phosphorylation were to reduce its interaction with PAR-6, then this phosphorylation should lead to an increase in diffusivity of CDC-42 at the membrane. We therefore analysed the diffusion of CDC-42 variants at the membrane. In all variants the distribution of GFP::CDC-42 diffusion coefficients (D) appeared bimodal (**Figure S3A-S3C)** and was well fit (R^2^>0.95) by a model comprising two mobility states: mobile (D> 0.1 µm^2^/s) and immobile (D< 0.1 µm^2^/s). We further confirmed these two states in our CDC-42 diffusion data by using nonparametric Bayesian statistics (SMAUG^83^) **(Figure S3E**).

Interestingly, the mobile pool of CDC-42(S71E) diffused significantly faster than the mobile pools of wild-type CDC-42 or CDC-42(S71A) **(Figure 1H)**. Moreover, the proportion of immobile CDC-42(S71E) was reduced to levels close to the minimum levels that are possible based on simulations, i.e. CDC-42(S71E) presented near total reduction in immobile CDC-42 (**Figure 1I** and **Figure S3F**). Note that we detected a similar number of CDC-42 molecules across strains (see legend **Figure 1I**), indicating that the overall recruitment of CDC-42(S71E) to the membrane is not reduced; rather, its overall reduced levels at the membrane (**Figure 1D-1F**) might be explained by reduced stability at the membrane. We also tracked PAR-6 dynamics under the same imaging conditions (**Figure 1H, 1I** and **Figure S3D**), highlighting that CDC-42 wild type and CDC-42(S71A) presented dynamics closer to those of PAR-6 than CDC-42(S71E) did. The faster dynamics of the phosphomimetic mutant are consistent with a reduced interaction of this CDC-42 variant with the slower-diffusing PAR-6.

In summary, the CDC-42 phosphomutant analyses indicate that phosphorylation of CDC-42 leads to destabilisation of CDC-42 at the membrane and to loss of CDC-42 asymmetry, possibly through disruption of CDC-42/aPAR interactions.

### Phosphorylation of CDC-42 at serine 71 promotes the dissociation of PAR-6/aPKC from CDC-42

We sought to directly test if phosphorylation of S71 affected the interaction between CDC-42 and PAR-6/aPKC. Although PAR-6/aPKC can bind to GTP-locked mutants of CDC-42 *in vitro* and in co-immunoprecipitation experiments, recent evidence suggests that native CDC-42/PAR-6 complexes are found at the plasma membrane and are disrupted by cell lysis in detergent.^36^ Therefore, we tested CDC-42 association with aPKC using two independent approaches that aimed to preserve the integrity of the membrane as much as possible. First, we artificially docked wild-type CDC-42, CDC-42(S71A) or CDC-42(S71E) to the membrane of polarised zygotes and measured their ability to recruit aPKC. Each GFP::CDC-42 variant was driven to the membrane at similar levels by expressing a membrane-tethered nanobody that recognises GFP (PH-GBP^21^) (**Figure 2A, 2B** and **Figure S1C and S1D**). This eliminates the differences in CDC-42 membrane levels between the different CDC-42 variants (**Figure S1D-S1E**), so that differences in aPKC membrane levels can be used to asses aPKC recruitment by the CDC-42 variants in an unbiased way. We observed that aPKC levels at the membrane were greater following membrane-docking of CDC-42(S71A) compared to the docking of wild-type, whilst membrane-docked CDC-42(S71E) presented reduced aPKC membrane levels compared to docked wild-type CDC-42 (**Figure 2C and 2D and Figure S1E**). We can conclude that the S71A mutation increases, while the S71E mutation decreases, CDC-42’s ability to recruit aPKC. Note that aPKC is still recruited mostly at the anterior, although membrane-docked CDC-42 reporters were observed throughout the zygotes’ membrane (**Figure S1C**), indicating that a factor at the anterior of the zygote promotes the cortical recruitment of aPKC by CDC-42 and/or a factor at posterior reduces it. Overall, these results indicate that phosphorylation of CDC-42 on S71 disrupts CDC-42/aPKC association.

**Figure 2.**
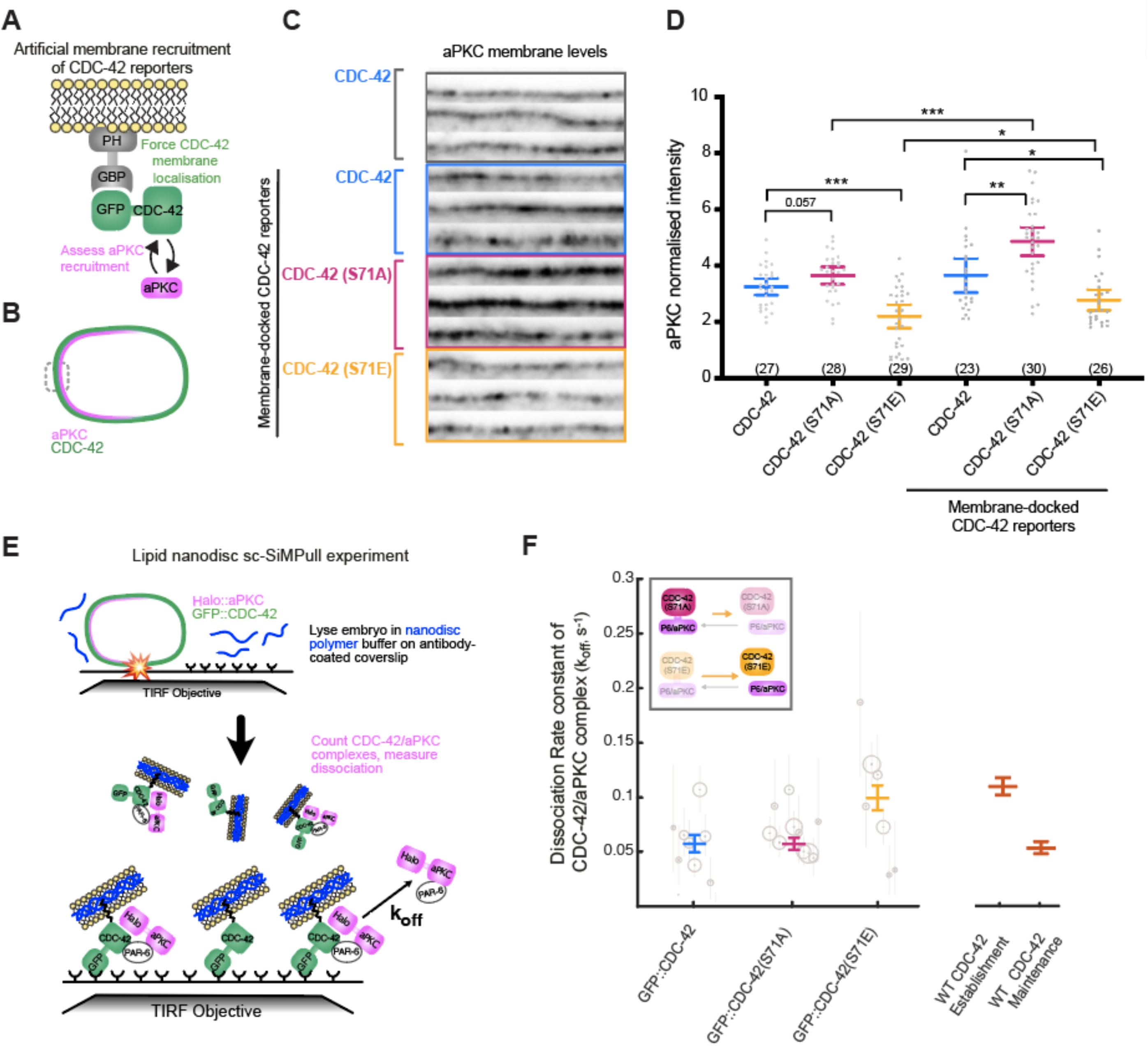
CDC-42 phosphorylation on serine 71 prevents CDC-42-dependent cortical recruitment of aPKC and promotes CDC-42/aPKC dissociation. **A.** *In vivo* approach to artificially recruit CDC-42 reporters at the membrane (membrane-docked) using a membrane-tethered nanobody against GFP (PH-GBP). **B.** Zygotes with membrane-docked CDC-42 were immunostained for aPKC to determine aPKC membrane levels at the anterior domain (anterior-most selected region). **C.** Representative anterior membrane regions (three per condition) showing aPKC levels in the non-membrane docked GFP::CDC-42 reporter and in the membrane-docked GFP::CDC-42, GFP::CDC-42(S71A) and GFP::CDC-42(S71E) reporters (endogenous CDC-42 is depleted in all CDC-42 reporters). **D.** Average aPKC intensity values at the anterior membrane (measured in a 60x60 pixel area from a straightened anterior-most cortex) of zygotes during polarity maintenance. Anterior membrane intensity of aPKC for each zygote is normalised with its corresponding aPKC cytoplasmic level (graph shows all data points and mean ± CI 95%). Unpaired, two-tail Student’s T test *p<0.05, **p<0.01, ***<0.001. See also Figure S1C-E. **E.** Schema of the lipid nanodisc single-cell pull-down (sc-SiMPull) experiment. Cells expressing GFP::CDC-42 variants and carrying endogenously tagged HaloTag::aPKC are lysed using a pulsed infrared laser in the presence of amphipathic polymers to form native lipid nanodiscs. sc-SiMPull was used to capture GFP::CDC-42 variants, which were detected using single-molecule TIRF microscopy. Complexes containing HaloTag::aPKC were identified and dissociation rate constants (k_off_) of those complexes were measured (see Methods and Figure S4). **F.** Dissociation rate constants measured for CDC-42/aPKC, CDC-42(S71A)/aPKC and CDC-42(S71E)/aPKC complexes extracted from mixed-stage single *C. elegans* zygotes. Gray circles represent individual experiments, with the size of the circle indicating the number of CDC-42/aPKC complexes captured from each zygote and the error bars representing the Bayesian 95% credible interval for the estimated k_off_. Coloured bars show the maximum probability estimate and Bayesian 95% credible intervals for the k_off_ obtained by pooling all single-molecule measurements. Note that statistical confidence of these data is assessed by the width of the 95% credible interval, and differences between conditions are considered significant when the 95% credible intervals (coloured bars) do not overlap. Total number of complexes analysed: 855 CDC-42/aPKC complexes from 9 embryos; 1,579 CDC-42(S71A)/aPKC complexes from 10 embryos; 733 CDC-42(S71E)/aPKC complexes from 7 embryos. Orange bars are measurements of k_off_ for the complex of wild-type mNG::CDC-42 and PAR-6 during polarity establishment or polarity maintenance (Deutz et al., 2023), shown here for comparison. See also Figure S4.

Next, we directly measured the interaction between CDC-42 and aPKC using single-cell, single-molecule pull-down experiments (sc-SiMPull^35,36^). We used a pulsed infrared laser to instantaneously lyse single zygotes in the presence of an amphipathic polymer (diisobutylene maleic acid, DIBMA) that forms lipid nanodiscs from the native cellular membrane. Then, we captured GFP::CDC-42 molecules using anti-GFP nanobodies immobilized on a glass coverslip, identified those associated with labelled HaloTag::aPKC, and measured the dissociation rate constant (k_off_) of the CDC-42/aPKC complex (**Figure 2E**). We readily detected complexes containing GFP::CDC-42 variants and endogenous HaloTag::aPKC and observed that mutations of S71 altered the stability of these complexes: the S71E variant exhibited a k_off_ of 0.099 ± 0.012 s^-1^, which was approximately 2-fold faster than the S71A variant (k_off_ = 0.057 ± 0.006 s^-1^) or wild-type GFP::CDC-42 (k_off_ = 0.057 ± 0.008 s^-1^) (**Figure 2F and Figure S4**). These data demonstrate that aPKC interacts more stably with CDC-42(S71A) than with CDC-42(S71E), further suggesting that phosphorylation destabilises CDC-42/aPKC association.

We recently reported that CDC-42/PAR-6 complexes are more stable during polarity maintenance, when CDC-42 is strongly required for normal PAR-6/aPKC localisation (**Figure 2F**, orange bars).^36^ Remarkably, the CDC-42/aPKC k_off_ values that we measured for the S71A and S71E variants were very similar to the wild-type CDC-42/PAR-6 k_off_ values during maintenance and establishment, respectively (**Figure 2F**). Note that the PAR-6 and aPKC dissociate from each other approximately 50-fold slower than either protein dissociates from CDC-42;^35^ thus, the k_off_ values that we are comparing here most likely reflect dissociation of PAR-6/aPKC heterodimers from CDC-42. Together, these results suggest that phosphorylation of CDC-42 by aPKC may account for the differential stability of CDC-42/PAR-6/aPKC complexes during polarity establishment compared to polarity maintenance. We propose that in establishment phase, CDC-42 phosphorylation may promote the turnover of the CDC-42/PAR-6/aPKC complex, whereas in maintenance, the CDC-42/PAR-6/aPKC complex may be stabilised by a decrease in CDC-42 phosphorylation. This model is consistent with previous work showing that the PAR-3/PAR-6/aPKC complex is more abundant in establishment while the CDC-42/PAR-6/aPKC complex is favoured during maintenance.^21,31,36,41,67,72^

### Phosphorylation of CDC-42 at serine 71 restricts aPKC to the anterior membrane domain

Thus far our data suggests that phosphorylation of CDC-42 on S71 weakens the interaction between CDC-42 and PAR-6/aPKC. Previously we hypothesised that aPKC release from CDC-42 would lead to aPKC recruitment by PAR-3 clusters that segregate with actomyosin flow, thereby promoting aPKC asymmetry^21^. Thus, we next analysed if CDC-42 phosphorylation regulates the asymmetric localisation of aPKC, by analysing the localisation of PAR-3 and aPKC in the CDC-42 phosphomutants.

In mid-plane images of the zygote focusing at the boundary of the aPAR membrane domain, we consistently observed that aPKC localization extends more towards the posterior compared to PAR-3, and this posterior aPKC expansion is CDC-42 dependent (**Figure 3A**, grey double head arrow and **Figure 3B and 3C**). Hence, this aPKC extension can be used as a measure of the abundance of CDC-42/PAR-6/aPKC complex at the membrane.^21^ We observed that in the CDC-42(S71E) mutant, aPKC expansion was absent and aPKC localisation matched that of PAR-3 (**Figure 3B, 3C and Figure S5G**). Cortical sections of CDC-42(S71E) zygotes also showed increased co-localisation of aPKC with PAR-3 and enhanced aPKC asymmetry (ASI index) (**Figure 3D-3F**). Moreover, in CDC-42(S71E) zygotes, aPKC presented an increased clustered organisation during maintenance phase (analysed by the coefficient of variation, CV) (**Figure 3G**), more typical of PAR-6/aPKC during establishment phase when co-localising with PAR-3.^21,41^ These results indicate that membrane localisation of PAR-6/aPKC in the CDC-42(S71E) mutant depends primarily on PAR-3. We observed a similar dependency when CDC-42 is depleted by RNAi (**Figure 3E-3G**).

**Figure 3.**
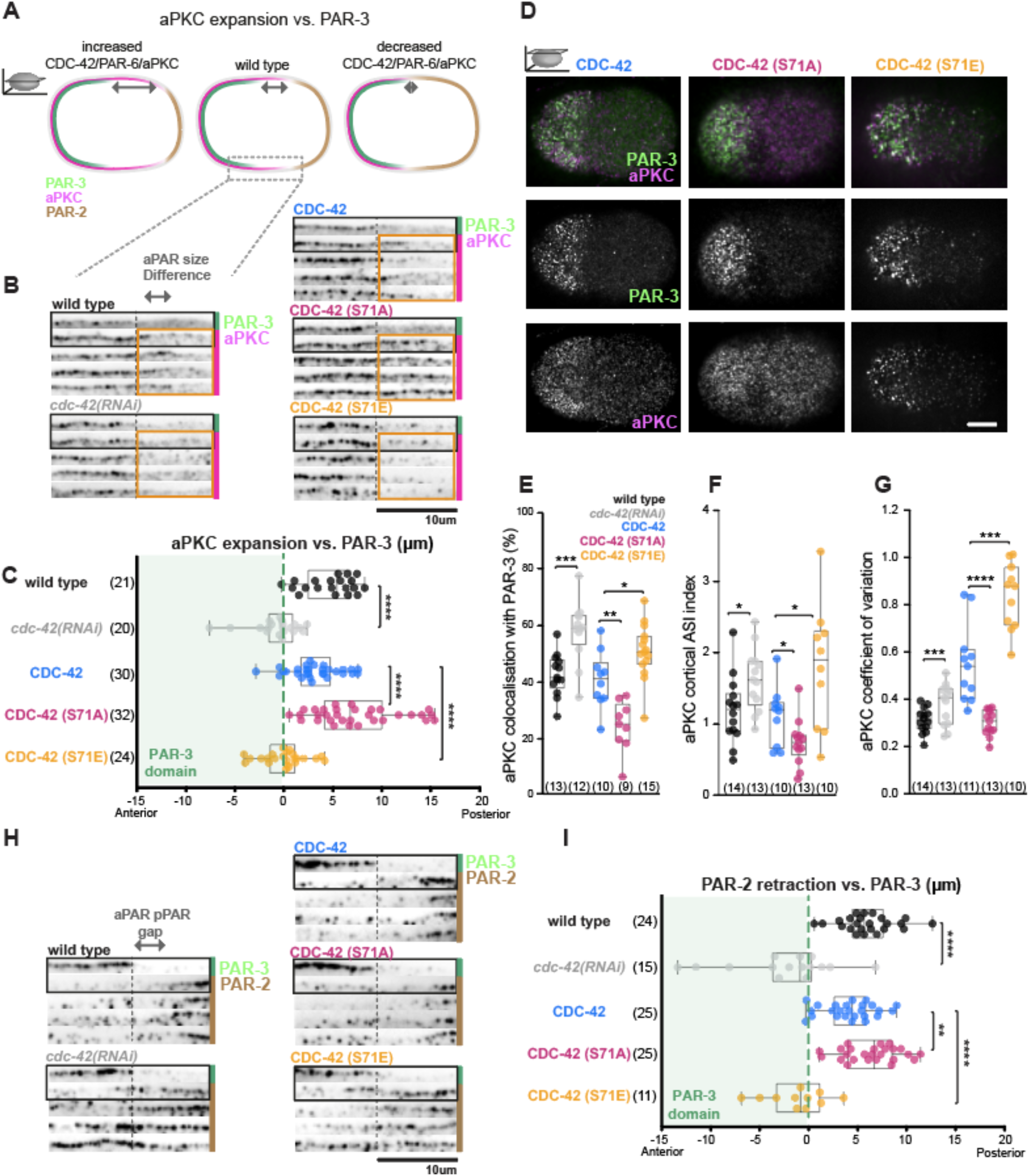
Preventing CDC-42 phosphorylation at serine 71 leads to aPKC cortical expansion and posterior PAR domain reduction. **A.** Schema showing that aPKC expansion vs. PAR-3 can be used as a readout of the abundance of the CDC-42/PAR-6/aPKC complex at the membrane. **B.** Representative flattened-out membranes from PAR-3 and aPKC co-immunostained zygotes, showing the difference between PAR-3 and aPKC domain size in wild type, *cdc-42* depleted (RNAi of its 3’ UTR), GFP::CDC-42, GFP::CDC-42(S71A) and GFP::CDC-42(S71E) strains (endogenous CDC-42 is depleted in all CDC-42 reporters). Cortical sections at the end of the aPAR domain are shown for each condition (four zygotes per condition). PAR-3 and aPKC are shown for the first zygote in each condition. For the other zygotes only aPKC is shown but its position is relative to the end of its corresponding PAR-3 domain (dashed vertical line). Orange squares highlight the region, expanding 10µm from the posterior boundary of the PAR-3 domain, where we see differences in aPKC expansion relative to PAR-3 in the different conditions. **C.** Quantification of aPKC expansion vs. PAR-3 at polarity maintenance. S71A mutation in CDC-42 promotes the CDC-42/PAR-6/aPKC membrane domain (as shown by the larger expansion of aPKC relative to PAR-3), whereas S71E mutation decreases membrane CDC-42/PAR-6/aPKC and aPKC localisation matches that of PAR-3. **D-G.** Representative cortical confocal images of PAR-3 and aPKC in the indicated CDC-42 strains (**D**). In cortical sections we have analysed aPKC and PAR-3 co-localisation (**E**), aPKC asymmetry using ASI index (normalised against wild-type asymmetry, “1” denotes wild-type asymmetry and “0” no asymmetry, see methods) (**F**) and aPKC coefficient of variation as a means to measure clustered (closer to 1) vs. diffusive state (closer to 0) (**G**). **H.** Representative flattened-out membranes from PAR-3 and PAR-2 co-immunostained zygotes at the PAR-2 / PAR-3 domain boundary in wild type, *cdc-42* depleted (RNAi of its 3’ UTR), GFP::CDC-42, GFP::CDC-42(S71A) and GFP::CDC-42(S71E) strains (endogenous CDC-42 is depleted in all CDC-42 reporters). Four cortices are shown for each condition. PAR-3 and PAR-2 are shown for the first zygote in each condition. For the other zygotes only PAR-2 is shown but its position is relative to the end of its corresponding PAR-3 domain (dashed vertical line). **I.** Quantification of PAR-2 retraction vs. PAR-3 at polarity maintenance. *cdc-42* depleted and S71E mutation in CDC-42 strongly reduce PAR-2 retraction with many zygotes presenting overlapping PAR-3 and PAR-2 domains. Zygotes with full PAR-3 and PAR-2 overlap (27% in *cdc-42* depleted and 45% in CDC-42(S71E) zygotes) have been removed for ease of comparison between graphs C and I. All box plots show the median ± IQR and all data points. Unpaired, two-tail Student’s T test *p<0.05 **p<0.01, ***p<0.001, ****p<0.0001. Scale bar: 10 µm. See also Figure S5.

Conversely, in the CDC-42(S71A) mutant, aPKC expansion relative to PAR-3 was larger compared to that observed in wild type (**Figure 3B and 3C and Figure S5G**), indicating a stabilisation of the CDC-42/PAR-6/aPKC complex at the membrane (**Figure 3A**). In cortical sections we also observed that aPKC displayed a more homogenous and less clustered distribution, more typical of CDC-42^21,72^, and aPKC co-localised less with PAR-3 compared to the co-localisation observed in wild-type CDC-42 or CDC-42(S71E) (**Figure 3D-3G**). aPKC expansion in CDC-42(S71A) was not only detected in immunostained zygotes but also in live zygotes, becoming apparent at establishment stage but stronger by maintenance (**Figure S5A, S5B, S5D**). Since CDC-42(S71A) membrane levels at the posterior are similar to wild-type CDC-42 (**Figure 1E**), aPKC expansion towards the posterior membrane most likely reflects the more stable interaction observed for aPKC with CDC-42(S71A) than with wild-type CDC-42 (**Figure 2D and 2F**).

In summary, these results show that the disruption of the CDC-42/PAR-6/aPKC complex by phosphorylation of CDC-42 on S71 leads to increased co-localisation of PAR-3 and aPKC and enhances aPKC asymmetry. Conversely, stabilisation of the CDC-42 complex by preventing S71 phosphorylation decreases aPKC co-localisation with PAR-3 and promotes aPKC expansion into the posterior. Overall, our findings suggest that turnover of CDC-42 phosphorylation can mediate the proposed shuttling of PAR-6/aPKC between CDC-42 and PAR-3 clusters (aPKC cycle^21^), controlling aPKC asymmetry in the polarisation of the zygote. aPKC-mediated phosphorylation of CDC-42 constitutes a self-regulatory mechanism, whereby aPKC promotes its own asymmetric localisation.

### Phosphorylation of CDC-42 at serine 71 limits aPKC activity and regulates the aPAR/pPAR boundary

Previously, we have found that the ability of aPKC to exclude the posterior protein, PAR-2, from the anterior cortex of the zygote is promoted by CDC-42.^21^ Hence, we wanted to investigate if CDC-42 phosphorylation could impact aPKC activity. To assess aPKC activity *in vivo,* we analysed PAR-2 posterior localisation.^21,47^ In wild-type zygotes co-immunostained for PAR-2 and PAR-3, we observed a gap between the PAR-3 and PAR-2 domains (**Figure 3H, 3I and Figure S5H**), whose size matched the ∼5µm expansion of aPKC relative to PAR-3 (**Figure 3C**). Upon CDC-42 depletion by RNAi, this PAR-3/PAR-2 gap was no longer present and in many zygotes the PAR-2 and PAR-3 domains overlapped, with some zygotes (27%) showing almost complete overlap. This agrees with our *in vitro* kinase assay (**Figure 1C**, lane 4) and our previous observations indicating that CDC-42 promotes aPKC activity.^21^ In CDC-42(S71E) mutant zygotes, which present a perturbed CDC-42/aPKC association (**Figure 2D, 2F**), we observed a very similar phenotype to *cdc-42(RNAi)* (**Figure 3H, 3I and Figure S5H**), suggesting that a stable association between CDC-42/aPKC is required for aPKC activity. Conversely, the CDC-42(S71A) variant showed a larger gap between PAR-3 and PAR-2 domains (**Figure 3H, 3I and Figure S5H**), which matched the larger aPKC expansion relative to PAR-3 observed in this strain (**Figure 3C**). These results indicate that CDC-42 phosphorylation can regulate aPKC activity, at least in part by tuning the abundance of the active CDC-42/aPKC complex. Together, our findings suggest the existence of a negative feedback loop whereby CDC-42 promotes aPKC activity, but aPKC-mediated phosphorylation of CDC-42 limits the stability and/or abundance of active CDC-42/aPKC complexes.

### Phosphorylated CDC-42 is found in anterior cortical foci that depend on aPKC activity and actomyosin organisation

Thus far our data suggest a mechanism by which turnover of CDC-42 phosphorylation tunes aPKC association with CDC-42 or PAR-3 and, as a result, controls aPKC activity and asymmetry. Besides controlling aPKC, CDC-42 also has well known roles regulating the actin cytoskeleton. ^84,85^ Hence, CDC-42 phosphorylation could impact aPKC asymmetry, not only through the turnover of the CDC-42/PAR-6/aPKC complex, but also by regulating the actomyosin cortex. Interestingly, staining with a phospho-specific antibody against CDC-42 phospho-S71 (CDC-42 pS71) revealed an enrichment of phosphorylated CDC-42 at cortical foci in the anterior half of wild-type zygotes. These foci closely resemble the actomyosin foci that underlie the membrane during polarity establishment (**Figure 4A and 4C**). Upon RNAi knockdown of CDC-42 or mutation of CDC-42 to a non-phosphorylatable form (S71A), we did not detect staining of these cortical foci (**Figure 4A and 4B**), supporting the specificity of CDC-42 pS71 detection. Furthermore, knock-down of *C. elegans* homologues of Rac1, which can also be phosphorylated at this conserved site and recognised by the pS71 antibody,^75,76,86^ did not lead to reduction in pS71-foci staining (**Figure S6A and S6B**). We did not observe cortical foci of total CDC-42 when imaging the GFP::CDC-42 transgene (**Figure S6C)**. This could indicate that only a minority of CDC-42 is in this phosphorylated and focally-localised state. We also did not observe the phosphomimetic reporter of CDC-42 (GFP::CDC-42(S71E)) in foci (**Figure S6C)**, suggesting that the phosphomimetic reporter does not mimic perfectly the phosphorylated state of CDC-42. Accumulation of CDC-42 pS71 at such foci might require specific recognition of the phosphate moiety and/or prior stabilisation of non-phosphorylated CDC-42 at the membrane, which is impeded in the phosphomimetic mutant (**Figure 1D-1F**).

**Figure 4.**
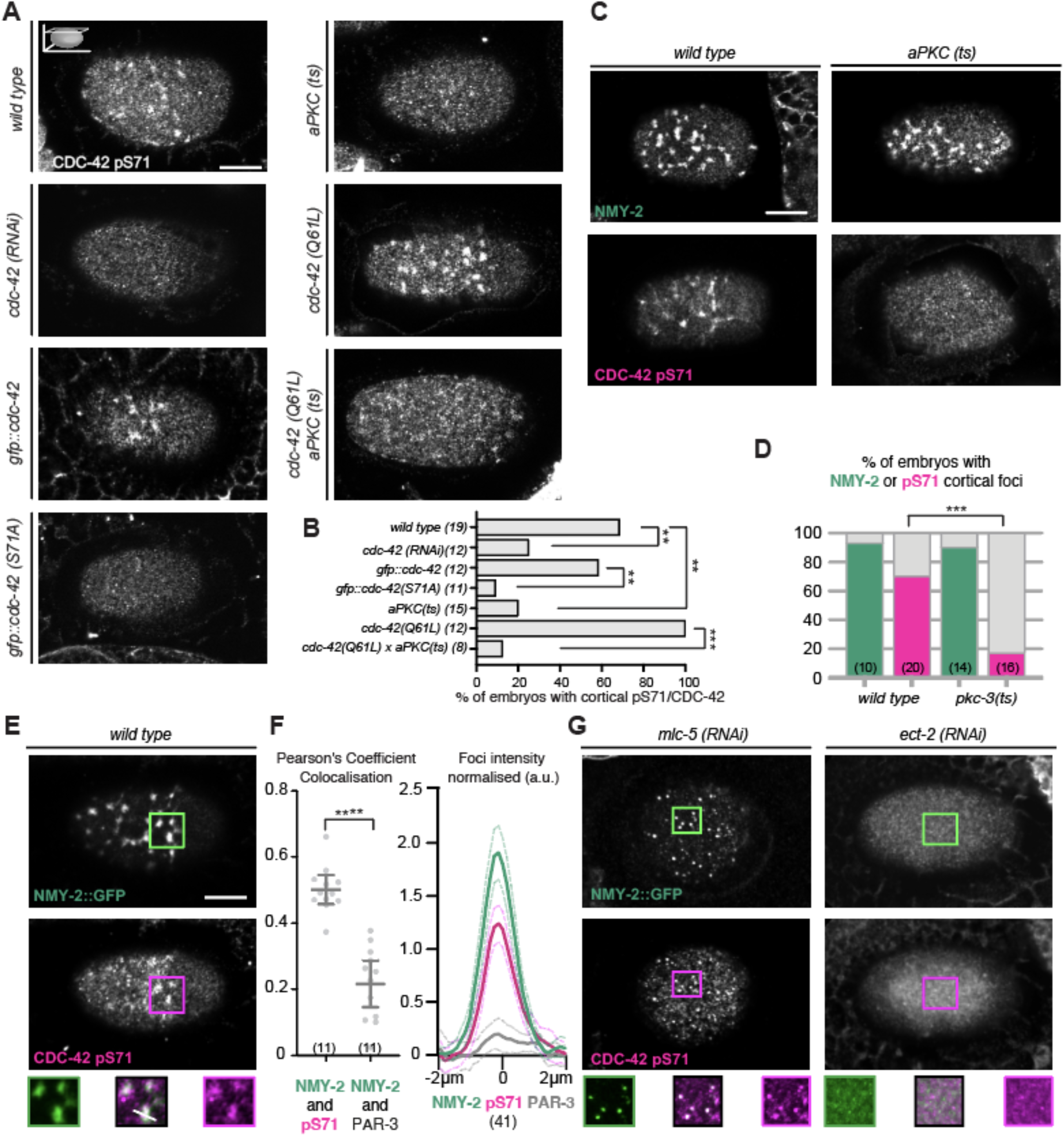
aPKC activity promotes anterior cortical foci of CDC-42 pS71 that co-localise with and depend on myosin foci. **A.** Confocal images showing the immunofluorescent detection of CDC-42 phosphorylation on serine 71 at the cortex of polarity establishment zygotes, revealing the presence or absence of cortical foci in the conditions tested. **B.** Quantification showing the percentage of zygotes with CDC-42 pS71 cortical foci inferred from 2D intensity correlation analysis (see methods). CDC-42 pS71 cortical foci are absent from the cortex upon mutation of the CDC-42 S71 phosphorylation site to alanine (endogenous CDC-42 is depleted in this strain) and are strongly reduced upon inhibition of aPKC kinase activity. **C.** Confocal images of wild-type and *aPKC(ts)* zygotes at polarity establishment, immunostained for NMY-2 and CDC-42 pS71. **D.** Quantification of NMY-2 and CDC-42 pS71 foci presence in wild-type and *aPKC(ts)* establishment zygotes. Although *aPKC(ts)* embryos show clear cortical NMY-2 foci we observed a clear reduction in CDC-42 pS71 foci compared to wild type. **E-G.** Representative cortical confocal images of polarity establishment wild-type zygote and zygotes where the actomyosin cytoskeleton is perturbed (*mlc-5* or *ect-2 RNAi*), co-labelled for NMY-2 and CDC-42 pS71. Insets highlight the partial co-localisation of CDC-42 pS71 (magenta) with NMY-2 (green) in the indicated areas. **F.** Quantification of NMY-2 and CDC-42 pS71 foci colocalization (Pearson’s Coefficient, 1 would indicate perfect co-localisation) and plot profiles of NMY-2 and CDC-42 pS71 intensity across foci (mean ± SEM, example of ROI analysed in middle inset in E -white line-). PAR-3 used as negative control. Chi-square test **p<0.01, ***p<0.001 in B and D. Unpaired, two-tail Student’s T test ****p<0.0001 in F. Scale bar: 10 µm. See also Figure S6.

To address if CDC-42 phosphorylation depends on aPKC *in vivo*, we examined CDC-42 pS71 foci upon aPKC knockdown (*aPKC(RNAi)*) or in a temperature sensitive kinase-inactive aPKC mutant (*pkc-3 (ts)).*^21^ Under these conditions, CDC-42 pS71 foci were strongly reduced (**Figure 4A-4D and Figure S6A and S6B**), indicating that CDC-42 phosphorylation depends on aPKC activity in the zygote. Furthermore, CDC-42/GTP has been proposed to promote aPKC activity^21–23,25^ and, in agreement with this, we found that the constitutively active mutant of CDC-42, CDC-42(Q61L), which favours its GTP state, exhibited more prominent CDC-42 pS71 foci that are dependent on aPKC activity (as seen in *cdc-42(Q61L)* and in *cdc-42(Q61L); aPKC(ts)* strains, **Figure 4A and 4B**). In other systems, Akt kinase can phosphorylate small GTPases in this conserved site.^86^ However, depletion of the *C. elegans* homologues (AKT-1 and AKT-2), did not lead to a reduction in the pS71-foci staining in the zygote (**Figure S6A and S6B**). Overall, these data show that aPKC promotes CDC-42 phosphorylation on S71 and that, in the *C.elegans* zygote, this phosphorylation is enriched in anterior cortical foci during polarity establishment. Given that we did not observe an enrichment of aPKC at these foci (**Figure S6D and S6E**), consistent with the destabilisation of CDC-42/PAR-6/aPKC complex by CDC-42 phosphorylation (**Figure 2**), we suggest that aPKC phosphorylation of CDC-42 could be promoted at these cortical foci or alternatively CDC-42 pS71 could be somehow stabilised at these foci.

Due to their similar appearance, we investigated if CDC-42 pS71 foci depend on the presence of non-muscle myosin II (NMY-2) foci. Cortical foci of NMY-2::GFP and immunostained CDC-42 pS71 partially co-localised in the zygote (**Figure 4E and 4F**). PAR-3 was used as a negative control as it is also present at the anterior domain but not enriched at cortical actomyosin foci.^68^ NMY-2 and CDC-42 pS71 foci were observed during polarity establishment and strongly reduced during polarity maintenance (**Figure S6F**). Next, we perturbed the actomyosin cortex, and we observed resulting changes in localisation of CDC-42 pS71. First, depletion of the myosin essential light chain, MLC-5, led to the condensation of NMY-2 foci into tight immobile puncta that did not disassemble (**Figure 4G**).^87^ CDC-42 pS71 staining in *mlc-5(RNAi)* zygotes co-localised with these NMY-2 puncta (**Figure 4G**).

Second, down-regulation of the RHO pathway through partial depletion of RHO-1 or depletion of the Rho guanine nucleotide exchange factor ECT-2 led to the disappearance of NMY-2 foci^52,53^, and CDC-42 pS71 foci also disappeared (**Figure 4G, Figure S6G**).

Overall our results indicate that CDC-42 pS71 enrichment at cortical foci depends on aPKC activity and on NMY-2 foci. Importantly, the well-defined NMY-2 foci observed in *aPKC(ts)* and in CDC-42(S71A), where CDC-42 pS71 foci are gone (**Figure 4C and 4D and Figure 5F and 5G**), indicates that in these strains the loss of CDC-42 pS71 staining is not an indirect consequence of actomyosin disruption; but rather, reflects CDC-42 pS71 foci dependency on aPKC phosphorylation of CDC-42.

**Figure 5.**
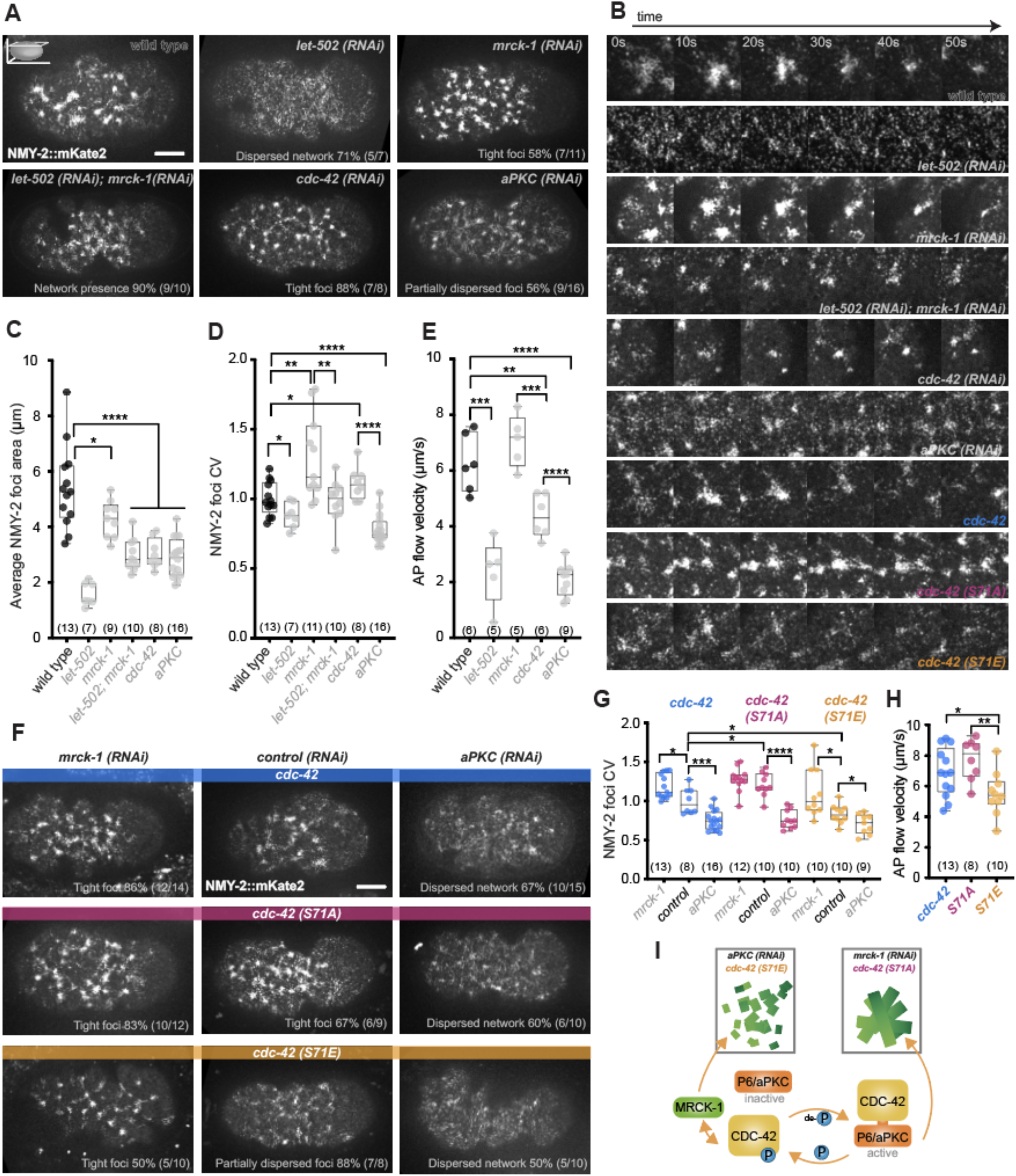
Phosphorylation of CDC-42 balances aPKC and MRCK-1 function in actomyosin foci organisation during polarity establishment. **A.** Representative cortical confocal live images of NMY-2 in polarity establishment wild-type, *let-502*, *mrck-1*, double *let-502/mrck-1* (1:1 RNAi bacteria mix, see methods for more details), *cdc-42* and *aPKC (RNAi)* depleted zygotes. Penetrance of phenotypes observed (qualitative analyses) indicated on images. **B.** Time-lapse images following the evolution of representative myosin foci for the different conditions tested. **C.** Average NMY-2 foci area plotted for the different conditions studied. **D.** Graph showing the normalised coefficient of variation (CV) of NMY-2 intensity (average of wild type set to 1) as a measurement of myosin cortex organisation. Well-defined foci lead to higher CV values, whereas a dispersed appearance leads to lower values**. E.** Anterior-Posterior (AP) flow velocity of the NMY-2 cortex (see methods) for the specified conditions. **F.** Cortical confocal images of NMY-2 in polarity establishment CDC-42, CDC-42(S71A) and CDC-42(S71E) zygotes in control, *mrck-1* or *aPKC (RNAi)* depletions. Here endogenous CDC-42 is depleted by 50% RNAi of its 3’ UTR (50% dilution of RNAi bacteria), to be able to combine with 50% control RNAi, *mrck-1 (RNAi)* or *aPKC (RNAi)* (see methods). Penetrance of phenotypes observed (qualitative analyses) indicated on images. **G.** Normalised NMY-2 foci CV (average of wild type set to 1) of the conditions shown in F. **H.** AP flow velocity of the NMY-2 cortex in CDC-42, CDC-42(S71A) and CDC-42(S71E) zygotes upon *control (RNAi)*. **I.** Schema summarising the conditions that led to prominent or dispersed foci; and showing how CDC-42 phosphorylation could act as a switch between these foci states by regulating aPKC and MRCK-1 function. All box plots show median ± IQR and all data points. Unpaired, two-tail Student’s T test *p<0.05, **p<0.01, ***p<0.001, ****p<0.0001, ns not significant. Scale bar: 10 µm. See also Figure S7.

### CDC-42 phosphorylation on serine 71 impacts the organisation of the actomyosin cortex through regulation of MRCK-1 and aPKC

CDC-42 pS71 localisation to actomyosin foci during polarity establishment was intriguing as CDC-42 and its downstream kinase, MRCK-1, have been implicated in the regulation of the actomyosin cortex, especially during polarity maintenance phase.^62,64^ In establishment, the cortex is a highly contractile network of actomyosin foci, interlaced by actin filaments, that is known to retract or ‘flow’ toward the anterior through the action of the small GTPase RhoA and its downstream kinase Rock (RHO-1 and LET-502 respectively in *C. elegans*). ^51–53,88,89^ RhoA signalling is also involved in the pulsatile behaviour of the actomyosin foci, constantly undergoing assembly, contraction and disassembly.^54,55^

To determine if CDC-42 could be involved in any of these processes, we tested whether depletion of CDC-42 or MRCK-1 had any effect on cortex organisation during polarity establishment and if so, how it compared to depletion of LET-502. We quantified cortex organisation by measuring the coefficient of variation (CV) of endogenously tagged NMY-2::mKate2 in the anterior cortex, where a higher CV is representative of a cortex with bright well-defined clustered foci, while a lower CV represents a more uniformly dispersed distribution of NMY-2 (**Figure 5A-5D**). We observed that contrary to *let-502(RNAi)*, which led to a strong dispersion of the actomyosin network^54^, both *mrck-1(RNAi)* and *cdc-42(RNAi)* led to smaller and tighter foci than those observed in wild type, with *mrck-1(RNAi)* having a stronger effect (**Figure 5A-5D**). Interestingly, double depletion of LET-502 and MRCK-1 led to an intermediate phenotype between MRCK-1 and LET-502 individual depletions (**Figure 5 A-D**), suggesting that a balance of activity between these kinases is important for actomyosin cortex organisation during polarity establishment. Next, we analysed aPKC depletion, and observed that it also led to smaller NMY-2 foci, but in this case, the foci were partially dispersed and much dimmer than the foci observed upon MRCK-1 depletion (**Figure 5A-5D**). In each of these conditions, changes in the structure of NMY-2 foci coincided with changes in the anterior-directed actomyosin cortical flow (AP flow) (**Figure 5E**). In *mrck-1(RNAi)* we observed a slight but not statistically significant increase in flow, while aPKC depletion led to a strong decrease in flow. Overall these results show that two CDC-42 effector kinases, MRCK-1 and aPKC, have broadly opposite effects on the actomyosin cortex during polarity establishment. Furthermore, the intermediate phenotype (based on foci CV and flow velocities) observed in *cdc-42(RNAi)* compared to *mrck-1(RNAi)* and *aPKC (RNAi)*, supports that CDC-42 is promoting both these kinases in the organisation of actomyosin foci. These results raised the question of how the activities of these kinases are coordinated.

In mammalian cells, S71 phosphorylation has been shown to regulate differently the interaction of Cdc42 with downstream effectors, depending on the effector.^76^ We therefore decided to analyse the actomyosin cortex organization in our CDC-42 phosphomutants. In non-phosphorylatable CDC-42(S71A) zygotes, NMY-2 foci were better delineated than those observed in control CDC-42 zygotes (**Figure 5F and 5G**), presenting a phenotype similar to MRCK-1 depletion (**Figure 5A)**. Conversely, phosphomimetic CDC-42(S71E) presented partially dispersed NMY-2 foci and weaker flow, similar to the phenotype observed upon aPKC depletion (**Figure 5A, 5D, 5E and Figure 5F-5H**). These phenotypes suggest that CDC-42 phosphorylation could aid the organisation of the actomyosin cortex by differentially regulating the coupling of CDC-42/aPKC and CDC-42/MRCK-1, with non-phosphorylated CDC-42 preferentially promoting aPKC function and phosphorylated CDC-42 promoting MRCK-1. This interpretation agrees with our findings indicating that CDC-42 phosphorylation disrupts CDC-42/aPKC association and aPKC activity (**Figure 2 and 3**), while it might not have an effect on CDC-42/MRCK-1 association, as suggested in mammalian cells.^76^ We propose that aPKC and MRCK-1 compete for a limiting pool of active CDC-42 and that CDC-42 phosphorylation would promote CDC-42-mediated activation of MRCK-1 by displacing aPKC from CDC-42.

To further test these functional relationships, we depleted MRCK-1 and aPKC in the CDC-42 variant strains. Depletion of MRCK-1 or aPKC in embryos expressing control GFP::CDC-42 caused similar phenotypes as in a wild-type background, albeit with small differences that may reflect differences in function relative to endogenous CDC-42 (**Figure 5F and 5G**, compare to **Figure 5A and 5D)**. More importantly, depletion of MRCK-1 in CDC-42(S71A) zygotes did not lead to any significant change in the CDC-42(S71A) phenotype (**Figure 5F and 5G**), indicating that MRCK-1 function is already perturbed in CDC-42(S71A). Moreover, aPKC depletion in CDC-42(S71A) zygotes led to disperse foci (**Figure 5F and 5G**), suggesting that the tighter foci observed in CDC-42(S71A) require aPKC. These results are consistent with our hypothesis that non-phosphorylated CDC-42 couples preferentially to aPKC and not MRCK-1. In the CDC-42(S71E), depletion of MRCK-1 led to tight foci in 50% of the zygotes (**Figure 5F and 5G**), indicating that in this strain the observed foci dispersion is partially due to MRCK-1, but also likely due to aPKC inactivation and hence CDC-42(S71E) phenotype cannot be fully rescued by removal of MRCK-1 alone. Depletion of aPKC in the CDC-42(S71E) background led to a slight increase in network dispersion (**Figure 5F and 5G**). Overall, these results suggest that the actomyosin foci organisation during polarity establishment phase depends on a balancing act between aPKC and MRCK-1 function, which is regulated by the turnover of CDC-42 phosphorylation (**Figure 5I**). This supports a role for CDC-42 in actomyosin organisation during polarity establishment.^52,56^

During polarity maintenance phase, the actomyosin network disassembles and NMY-2 forms an anterior cap of homogenously distributed puncta (**Figure S7A**). We have observed that, as previously described, CDC-42 and MRCK-1 are required for the cortical recruitment of NMY-2 during this stage (**Figure S7A and S7B**).^52,53,62,64^ Furthermore, depletion of aPKC led to loss of NMY-2 cap asymmetry, and reduced cortical NMY-2 levels throughout the zygote (**Figure S7A and S7B**). Analysing the CDC-42 variant strains to determine if CDC-42 phosphorylation could be regulating the actomyosin cortex at maintenance, we observed that, in both CDC-42(S71A) and CDC-42(S71E) mutants the cortical localisation of NMY-2 was strongly reduced, similar to that observed upon depletion of CDC-42 (**Figure S7C and S7D**). The proposed regulation of aPKC and MRCK-1 by CDC-42 phosphorylation in establishment might still be operative in maintenance phase, but because both kinases are required to sustain the anterior NMY-2 cap, the cap is absent in both phosphomutants. We therefore propose that the turnover of CDC-42 phosphorylation is important to allow the activity of both kinases in NMY-2 cap formation during polarity maintenance.

In summary, our data shows that phosphorylation of CDC-42 is involved in the organisation and dynamics of the actomyosin cortex likely through regulation of aPKC and MRCK-1 function.

## DISCUSSION

We previously proposed that in the *C. elegans* zygote, aPKC activity and localisation depend on a dynamic cycle of PAR-6/aPKC shuttling between PAR-3 and CDC-42 (aPKC cycle, **Figure 6**). ^21^ During polarity establishment, PAR-6/aPKC, initially localised to the entire zygote’s membrane, are carried towards the anterior on PAR-3 clusters, which are efficiently advected by the actomyosin flow.^80^ We hypothesised that while on these clusters PAR-6/aPKC are passed temporarily onto CDC-42 to allow short bursts of diffusive and active aPKC.^21^ While bound to CDC-42, aPKC can exclude posterior PARs from the membrane, however in this diffusive state, aPKC is less efficient at tracking actomyosin flow and would eventually lose its asymmetry. Thus, PAR-6/aPKC must cycle back to PAR-3 to efficiently polarise the zygote.

**Figure 6.**
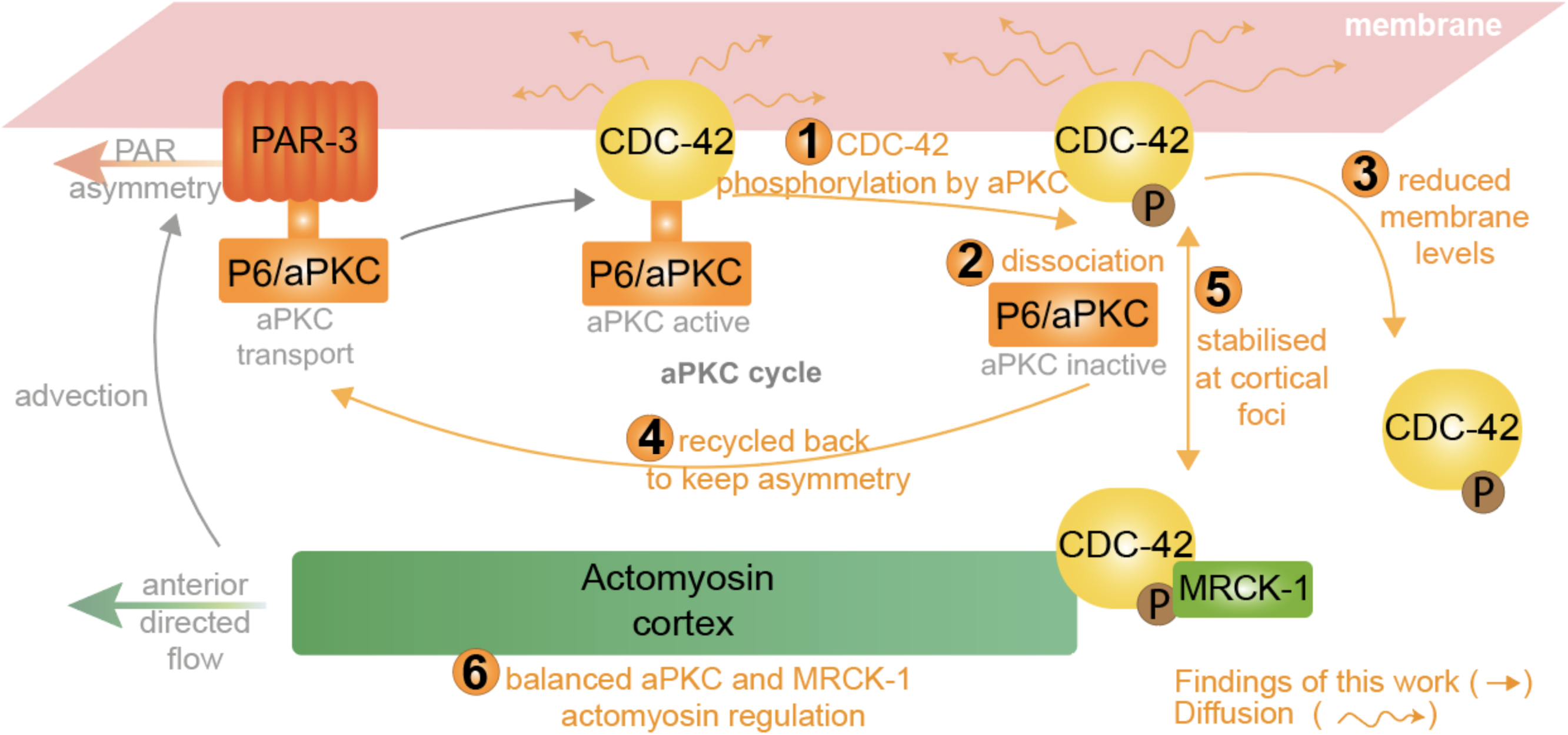
Model for aPKC self-patterning through phosphorylation of CDC-42. aPKC phosphorylation of CDC-42 (**step 1**, Figure 1A & 1C and Figure 4A & 4B), leads to CDC-42/aPKC dissociation (**step 2**, Figure 2). Released aPKC can then associate with PAR-3 clusters (**step 4**, Figure 3E), which promote aPKC asymmetry. Phosphorylated CDC-42 is destabilised at the membrane (**step 3**, Figure 1D-1F), except at actomyosin foci (**step 5**, Figure 4). Turnover of CDC-42 phosphorylation does not only promote the aPKC cycle but it is also needed for the correct actomyosin cortex organisation (**step 6**, Figure 5).

Here we have found that aPKC through phosphorylation of CDC-42 promotes the PAR-6/aPKC cycle by limiting the lifetime of PAR-6/aPKC/CDC-42 complexes. This constitutes an aPKC self-regulatory mechanism that could ensure aPKC optimal localisation and activity levels throughout polarisation. Our results indicate that phosphorylation of CDC-42 by aPKC (**step 1** in **Figure 6**) destabilises the CDC-42/PAR-6/aPKC complex (**step 2 & 3**) and promotes the recycling of PAR-6/aPKC back to the anterior-segregating PAR-3 clusters (**step 4**), maintaining in this way aPKC anterior localisation. Through disruption of the CDC-42/PAR-6/aPKC complex, aPKC not only limits its localisation but also its activity on posterior PAR exclusion and actomyosin cortex regulation (**step 5 and 6**). Subsequent CDC-42 dephosphorylation would revert these processes and allow the cycle to continue. We propose that, fuelled by the turnover of CDC-42 phosphorylation, PAR-6/aPKC will continuously cycle between active (CDC-42/PAR-6/aPKC) and anteriorly transported (PAR-3/PAR-6/aPKC) states, ultimately leading to the polarisation of the zygote. Following we will discuss the steps of our working model (**Figure 6**).

Here, we show that aPKC-dependent phosphorylation of CDC-42 on S71 limits the lifetime of the CDC-42/PAR-6/aPKC complex (**step 1 & 2**). PAR-6 and aPKC interact with CDC-42 via the PDZ and semi-CRIB domains of PAR-6.^17,23,25,27,90,91^ Phosphorylation of S71, which is positioned between the effector protein-binding and the GTP-binding domains of CDC-42,^92^ is known to disrupt the interaction of Cdc42 with the CRIB domains of N-WASP and PAK1 in mammalian cells.^76^ We speculate that, in a similar way, S71 phosphorylation could disrupt the interaction between CDC-42 and PAR-6. Alternatively, aPKC might make direct interactions with CDC-42 that contribute to stable assembly of a CDC-42/PAR-6/aPKC complex and are released upon completion of aPKC’s catalytic cycle. Similar aPKC “substrate phosphorylation and release” mechanisms have been proposed for other aPKC polarity targets. In epithelial cells phosphorylation of Par3 by aPKC weakens the association between Par3 and aPKC, and excludes Par3 from the apical membrane.^20,37,93–95^ In the *Drosophila* neuroblast, phosphorylation of LGL by aPKC also leads to the dissociation of LGL from Par6/aPKC^19,96^, and in the *C. elegans* zygote, LGL phosphorylation is thought to mediate the release of the whole PAR-6/aPKC/LGL complex from the membrane.^61^ The aPKC expansion we observed in the presence of the non-phosphorylatable CDC-42 mutant would be consistent with the existance of a similar mechanism for CDC-42/PAR-6/aPKC.

### CDC-42 phosphorylation and the regulation of aPKC activity and asymmetric localisation

CDC-42 is known to promote aPKC activity and diffusion at the membrane;^21–23,25,26^ hence, aPKC-mediated dissociation of PAR-6/aPKC from CDC-42 (**step 2**) could constitute a negative feedback mechanism. This negative feedback would not only restrict aPKC activity but also its domain of action by promoting aPKC association with PAR-3 (**step 4**). This aPKC self-regulatory mechanism could ensure optimal activity and localisation of aPKC during polarisation. aPKC could limit its activity by destabilising the CDC-42 complex it resides in, or alternatively aPKC phosphorylation of nearby CDC-42 molecules would prevent other aPKC molecules to form stable interactions with CDC-42. Our measured k_off_ of 0.099 s^-1^ for the CDC-42(S71E)/aPKC complex suggests an average lifetime of ∼7 seconds for these complexes once phosphorylation occurs, compared to ∼12 seconds for the unphosphorylated complex (k_off_ = 0.057 s^-1^). Depending on the catalytic rates of active aPKC, these lifetimes could still be adequate to allow aPKC to phosphorylate other substrates, especially those that are kinetically more favourable, accessible or abundant than CDC-42.^24^ It is noteworthy that CDC-42 does not present an optimal aPKC consensus site (**Figure 1B**), so other aPKC substrates could be more efficiently phosphorylated.

We have previously reported *ex vivo* kinetic measurements that suggest CDC-42/aPKC complexes are more stable during polarity maintenance (k_off_ = 0.050 s^-1^, lifetime ∼14 sec) than establishment (k_off_ = 0.109 s^-1^, lifetime ∼6 sec).^36^ Strikingly, the phosphomimetic and non-phosphorylatable mutants of CDC-42 exhibited CDC-42/aPKC lifetimes similar to those observed in establishment and maintenance phase, respectively (**Figure 2F**). Although this is only a correlation and we acknowledge that other factors could be in place, our data suggests that different levels of CDC-42 phosphorylation could account for the different stability of the CDC-42/PAR-6/aPKC complex observed in establishment vs. maintenance. In establishment phase, PAR-6/aPKC are mostly observed in anteriorly segregating clusters with PAR-3^21,41,72^ a state that would be promoted by phosphorylation-mediated dissociation of CDC-42/PAR-6/aPKC complexes. In agreement with this, we observed that aPKC is more asymmetric and exhibits a more clustered organisation, highly co-localising with PAR-3, in the CDC-42(S71E) variant strain (**Figure 3D-3G**). In maintenance phase, anteriorly localised PAR-6/aPKC diffuse more rapidly on the membrane^21,41,72^ and depend more strongly on CDC-42 for their membrane localisation^21,66,72^, all in agreement with the observed stabilisation of the active CDC-42/PAR-6/aPKC complex in maintenance.^36^ This CDC-42 dependent state would be supported by lower levels of CDC-42 phosphorylation. Consistent with this, the CDC-42(S71A) variant induced a homogenous membrane distribution of aPKC (**Figure 3D-3G**) that mimics that of CDC-42 itself. Overall, our results indicate that by tuning CDC-42 phosphorylation the system could be favouring aPKC asymmetric transport in establishment vs. aPKC activation in maintenance. We do not know the underlying mechanisms leading to these hypothesized different CDC-42 phosphorylation levels in these distinct polarity phases, but differential regulation of aPKC activity or different availability of CDC-42, competing substrates or interactors could be involved.^24,30^

### Regulation of CDC-42 membrane localisation by CDC-42 phosphorylation

aPKC is known to inhibit the association of some of its substrates with the membrane by phosphorylating and neutralising residues in short polybasic hydrophobic sequences (BH motifs) involved in membrane targeting.^97,98^ However, the CDC-42 S71 residue is not present in a BH motif (analysed using the BH scoring algorithm^99^); and, moreover, membrane localisation of CDC-42 depends mostly on C-terminal polybasic residues and a prenylated cysteine.^32,34,100,101^ Therefore, it is unlikely that S71 phosphorylation regulates the membrane localisation of CDC-42 in this direct way. We propose that S71 phosphorylation destabilises CDC-42 membrane localisation and asymmetry (**step 3**) through disruption of CDC-42 interaction with PAR-6/aPKC or with other CDC-42 partners. This idea is supported by the positive feedback described in *Drosophila* neuroblast and in the *C. elegans* zygote, where PAR-6 promotes robust membrane localisation of CDC-42 (**Figure S2**).^22,64^ PAR-6/aPKC could stabilise CDC-42 at the membrane by providing additional membrane contact sites, for example the recently identified polybasic domain of aPKC that is capable of targeting PAR-6/aPKC to the plasma membrane.^39,40^

The enrichment of phosphorylated CDC-42 at actomyosin foci (**step 5**), suggests that phosphorylated CDC-42 interacts in some way with the actomyosin cortex. We do not know if the phosphorylated state of CDC-42 is recruited to actomyosin foci or alternatively stabilised at the membrane by these foci (indicated by **step 5** double arrow). These processes could be mediated by known interactions of CDC-42 with actomyosin regulators. ^84,85,102^ Interestingly, aPKC phosphorylation-dependent recruitment of proteins to the actomyosin cortex has also been proposed in the *Drosophila* neuroblast. Here, the scaffold protein Miranda, involved in the subcellular distribution of cell-fate determinants, is phosphorylated by aPKC. This leads not only to Miranda release from the apical membrane, where aPKC resides, but also to the basal recruitment of Miranda to a specific domain of the actomyosin cortex.^103^ aPKC-driven recruitment of polarity effectors to certain actomyosin cortex domains might be an emerging mechanism in the creation of cellular asymmetries and it could also provide aPKC with the means to control actomyosin cortical dynamics.

### aPKC and CDC-42 roles in actomyosin cortex organisation and dynamics

The localisation of phosphorylated CDC-42 to actomyosin foci and the identified role for CDC-42 in actomyosin foci organisation (**step 6**) is intriguing. There is a clear regulatory feedback between PARs and the actomyosin cortex during polarity establishment,^51,52,104^ but the molecular mechanism by which PARs influence actomyosin remains unknown. Our results suggest that aPKC-dependent phosphorylation of CDC-42 may contribute to this feedback mechanism. During polarity establishment the actomyosin cortex is predominantly controlled by RhoA (RHO-1 in *C. elegans*) signalling, driving its anterior directed movement (flow) and orchestrating the pulsatile behaviour, assembly and disassembly, of actomyosin foci.^54,55^ Our data suggests that CDC-42 is also important for correct actomyosin foci organisation and flow, the turnover of CDC-42 phosphorylation likely promoting actomyosin cortical dynamics by balancing the opposite roles of the CDC-42 effector kinases MRCK-1 (foci dispersion) and aPKC (foci formation). A possible mechanism for this, suggested by structural studies^90,105^, could be the existence of a competition between PAR-6/aPKC and MRCK-1 for CDC-42 binding. In this manner, CDC-42 phosphorylation, by promoting the dissociation of CDC-42/PAR-6/aPKC, would not only inactivate aPKC but also promote CDC-42 interaction with and activation of MRCK-1. Note that in mammalian cell culture CDC-42 phosphorylation does not seem to prevent CDC-42 interaction with MRCK-1^76^, supporting the idea that CDC-42 phosphorylation could mediate a CDC-42 effector switch from aPKC to MRCK-1.

At first glance, CDC-42 phosphorylation seems to have opposing effects on the generation of PAR asymmetry during polarity establishment. On the one hand, it promotes the PAR-3/PAR-6/aPKC clusters that segregate anteriorly with the actomyosin flow, whereas on the other hand, CDC-42 phosphorylation seems to reduce actomyosin foci organisation and flow. This apparent contradiction can be resolved by considering the constant turnover of CDC-42 phosphorylation, which likely occurs on timescales of a few seconds. aPKC, in its active state with CDC-42, would promote actomyosin foci contraction and flow. Upon reaching its peak of activity, aPKC would phosphorylate CDC-42, then dissociate from it leading to aPAR cluster formation. Once aPKC becomes dissociated from CDC-42, CDC-42 could promote MRCK-1 activation and actomyosin foci relaxation. In this manner the constant turnover of CDC-42 phosphorylation in the developing anterior domain could orchestrate successive waves of actomyosin foci contraction, aPAR clustering, and actomyosin foci relaxation, that would drive the overall anterior-directed flow of the cortex and aPARs. A more detailed spatio-temporal study of aPAR cluster formation together with actomyosin foci evolution is necessary to fully understand this process.

During maintenance phase, actomyosin foci disperse and both CDC-42 effector kinases are required to form the actomyosin anterior cap. We propose that at this stage, CDC-42 phosphorylation promotes actomyosin cap formation also by alternating the activation of aPKC and MRCK-1, which at this stage seem to cooperate in the organisation of the actomyosin cortex rather than oppose each other, as observed during polarity establishment. We do not know how this change in kinase roles happens between establishment and maintenance phases. However, a differential regulation of the actomyosin cortex between these polarity phases is not unexpected considering the dramatically different cortex architecture between establishment and maintenance and the reduced involvement of RhoA pathway during maintenance phase (**Figure S7**).^52,53^

In summary, we have identified a negative feedback in the polarisation of the *C. elegans* zygote - aPKC restricts its own localisation and activity through phosphorylation of CDC-42. Self-limiting mechanisms can prevent systems from over-shooting an optimal level of activity and could provide homeostasis and adaptability to cell polarity processes.^106–108^ aPKC and CDC-42 have widespread roles in cell polarity and phosphorylation of CDC-42 on S71 has been observed in other polarised cells, for example at the leading edge of migrating cells^77^, and in epithelial cells.^75^ Therefore, we predict that the identified aPKC-self regulatory feedback will be a widely conserved mechanism ensuring robust cell polarisation.

## METHODS

### *C. elegans* strains and maintenance

The strains used in this study are listed in strain table (key resources table). All strains were maintained on nematode growth media (NGM) under standard conditions.^109^ Temperature sensitive strain *pkc-3(ne4246)* was maintained at 15 °C and shifted to 25°C for aPKC inactivation. The rest of reporter the strains were maintained at 25 °C.

### *C. elegans* transgenic animals

Strains expressing GFP::CDC-42 (JRL1685), GFP::CDC-42(S71A) (JRL1686) and GFP::CDC-42(S71E) (JRL1687) constructs were generated for this study with the Mos-1 mediated single copy transgene insertion method (MoSCI).^110,111^ Genomic cdc-42 was amplified from genomic worm extract using the following primers (fwd: ggggacaagtttgtacaaaaaagcaggctcgatgcagacgatcaagtgc; rev: ggggaccactttgtacaagaaagctgggtctagagaatattgcacttcttcttct) and inserted into the pDONR221 plasmid (#12536017, Invitrogen) using the BP clonase II enzyme (#11789100, Invitrogen) to generate pDONR221-gCDC42attb. The plasmid was introduced by heat shock into DH5alpha and extracted with standard Miniprep (#K0503, ThermoFisher). The S71A and S71E mutations were introduced into the pDONR221-gCDC42attb plasmid with the QuickChange II XL Site Directed Mutagenesis Kit (#200521, Agilent Technologies) with the following primers, fwd S71A: gatcgattaaggcctcta**gcc**tatccacagaccgacgtg; rev S71A: cacgtcggtctgtggata**ggc**tagaggccttaatcgatc; fwd S71E: cgatcgattaaggcctcta**gag**tatccacagaccg acgtc; rev S71E: cacgtcggtctgtggata**ctc**tagaggccttaatcgatcg. The resulting plasmids were inserted into XL10-Gold Ultracompetent Cells (#200314, Agilent) via heat shock as indicated by the provider and extracted with standard Miniprep (#K0503, ThermoFisher). The resulting plasmids were used in an LR reaction (Gateway LR Clonase II Enzyme Mix, # 11791-020 ThermoFisher) with plasmids pCFJ210 (#30538 Addgene), pJA245 #21506 Addgene), pCM1.36 (#17249 Addgene). This reaction created a plasmid for MoSCI insertion in chromosome I (4348) containing mex-5 promoter and GFP sequence at the N-term of *cdc-42*, and *tbb-2* 3’UTR at its C-term. The resulting plasmid was co-injected with the transposase under *pie-1* promoter (pCFJ103),^111^ and two fluorescent reporters (pCFJ04, pCFJ90) into the gonad of young adult worms presenting *unc-119(ed3);ttTi4348 genotype* (EG6701 strain, CGC). Wild type moving worms without red fluorescence were selected for sequencing, to confirm the presence of the desired construct.

### Bacterial Strains

OP50 bacteria were obtained from CGC. Feeding by RNAi was performed with HT115(DE3) bacteria strains containing the indicated RNAi plasmid from the Ahringer library^112^ (key resources table). The RNAi clone that targets the 3’UTR of *cdc-42* was generated through amplification of genomic *cdc-42* 3’UTR (Fwd: gaagatctgaacgtcttccttgtctccatgt; Rev: ggggtaccacgtaacggtgtatccggac) and insertion into the L4440 plasmid (#1654 Addgene). Control bacteria is transformed with empty L4440 vector (#1654 Addgene).

### *C. elegans -* RNAi Feeding

HT115(DE3) RNAi clones were inoculated from LB-agar plates (10 μg/ml of carbenicillin, 10 μg/ml tetracycline, 100 U/ml nystatin) into 5 ml LB liquid cultures (10 μg/ml of carbenicillin, 10 μg/ml tetracycline, 100 U/ml nystatin) and grown over night at 37°C with agitation. The bacteria were induced for 4 h with 4 mM IPTG at 37 °C and concentrated five-fold before seeding 300 μl of the bacterial culture onto NGM plates (10 μg/ml of carbenicillin, 10 μg/ml tetracycline, 1mM IPTG, 100 U/ml nystatin). L4 larvae were added to RNAi feeding plates and incubated for 72 h at 15 °C. For temperature sensitive *pkc-3(ne4246)* line, the worms were shifted to 25 °C for 2 h before fixing or imaging. RNAi bacteria were not diluted (we refer to it as 100% RNAi) unless stated (partial depletions). For partial depletion of *cdc-42*, the RNAi bacteria clone targeting the 3’UTR of *cdc-42* was mixed at a 1:1 ratio with control bacteria (transformed with empty L4440 vector). For partial depletion of *par-6*, the RNAi bacteria clone targeting *par-6* was mixed at a 1:1 ratio with bacteria targeting the 3’UTR of *cdc-42*. For *rho-1* depletion *rho-1* RNAi clone was mixed with control bacteria at a 1:10 ratio. For double depletions (i.e. *let-502* and *mrck-1; cdc-42* and *mrck-1; cdc-42* and *pkc-3)* bacteria RNAi clones targeting the corresponding gene product were mixed at 1:1 ratio.

### *C. elegans -* HaloTag labelling in combination with RNAi feeding

RNAi NGM plates were prepared as indicated above and L4 larvae were added to RNAi feeding plates and incubated for 56 h at 15 °C. Adult worms where then grown in S-medium liquid culture with RNAi bacteria and with JFK549 HaloTag (10μM, Janelia Farm) overnight at 15°C and under rotation (30-50 worms in 65ul culture in a 0.5ml tube). The RNAi bacteria derived from a pellet of a 1.5 ml LB culture grown for 8 hours at 37°C (10 μg/ml of carbenicillin, 10 μg/ml tetracycline, 1mM IPTG, 100 U/ml nystatin), which was resuspended in 200 μl of S-medium with antibiotics and IPTG (150 mM NaCl, 1 g/L K2HPO4, 6 g/L KH2PO4,5 mg/L cholesterol, 10 mM potassium citrate pH 6.0, 3 mM CaCl2, 3 mM MgCl2, 65 mM EDTA, 25 mM FeSO4, 10 mM MnCl2, 10 mM ZnSO4, 1 mM CuSO4, 10 μg/ml of carbenicillin, 10 μg/ml tetracycline, 100 U/ml nystatin, 1mM IPTG). Following the overnight incubation in JFX549, adults were washed six times in M9 buffer (22 mMKH2PO4, 42 mMNaHPO4, 86 mM NaCl and 1 mMMgSO4) with Triton X-100 (0.01%) and then plated onto a fresh RNAi NGM plate until imaging.

### *In vitro –* aPKC Kinase Assays

*In vitro* kinase assays were performed in the presence or absence of different recombinant human atypical PKC (50 or 100 ng of PKC ζ, #14-525 Merk Millipore; 1µg of PKC I/αPar6 complex, gift from Shona Ellison, Mc Donald lab, Crick Institute) and with different recombinant Cdc42 (3µg of human Cdc42, #CD01 Cytoskeleton; 3µg of recombinant MBP tagged-*C. elegans* CDC-42 or CDC-42(S71A)), which were incubated for 1 hour at 30 °C in 30 µl kinase-assay buffer (25 mM Tris pH 7.5, 25 mM NaCl, 5 mM MgCl2, 0.5 mM EGTA, 1 mM DTT) containing 1mM ATP-gamma-S (# ab138911,Abcam). The phosphorylation reaction was stopped with 10mM EDTA, followed by alkylation for one hour at room temperature with 1.5 mM PNBM (p-nitrobenzyl mesylate, #ab138910 Abcam). Reactions were then stopped in 1x Laemmli buffer at 95 °C for 5min.

Samples were processed for Western-blot and phosphorylation was detected using an anti-thiophosphate ester antibody (#ab92570, Abcam).

### *In vitro –* Purification of CDC-42 recombinant proteins

MBP::CDC-42 and MBP::CDC-42(S71A) recombinant proteins were generated for this study Briefly, MBP::CDC-42 coding sequence was inserted into the pMAL-c5X plasmid (NEB). The S71A mutation was introduced into the pMALc-MBP::CDC-42 plasmid with the QuickChange II XL Site directed mutagenesis kit (#200521, Agilent technologies) with the following primers, fwd S71A: gatcgattaaggcctcta**gcc**tatccacagaccgacgtg; rev S71A: cacgtcggtctgtggata**ggc**tagaggccttaatcgatc. For protein production NEB Express Competent E coli (#C2523H) were transformed with the resulting plasmids. An overnight culture was used to inoculate 500 ml of 2TY medium and grown at 37 °C until an OD of 0.5 at 600 nm was reached. Then cultures were induced with 0.25 mM IPTG for 19 h at 18 °C with agitation (210 rpm). Bacteria were lysed with sonication and supernatant obtained after 20.000 xg for 30 min at 4 °C centrifugation. Supernatant was diluted 1:6 in column buffer (20 mM Tris pH7.4, 0.2 M NaCl, 1 mM EDTA) and passed through an amylose resin column (NEB#E8021L). Columns were washed prior to recombinant protein elution with 10 mM maltose in column buffer.

### *In vitro –* Purification of aPKCi-PAR6 recombinant proteins

Full length human PAR6alpha with an N-terminal Strep-Strep-TEV tag was co-expressed in Expi293F cells with untagged full length human aPKC*iota* (aPKC1). Expi293F cells were transfected at a density of 1-3x10^6^ viable cells/mL using 3ug polyethylenimine (PEI) per 1ug plasmid DNA and incubated for 96 hours at 37 °C, 8% CO2 with continuous shaking at 125 rpm. Recombinant protein was purified in 50mM HEPES, 150mM NaCl and 0.5mM TCEP at pH7.5 supplemented with protease inhibitors (#11873580001 cOmplete, Roche). Cells were sonicated, then clarified by high-speed centrifugation and lysates were incubated with Strep-Tactin Sepharose affinity beads (#2-1201-010, IBA) for 2 hours to allow tagged PAR6 to bind along with aPKC1 bound in complex. The beads were recovered in an PD-10 column using a vacuum manifold and washed with 5 x column volumes (CV). Recombinant protein was eluted with buffer containing 0.25mM D-desthiobiotin 50mM HEPES, 150mM NaCl and 0.5mM TCEP at pH7.5. Eluted fractions were loaded onto a size exclusion Superdex® 200 Increase 10/300 column (#17-5175-01, Cytiva) and peak fractions were collected and assessed using SDS-PAGE. Fractions containing aPKCi-PAR6 complex were pooled and concentrated to 0.36mg/ml.

### C. elegans embryos – Western Blots

Embryos were obtained by a standard bleaching protocol and resuspended in NuPAGE LDS sample buffer (Invitrogen) prior to sonication using the Biorupture (Diagenode) for 5 cycles – 30 sec on, 30 sec off. Samples were heated at 70 °C for 10 min before centrifugation at 13000 rpm for 20 min to obtain cleared supernatants. Samples were run on a 12% NuPAGE gel using MOPS SDS running buffer (Invitrogen) and transferred in semi-dry conditions onto PVDF membrane (Immobilon-P membrane 0.45 μm, Millipore). A primary anti-GFP (#11814460001 Roche, 1:1000) was used to detect the different GFP constructs. A primary anti-HSP-60 was used for the loading control (DSHB, P50140 - CH60_CAEEL, 1:1000). Detection of thiophosphate ester tagged proteins from the *in vitro* kinase assay was done with a specific anti-thiophosphate ester antibody (#ab92570 Abcam 1:10000). Antibody against total CDC-42 was use for the loading control (#sc-390210, Santa Cruz 1:1000). Secondary antibodies indicated in the key resources table were used as recommended by provider. The blots were revealed via chemiluminescence (# GERPN2236 ECL prime, GE Healthcare Life Sciences). Western blot bands were analysed using Fiji image analyses software.

### C. elegans zygote - Immunofluorescence

Immunofluorescence was performed as previously described.^113^ Briefly, adult worms were collected and washed with M9 buffer (22 mM KH_2_PO_4_, 42 mM NaHPO_4_, 86 mM NaCl and 1 mM MgSO_4_). 30 worms (in 7-10 μl of M9 suspension) were transferred to a microscopy slide (# 10-2066a Erie Scientific) coated with 0.1% poly-lysine. Embryos were released from the adult worms using a needle and then compressed with a coverslip, before being snap-frozen on dry ice for 30 minutes. The coverslip was then removed, and the slide fixed in methanol at room temperature for 30 minutes, following rehydration with PBS (two 10 min washes) and PBS with 0.1% Tween 20 (10 minutes) before proceeding with antibody incubation, DAPI staining and Mowiol mounting (#81381 Sigma). All antibodies used are listed in key resources table. Primary antibodies dilutions used: anti-PAR-2 (1:500, Dong et al., 2007)^114^ anti-PAR-3 (1:50, # P4A1 Developmental Studies Hybridoma Bank) and anti-PKC-3 (1:500, Tabuse et al., 1998)^6^, anti-CDC-42 pS71 (1:500, #44214G Invitrogen), anti-NMY-2 (1:25.000, Ahringer Lab). Secondary antibodies were used as recommended by provider.

Confocal images were capture with a Nikon A1R+ scanning confocal on a Nikon Ni body, equipped with a PlanApochromat 63x 1.4 NA oil-immersion lens. The system presents GaAsP (for green and red channels) and PMTs (for blue and far rad) detectors and runs with Nikon elements software. Immunofluorescence images in **Figure 3B**, **3D** and **3H** and **Figure S5G** and **S5H** were captured in a spinning disk with a bespoken 3i system (see specs bellow – live imaging section)

### *C. elegans –* Live Imaging

Embryos were dissected in 3 μl of egg buffer (2 mM CaCl2, 118 mM NaCl, 48 mM KCl, 2 mM MgCl2, 25 mM HEPES (pH 7.4)) on top of a 30 mm circular coverslip (#10343435 Fisher Scientific). The coverslip with the dissected worms was then inverted onto 2% agar pads in egg buffer solution (2 mM CaCl2, 118 mM NaCl, 48 mM KCl, 2 mM MgCl2, 25 mM HEPES (pH 7.4)) placed in a custom-made temperature control stage holder (Bioptechs Oasis Cooling System). To gain further sample temperature stability the objective temperature is also controlled with an objective cooling collar (Bioptechs Oasis Cooling System). Sample temperature is set at 20 °C unless stated otherwise.

Confocal images were capture with a PlanApochromat 100x 1.4 NA oil-immersion lens on a Leica DMRBE (Leica) equipped with an CSU-X1 A1 Spinning Disk Confocal (Yokogawa) and LaserStack v4 Base holding 405, 488, 561 and 640 nm lasers (3i, Intelligent Imaging and Innovation). Images are acquired with a Prime 95B back-illuminated scientific CMOS and the system runs with SlideBook 6 software (3i).

For GFP::CDC-42 (and S71 phosphomutants) live imaging, embryos had midplane images taken at establishment (male pronuclei in close proximity to the posterior cortex), maintenance (between pronuclear meet and metaphase) and 2-cell stage with the 488 laser (**Figure 1D-1F** and **Figure S2**). For simultaneous observation of GFP::CDC-42 (and S71 mutants) and NMY-2::mKate, cortical images were capture at establishment phase (male pronuclei in close proximity to the posterior cortex or starting pronuclear migration) with the 488 and 561 laser, both with single pass filters to minimise bleed-through (**Figure S6C**). For HaloTag::aPKC labelled with JFK549 HaloTag (Janelia Farm), cortical and midplane sections of embryo were taken at establishment, maintenance and 2-cell stage with the 561 laser (**Figure S5A-F**). For PIV analysis of NMY-2::mKate foci, cortical sections of embryos were imaged every second with the 561 laser.

### Single-molecule near-TIRF microscopy in live embryos (HILO)

Imaging was performed on a bespoke single-molecule TIRF microscope constructed around a Nikon Ti-E microscope body, using an Obis 488 laser set to 20 mW, beam expanded to ∼20 µm and using highly inclined illumination (HILO). Imaging was performed using Photometrics Evolve 512 at 100 nm/pixel with a 100x Nikon 1.49NA TIRF objective lens. All images of GFP::CDC-42 and CDC-42 mutant variants were taken of the anterior cortex of maintenance phase zygotes (between pronuclear meet and metaphase and for up to 15 s), subarraying the camera sensor to a ROI of 128x128 pixels to achieve 15ms/frame exposure time. PAR-6::GFP was also imaged under the same conditions as the GFP::CDC-42 variants for their comparison in **Figure 1H**.

### Single-cell, single-molecule pull-down in lipid nanodiscs

CDC-42 lipid nanodisc pull-downs were performed exactly as described.^36^ Briefly, *C. elegans* nematodes expressing the desired GFP::CDC-42 variant and carrying endogenously tagged HaloTag::aPKC were cultured overnight at 20 °C in liquid culture containing 15 µM JF_646_ Halo ligand^115^ to label the HaloTag::aPKC molecules. Labelled worms were dissected in egg buffer and embryos were rinsed twice with nanodisc buffer (10 mM Tris (pH 8.0), 150 mM NaCl, 1% DIBMA (DIBMA 12 Tris, Cat# 18014 from Cube Biotech), and 0.1 mg/mL bovine serum albumin). A labelled zygote was transferred to a PDMS microfluidic device that had been passivated with PEG, functionalized with anti-GFP nanobodies, and equilibrated in nanodisc buffer. The device was sealed with clear tape and transferred to a custom-built TIRF microscope equipped with a pulsed 1064 nm laser for rapid cell lysis; a 60x, 1.49 NA Olympus TIRF objective; 488 nm and 638 nm lasers for excitation of GFP and JF_646_, respectively; a home-built 4-color image splitter; and a Photometrics PrimeBSI Express camera.^35^ A brightfield image was acquired to document the embryonic stage, and then the embryo was lysed with a single shot from the pulsed laser. Beginning immediately after lysis, TIRF images were collected at 20 frames per second for 500 s to detect binding of GFP::CDC-42 and HaloTag::aPKC molecules to the coverslip. To allow correction of k_off_ values for JF_646_ photobleaching, control experiments were performed on the same days using a transgenically expressed YFP::HaloTag protein to measure JF_646_ photobleaching rates. Images were processed to extract kinetic information as described below.

## QUANTIFICATION AND STATISTICAL ANALYSIS

### Image analysis-General

Images were analysed with FIJI^116^ and MATLAB (Mathworks). Secondary processing of images was performed using Photoshop and Illustrator (Adobe).

### Image analysis – Membrane intensity levels

To determine the membrane intensity of aPKC and CDC-42 of immunostained zygotes (**Figure 2C-2D and Figure S1D-S1E**) a 20-pixel wide stripe with the cell membrane at its centre was obtained from straightening the zygotes perimeter from anterior to posterior and back to anterior (custom-built FIJI macro to create cortical flat-outs from midplane images). The flat-out was then read by a custom-built MATLAB script. Each stripe was split in half and two anterior values were obtained per zygote. We analysed the first 20 x 100-pixel band at the anterior domain of each stripe. For each of the 1px columns in this band the average pixel intensity of the highest values encompassing the cortex (10 highest pixels for aPKC and 5 highest for CDC-42) were calculated. The average intensity of the anterior-most domain was then normalised by the cytoplasmic intensity. For CDC-42 this corresponded to the average value intensity of a 10 x 100 pixel band at the anterior most domain but positioned 30 pixels away from the membrane and in the cytoplasm. For aPKC, which also presents an asymmetry within the cytoplasm, we normalised aPKC membrane intensity with the average intensity of a cytoplasmic 10 x 50 pixel band at the posterior most domain (last 50-pixel column band).

For CDC-42 membrane intensity levels from live embryos (**Figure 1D-1E and Figure S2A, S2B, S2D and S2E**) each stripe was split in half and two values were obtained per zygote for each the anterior and posterior domain. We analysed a 60 x 60 pixel square at the anterior domain of each stripe, this was located 60 pixels away from the anterior pole (leftmost edge of the image) between pixels 60 to 120 in the x axis. For each of the 1 px rows in this band the average pixel intensity was calculated to give an average cross section of (from top to bottom) background-cortex-cytoplasm. The average intensity of the anterior most domain was then normalised by the cytoplasmic intensity. This corresponded to the average value intensity of a 60 x 10 pixel band within the anterior defined area but positioned between 50 and 60 pixels in the y axis (ie. the bottom of the image). Posterior analyses were conducted on the final 60 x 60 pixel region of the halved cortex and had the same cytoplasmic correction applied. Profiles and measurements were averaged between each half of the cortex to give a single measurement for each embryo.

### Image analysis – Intensity Profile Extraction

To plot CDC-42 profiles (**Figure 1F and Figure S2C and S2F**) a 60 x 60 area was selected cutting across a stripe of straightened cortex at the anterior most of each zygote. The intensity values in this area were averaged in the Y-axis leading to an intensity profile encompassing the cell membrane. The cytoplasmic level of each profile was set to one by dividing the profile values by the average of a cytoplasmic section (last 16 pixels). For representation, all normalised intensity profiles were centred around their highest values, corresponding to the membrane/cortex. This analysis was performed on images of maintenance stage embryos – defined as pronuclear meet to metaphase.

### Image analysis – Cortical intensity profile extraction

To plot aPKC cortical levels (**Figure S5B**) each embryo had a 100 pixel height ROI drawn along its cortex length - starting from the anterior. This was used to crop an image of the cortex running from anterior left to posterior right. These images were fed into MATLAB and analysed using a bespoke script. Briefly, a 100(H) x 50(W) pixel ROI is iterated (1 pixel step) along the length of the cropped image with the average intensity being measured at each iteration. Average intensity was then binned by percentile along the length of the cropped image to allow for standardization of cortex length between embryos. Percentile intensity was then averaged by condition and normalised using a HiLo ratio method. This sorts the data into ascending order then takes an average of the lowest ten values (lo) and the highest ten values (hi). The normalised data is then calculated as; 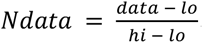, to give profiles that are constrained by both their internal maxima and minima.

### Image analysis – foci profiles

A line of 5 μm in length was drawn across CDC-42 pS71 foci of NMY-2::GFP zygotes stained for CDC-42 pS71 and PAR-3 or of wild-type zygotes stained for CDC-42 pS71 and aPKC. Plot intensity profiles along the drawn line were obtained for all co-labelled proteins (CDC-42 pS71, NMY-2 and PAR-3 or CDC-42 pS71 and aPKC) (**Figure 4F, Figure S6D**). Each profile was normalised by substracting background signal (average intensity of the profile ends, 0.5 μm at either end). Then the average plot profile for each label was plotted together with its SEM. In analysis where only NMY-2 foci were present, the 5μm length line was drawn across NMY-2 foci and then profiles of all the co-labelled proteins (CDC-42 variants, aPKC or PAR-3) were obtained and processed in the same way as indicated above (**Figure S6C, S6E**).

### Image analysis – Diffusion coefficient and track length

TIRF imaging data was analysed using bespoke MATLAB software^117^ to track and quantify the intensity of particles as a function of time. Briefly, centroids were determined using iterative Gaussian masking and intensity calculated using the summed intensity inside a circular region of interest (3 pixel radius), and corrected for the local background in a 10 pixel wide square region of interest around the particle. Particles were linked together in trajectories based on a linking distance of 4 pixels and accepted if their signal to noise ratio was above 0.4 and lasted longer than 3 frames. Diffusion coefficients were calculated by a linear fit to the first 4 mean-squared displacements as a function of time interval (τ).^118^ Probability distributions of diffusion coefficients were made from normalised histograms and fit using multiple Gamma functions.^119^ Here, a two Gamma function model comprising a mobile and immobile population to yield the diffusion coefficient of the mobile population and proportion of tracks in the immobile population was used:

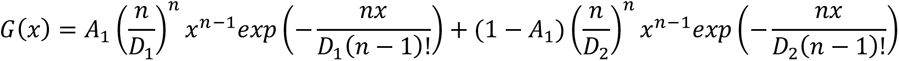

Where G(x) is the probability of a track having x diffusion coefficient; n is the number of mean squared displacement values used in the diffusion coefficient fits, A_1_ is the proportion of immobile particles and D_1_ and D_2_ are the population immobile and mobile diffusion coefficients respectively.

SMAUG algorithm was run on the raw linked trajectory position data from CDC-42 wild-type variant to identify the number of mobility states. Iterations were started with 10 mobility states, 15 ms frame time and 100 nm pixel size, the rest of parameters were kept as default.^83^

Simulated data was generated by simulating image stacks of Gaussian approximated point spread functions with the same intensity (75 kcounts), background (17 kcounts) and noise (250 counts standard deviation) as found in real TIRF images of GFP::CDC42 in live embryos. Brownian motion with diffusion coefficient of 0.65μm^2^/s in trajectories was simulated and image stacks tracked with the same Matlab algorithm as for real image data.

### sc-SiMPull data anaylsis

The raw TIRF movies from sc-SiMPull experiments were processed using open-source software written in MATLAB and available at https://github.com/Dickinson-lab/SiMPull-Analysis-Software.^35^ The software uses a sliding window subtraction approach followed by image segmentation to identify single bait protein capture events. Each bait protein capture event is checked for presence of a corresponding prey protein signal; simultaneous binding of bait and prey proteins to the coverslip (termed co-appearance) indicates that the proteins are in complex. After identifying GFP::CDC-42 / HaloTag::aPKC complexes in this way, the dwell time of the HaloTag signal (i.e., the time from its appearance to its disappearance) was measured. The distribution of dwell times was converted to a disappearance rate constant k_disappear_ using a Bayesian inference approach that yields both the maximum likelihood value for k_disappear_ and its 95% credible interval.^120^ k_disappear_ is the sum of the dissociation rate constant k_off_ and the photobleaching rate constant k_bleach_. To correct for photobleaching and extract the value of k_off_, we measured k_bleach_ using a YFP::HaloTag fusion protein (for which k_off_ = 0).^35^ Each measurement of k_disappear_ was converted to a k_off_ using a splitting probability analysis^36^ and single-embryo measurements of k_off_ are plotted as grey circles in **Figure 2F**. Briefly, we used the value of k_bleach_ from a matched control experiment to calculate the number of molecular disappearance events that are expected due to photobleaching in each dataset. Disappearance events in excess of this expected value are inferred to be due to dissociation, and the frequency of these events is used to infer k_off_ and its 95% credible interval. The splitting probability approach is equivalent to calculating k_off_ by simple subtraction (k_off_ = k_disappear_ – k_bleach_), but is more rigorous because it allows us to account for error in the estimate of k_bleach_ (See Deutz et al.^36^ for details).

### Image analysis – Differences in PKC-3 vs PAR-3 domain size and in PAR-2 vs PAR-3 domain size

Analysis performed on midplane images of establishment (embryos staged between pronuclear touch and meet) and maintenance stage embryos (staged between pronuclear meet to metaphase). We used custom-built FIJI macro to create cortical flat-outs. aPAR retraction difference was determined by subtracting the length of PAR-3 domain from the total domain length of PKC-3 for each zygote. The difference in domain sizes (PKC-3 – PAR-3) is plotted in μm (**Figure 3C**). aPAR and pPAR gap was determined by subtracting the length of PAR-3 and PAR-2 domains from the total cortex length, giving a positive value in the presence of a gap or a negative value when the domains overlapped **(Figure 3I**).

### Image analysis – PKC-3 domain size in vivo

Embryos were staged at establishment (pronuclear migration) and maintenance (pronuclear meet, pronuclear centration/rotation and nucelar envelop breakdown) by position and size of the pronuclei (**Figure S5D**). At each of these stages a 5 frame Z projection of max intensity was created. Cortical flat-outs of these images were then created using a FIJI macro where the cortex was selected in a clockwise rotation starting (and finishing) at the anterior pole. The total HaloTag::aPKC domain was obtained to calculate the percentage of HaloTag::aPKC membrane occupancy.

### Image analysis - Analysis of AB/P1 PKC-3 normalised intensity

Midplane 2-cell images had their cortexes flattened using a FIJI macro. Cortexes were selected as either the outward facing AB or P1 membranes - neither included the membrane between the two cells. The flatouts then had a 5-pixel width line drawn along the length of the cortex to measure average intensity, a similar line was then used to determine the background outside of the embryo in the same image - avoiding any debris. The background subtracted cortical intensity of the AB cell was divided by that of the P1 cell to give an intensity ratio of HaloTag::aPKC (**Figure S5F**).

### Image analysis – Protein co-localisation

PKC-3 and PAR-3 co-localisation (**Figure 3E**) was studied on anterior cortical images of maintenance stage embryos – defined as pronuclear meet to metaphase. We used a custom-built FIJI macro to select an 11 x 11 μm area of anterior cortex free from debris. The pre-built FIJI plugin ComDet (v.0.0.5) (https://github.com/UU-cellbiology/ComDet) was used to detect particles above set thresholds (size – 4 pixels, intensity – 11) across all images to yield a readout of the levels of co-localisation between particles of PKC-3 and PAR-3. For the analyses of NMY-2 co-localisation with CDC-42 pS71 or with PAR-3 the BIOP JACoP plugin in FIJI was used (**Figure 4F**). A focused region at the anterior cortex was select and default co-localisation parameters (Otsu threshold) were used to obtain a Pearson coefficient value, a value of 1 indicating a perfect co-localisation.

### Image analysis – PKC-3 cortical Asymmetric Index (ASI)

Analysis performed on cortical images of maintenance stage embryos – defined as pronuclear meet to metaphase (**Figure 3F**). We used a custom-built FIJI macro to select three 5.5 x 5.5 μm areas in the image; one, the background outside the embryo, two, an area of anterior cortex, and three, an area of posterior cortex. Background subtraction was performed on the anterior and posterior areas. Mean pixel intensity for anterior (A) and posterior (P) areas were calculated and imputed into the ASI formula:

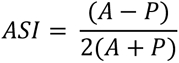

All ASI values were normalised by dividing each value by the mean ASI value of wild-type. After normalisation “1” depicts the average asymmetry observed in wild-type zygotes, above “1” indicates a stronger asymmetry than wild type and below “1” less asymmetry, with “0” indicating that the feature under study is not asymmetric.

### Image analysis – PKC-3 anterior cortical coefficient of variation (CV)

Analysis performed on cortical images of maintenance stage embryos – defined as pronuclear meet to metaphase (**Figure 3G**). Using the same background subtracted anterior selected areas as the ASI analysis, coefficient of variation was calculated as standard deviation divided by the mean pixel intensities;

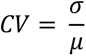

### Image analysis – NMY-2 coefficient of variation

Coefficient or variation was calculated (as above) on a 11 x 11 μm cortical area that was centred within the embryo to analyse a region where NMY-2 foci are present. In this area the mean intensity and standard deviation were background corrected using a 5.5 x 5.5 μm area outside the embryo. The average coefficient of variation was calculated for frames 10-80 (70 seconds) of establishment zygotes (starting at pronuclei touch). For representation purposes CV values were normalised by the average CV value in the control (wild type or GFP::CDC-42 strains) (**Figure 5D, 5G**).

### Image analysis – NMY-2 foci area

Foci thresholding was performed on establishment embryos using a custom-built FIJI macro. Briefly, the movie was cropped to frames 5-95 of its total length. The embryo was then selected using the polygon ROI to avoid any external debris. The user then determined the pixel intensity value threshold to select NMY-2 foci. Threshold was set to include the more diffuse accumulation of NMY2 preceding foci peak so the full range of NMY-2 foci size throughout its lifetime could be averaged. Area of the foci was obtained with the Analyse Particle tool from FIJI (lower foci size limit set to 50 pixel^2^) (**Figure 5C**).

### Image analysis – Determining the presence of CDC-42 pS71 and NMY-2 foci

The analyses of CDC-42 pS71 or NMY-2 foci presence (**Figure 4B, 4D, and Figure S6B**) was determined by 2D intensity correlation matrix of a squared area encompassing the anterior of each zygote (MATLAB script), followed by the identification of the first local minima by scanning the resultant matrix from the centre with and increasing circumference.^56^ The position of this local minima is used as a proxy of foci size and zygotes with average foci size >1.14µm were considered as positive. 2D intensity correlation results were compared to blindly sorting embryos with and without foci, confirming our approach.

### Image analysis – NMY-2 flow measurements

Actomyosin cortical flow velocity was determined by performing Particle Image Velocimetry (PIV) on the time-lapse movies using the freely available PIVlab MATLAB algorithm (pivlab.blogspot.de). A 2-step multi pass with linear window deformation and a final interrogation area of 32 pixels with a step size of 16 pixels were used as settings in the PIVlab code. The resulting 2D velocity fields obtained from PIVlab were then divided into 18 bins along the anteroposterior axis of the embryo with a stripe height of 60 pixels. To determine the posterior velocity, the x-component of the velocity in the posterior across bins 13 to 17 was then spatially averaged in each frame, before performing a temporal average from the start of flow until the start of pseudocleavage. This spatiotemporal average was then compared across the different conditions (**Figure 5E**, **5H**).

### Image analysis – Maintenance NMY-2 cortical intensity measurements

Analysis was performed on images of maintenance phase embryos – defined in the time between NMY-2 puncta disappearance and before nuclear envelope breakdown. 11 x 11 μm areas were selected in the centre of the anterior and posterior cortex. In these areas the mean intensity and standard deviation were background corrected using a 2.2 x 2.2 μm area outside the embryo. The average coefficient of variation was calculated for 50 frames (50 seconds). The value of the anterior was then divided by that of the posterior to give the plotted ratios (**Figure S7**).

### Embryo lethality

Embryo lethality percentage was calculated by estimating the number of eggs hatched at 15 °C from a 62 to 72-hour time-window (from L4 stage) from a progeny of 3 adult worms and following the formula: 100x larvae/(eggs+larvae). Two independent experiments in duplicate were performed.

### Statistical analysis

For most analysis, significance was assessed using an unpaired, two-tail Student’s T test unless otherwise noted, with the following criteria: *p<0.05, **p<0.01, ***p<0.001, ****p<0.0001. Chi-square test was used to determine differences in the number of zygotes with pS71 or NMY-2 foci, with the following criteria: *p<0.05, **p<0.01, ***p<0.001.

For sc-SiMPull data, a single maximum likelihood and credible interval for k_off_ in each genetic background was calculated by pooling the data from all experiments and is shown by coloured bars in **Figure 2F**. To determine the upper and lower bounds of the confidence interval, the calculation was repeated using the lower and upper bounds (respectively) of the credible interval for k_bleach_, thereby ensuring that errors in the determination of k_bleach_ are reflected in the credible interval for k_off_. k_off_ estimates from sc-SiMPull data are considered significantly different from one another if their 95% credible intervals do not overlap.

For HILO data (**Figure 1H)**, D is extracted from gamma fitting and error measured from 95% confidence intervals to the fit. D estimates are considered significantly different from one another if their 95% credible intervals do not overlap.

All other data are presented as mean values together with all data points, mean values with indicated (n), mean ± 95% confidence interval (CI) and all data points, mean ± standard error of the mean (SEM) and all data points, mean ± standard deviation (SD) and all data points, or box plots showing all data points, the median and interquartile range (IQR) with whiskers extending to the max and min values.

### Limitations section

We are mindful of the limitations of studying phosphomutants, as they cannot perfectly mimic the phosphorylated or de-phosphorylated states. One reason for this is that, in wild-type, these states are constantly interchanging, while the phosphomutants are stalled in a given phospho-state. We have been cautious when interpreting the phenotypes of CDC-42 phosphomutants and indicated in the text an instance where we found the phosphomimetic mutant of CDC-42 not mimicking the localisation of CDC-42 pS71. We do not observe CDC-42(S71E) enriched at cortical foci, where we can detect CDC-42 pS71 by immunostaining. However, CDC-42(S71E) leads to actomyosin foci dispersion suggesting that even if not enriched at these foci, CDC-42(S71E) might still have an effect on them. Moreover, CDC-42(S71E) is not simply a loss-of-function mutant, as CDC-42 (RNAi) depletion presents the opposite phenotype (tight NMY-2 foci). Overall, the phosphomimetic and non-phosphorylatable mutants of CDC-42 display opposite phenotypes, as expected, supporting the study of these mutants to suggest possible ways in which CDC-42 phosphorylation could be regulating CDC-42 function and its interaction with other proteins.

In our *in vitro* kinase assay in **Figure 1C** we detect a background signal in absence of ATPgS. We cannot explain this background, but after background correction our results still indicate that aPKC phosphorylates CDC-42 *in vitro*. This outcome is further supported by the *in vitro* kinase assay using purified human aPKC where we do not observe this background (**Figure 1A**).

The findings on CDC-42 phosphorylation regulating actomyosin cortex are very novel (**Figure 5** and **Figure S7**), and more work is required to fully understand mechanistically how CDC-42 phosphorylation is having this role. For example, it is hard to disentangle if CDC-42 phosphorylation is mostly acting upstream of aPKC (controlling its activity) or also downstream of aPKC, for example via MRCK-1. Overall, more work is needed to understand the complicated and interconnected network controlling actomyosin organization in the zygote.

## Supporting information

Key Resource table

## Acknowledgements

We thank Nathan Goehring (The Francis Crick Institute) and Jonathan Higgins (Newcastle University) for helpful discussions and comments on the manuscript. We also thank Luke Lavis for JaneliaFluor dyes; Eva Zeiser and Julie Ahringer for guidance on the generation of the MosSCI lines; and Artur Ribeiro Fernandes and Daniel St Johnston for help on the generation of MBP recombinant proteins. This work was supported by BBSRC Research Grant BB/R019436/1 (JR, JP), BBSRC PhD studentships (IS, AB), a PhD studentship from Newcastle University (AG), an Undergraduate Research Fellowship from UT Austin (LND), and NIH R01 GM138443 (DJD) and NSF MCB 2237451 (DJD). DJD is a CPRIT scholar supported by the Cancer Prevention and Research Institute of Texas (RR170054). SRN acknowledges funding support from the Department of Atomic Energy (DAE), Govt. of India (Project Identification no. RTI4003, DAE OM no. 1303/2/2019/R&D-II/DAE/2079 dated 11.02.2020). Academy of Medical Sciences Springboard Award (SBF007\100046) (AW). SN acknowledges Cancer Research UK (CC2119, CC2068), the UK Medical Research Council (CC2119, CC2068), and the Wellcome Trust (CC2119, CC2068). We also would like to acknowledge The Newcastle University Bioimaging Facility.

## Author Contributions

JR conceived the project, supervised the work and secured funding. JP, AG, AB, IS, CS performed the experiments and analysed the data. LND performed the sc-SiMPull experiments, and LND and DJD analysed the sc-SiMPull data. AW supervised JP in the acquisition and analyses of TIRF data. SRN provided MATLAB scripts and supported NMY-2 video analyses. SE purified the aPKC iota - Par-6 complex. JR wrote the manuscript, all authors discussed and contributed to the final version.

## Declaration of interest

The authors declare no competing interest

---

**Figure S1.**
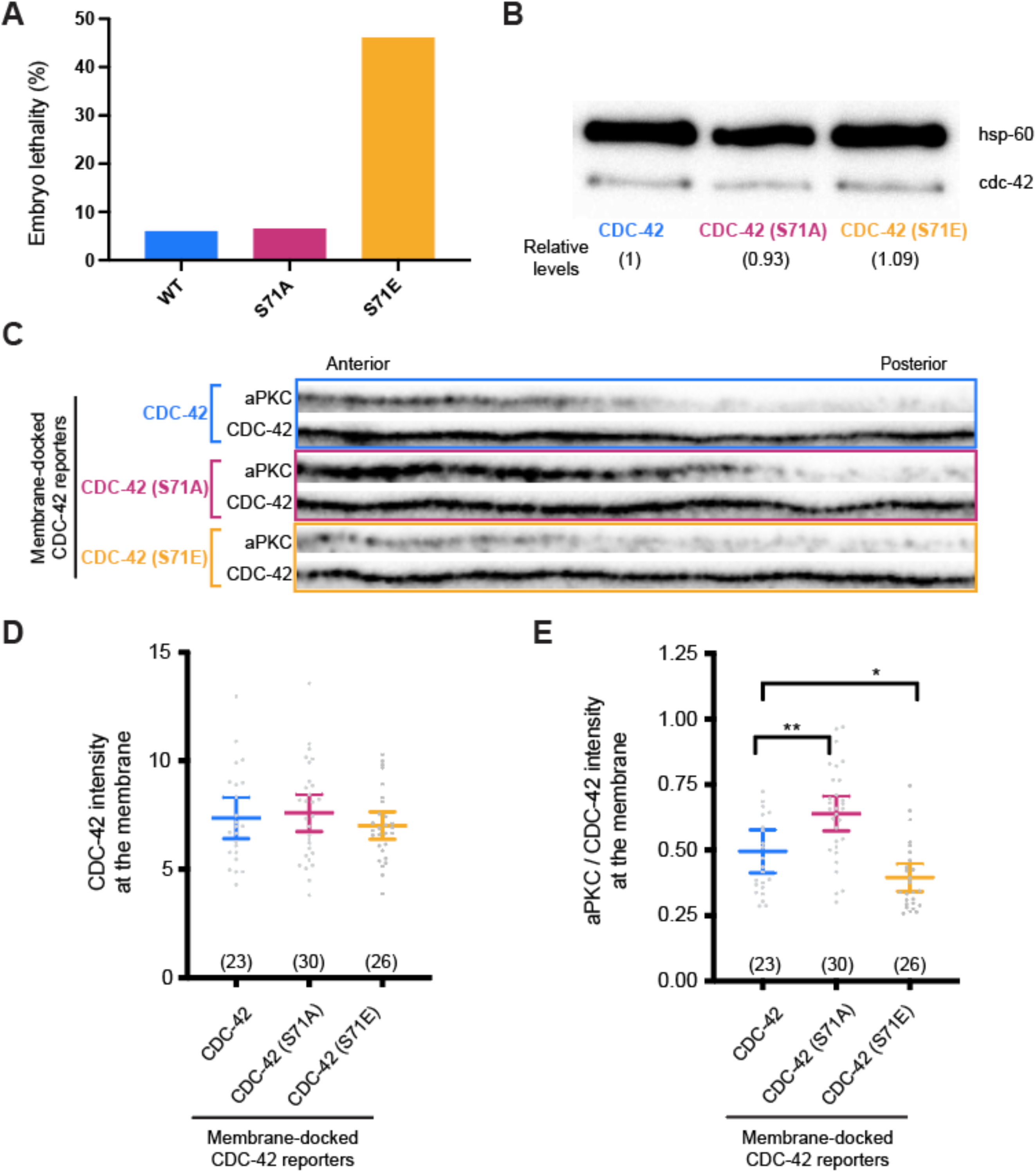
Characterisation of CDC-42 phosphomutant strains. **A.** Mean embryonic lethality observed for GFP::CDC-42, GFP::CDC-42(S71A) and GFP::CDC-42(S71E) strains (endogenous CDC-42 is depleted in all CDC-42 reporters) in two independent experiment performed in duplicate. **B.** Immunoblot of embryo extracts showing the protein levels for GFP::CDC-42, GFP::CDC-42(S71A) and GFP::CDC-42(S71E) constructs detected with anti-GFP. Hsp-60 was used as a loading control. Numbers are the average intensity of CDC-42 in the different strains (loading-corrected) shown relative to GFP::CDC-42 intensity (mean from three blots). **C.** Representative flattened-out membranes at the transition from the anterior to the posterior domain in zygotes where the different CDC-42 constructs have been artificially recruited to the membrane (membrane-docked). Profiles are centred on 50% of the zygote length and 100 pixels shown in either direction. aPKC (immunofluorescent) and CDC-42 (GFP detection) are shown for the indicated strains during polarity maintenance. Note how GFP::CDC-42, GFP::CDC-42(S71A) and GFP::CDC-42(S71E) are observed throughout the membrane whereas aPKC is still enriched to the anterior side of the zygote. **D.** Measurement of CDC-42 levels in the anterior membrane (measured in a 60 x 60 pixel area from a straightened anterior cortex) of polarity maintenance zygotes. Anterior membrane intensity is normalised with the corresponding CDC-42 cytoplasmic level (mean ± CI 95%). No significant differences are observed between the different strains. **E.** Measurement of aPKC intensity at the anterior membrane normalised to the observed CDC-42 intensity (mean ± CI 95%). After CDC-42 normalisation there is still a significant difference in the levels of anterior aPKC observed for GFP::CDC-42(S71A) and GFP::CDC-42(S71E) when compared to GFP::CDC-42 wild type. In C-E endogenous CDC-42 is depleted in all strains, hence aPKC recruitment is dependent on the ectopically expressed CDC-42 constructs. Unpaired, two-tail Student’s T test *p<0.05, **p<0.01.

**Figure S2.**
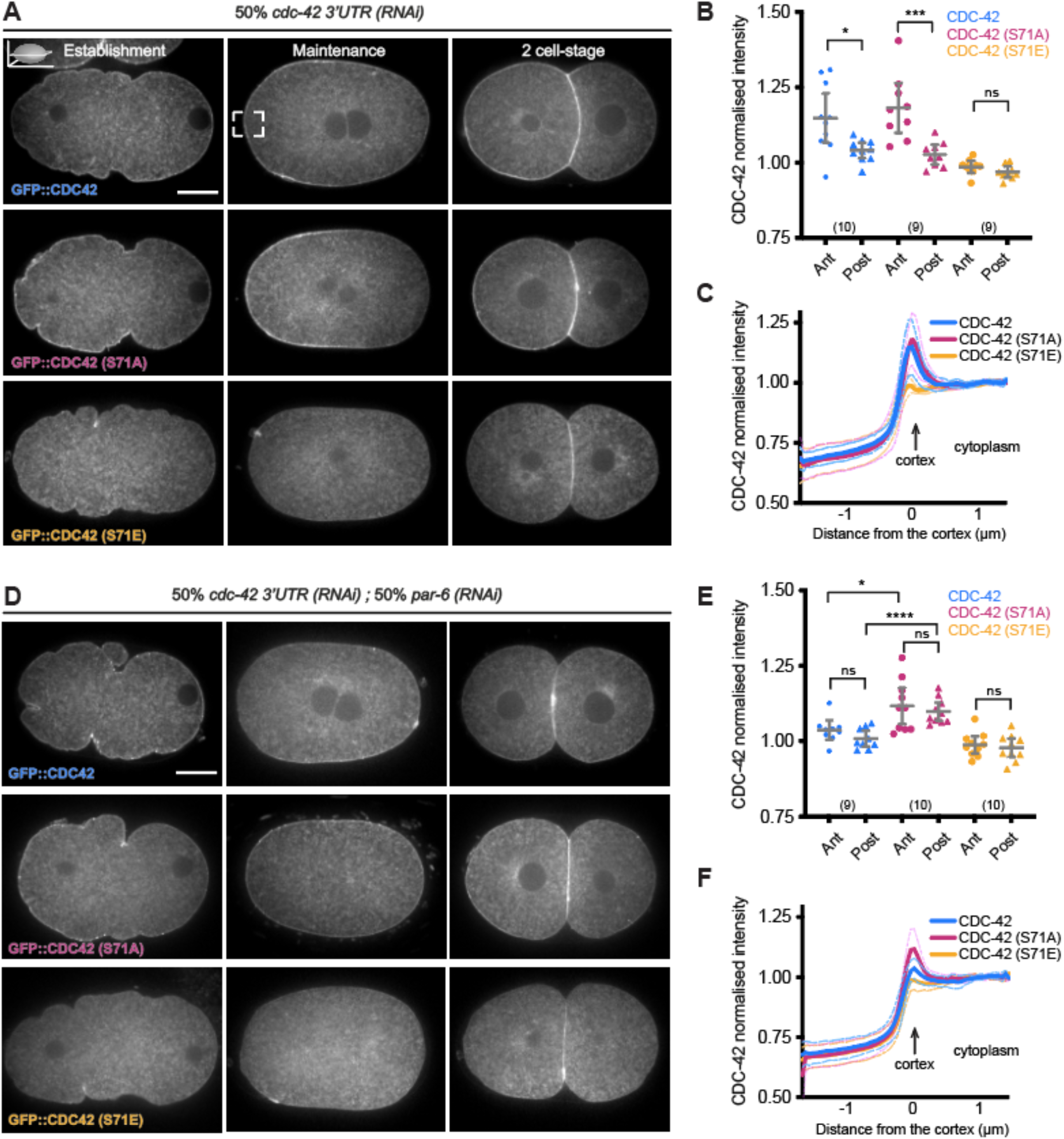
PAR-6 is required for CDC-42 asymmetry and membrane stabilisation. **A.** Representative midsection confocal images of live embryos at establishment, maintenance and 2-cell stage showing GFP::CDC-42, GFP::CDC-42(S71A) and GFP::CDC-42(S71E). In these CDC-42 reporter lines we partially depleted endogenous CDC-42 by 50% RNAi of its 3’ UTR (50% dilution of RNAi bacteria, see methods for more details). Note that the phenotypes observed in this RNAi condition are very similar to those observed when we deplete CDC-42 by 100% RNAi of its 3’UTR (no RNAi bacteria dilution) (Fig. 1D-F) **B.** CDC-42 anterior and posterior cortical intensities (values are normalised by their corresponding cytoplasmic CDC-42 levels) observed in CDC-42, CDC-42(S71A) and CDC-42(S71E) maintenance stage embryos (mean ± CI 95%). **C.** CDC-42 intensity profiles spanning the anterior membrane of CDC-42, CDC-42(S71A) and CDC-42(S71E) maintenance stage zygotes, showing mean ± SD. Briefly a 60 x 60 pixel area from a straightened anterior cortex (see inset in A in GFP::CDC-42 maintenance) was projected in the y-axis to give a cross section profile spanning the cortex/membrane. The values are normalised so that the cytoplasmic levels are set to 1. **D. E. F.** show the zygotes and analyses of the GFP::CDC-42, GFP::CDC-42(S71A) and GFP::CDC-42(S71E) strains upon 50% RNAi depletion of both, PAR-6 and endogenous CDC-42 (see methods). Note that in this condition GFP::CDC-42 and GFP::CDC-42(S71A) lose their asymmetric localisation (E) just like GFP::CDC-42(S71E) upon depletion of only endogenous CDC-42 (B). However, GFP::CDC-42(S71A) retains some of its membrane localisation upon depletion of PAR-6 unlike GFP::CDC-42 (E,F). Unpaired, two-tail Student’s T test *p<0.05, ***p<0.001, ****p<0.0001, ns not significant. Scale bar in establishment zygotes: 10 µm.

**Figure S3.**
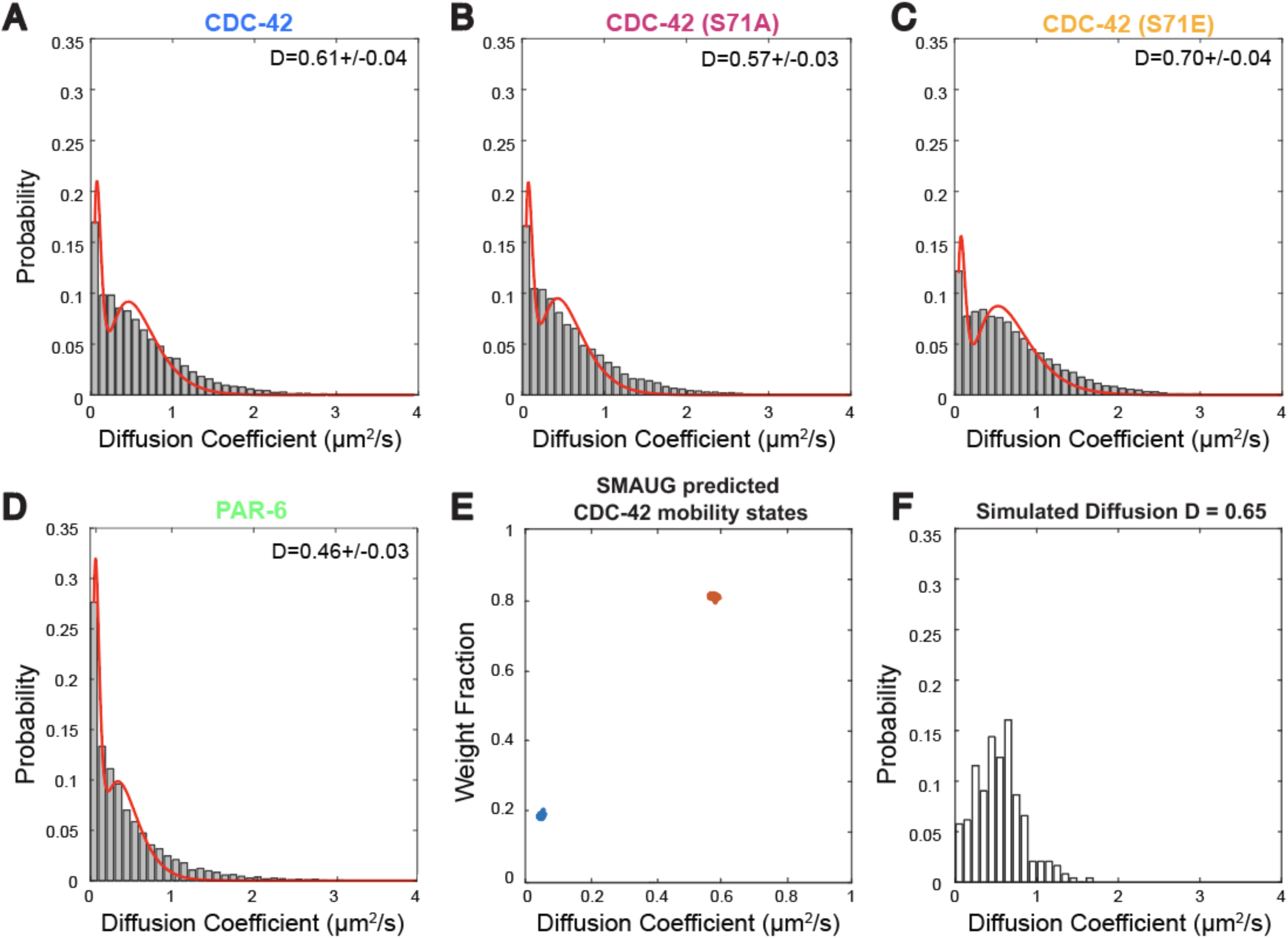
Phosphorylation of CDC-42 accelerates its movement within the membrane. Distribution of diffusion coefficients for GFP::CDC-42, GFP::CDC-42(S71A), GFP::CDC-42(S71E), GFP::PAR-6 and simulated data. **A.** to **D.** Double Gamma fits (red curve) to each distribution captures two mobility states for each CDC-42 variant and for PAR-6, mobile (D> 0.1 μm^2^/s) and immobile (D< 0.1 μm^2^/s). Diffusion coefficient of the mobile fraction indicated above each graph (data also shown in Fig. 1H). **E.** We further confirmed two states in our CDC-42 diffusion data using nonparametric Bayesian statistics with SMAUG algorithm, which suggested ∼20% CDC-42 particles were in an immobile state (blue cluster, D∼0.05μm^2^/s), with the remaining mobile population diffusing with D∼0.6μm^2^/s (red cluster), agreeing well with our fitting. **F.** Simulation is done based on real data from the mobile fraction of CDC-42 considering particles moving at 0.65μm^2^/s (see methods for more details). From the simulation we extract the baseline fraction of apparently immobile particles (0.06) obtained due to inherent errors in fitting mean squared displacement data. We show this baseline in Fig. 1I.

**Figure S4.**
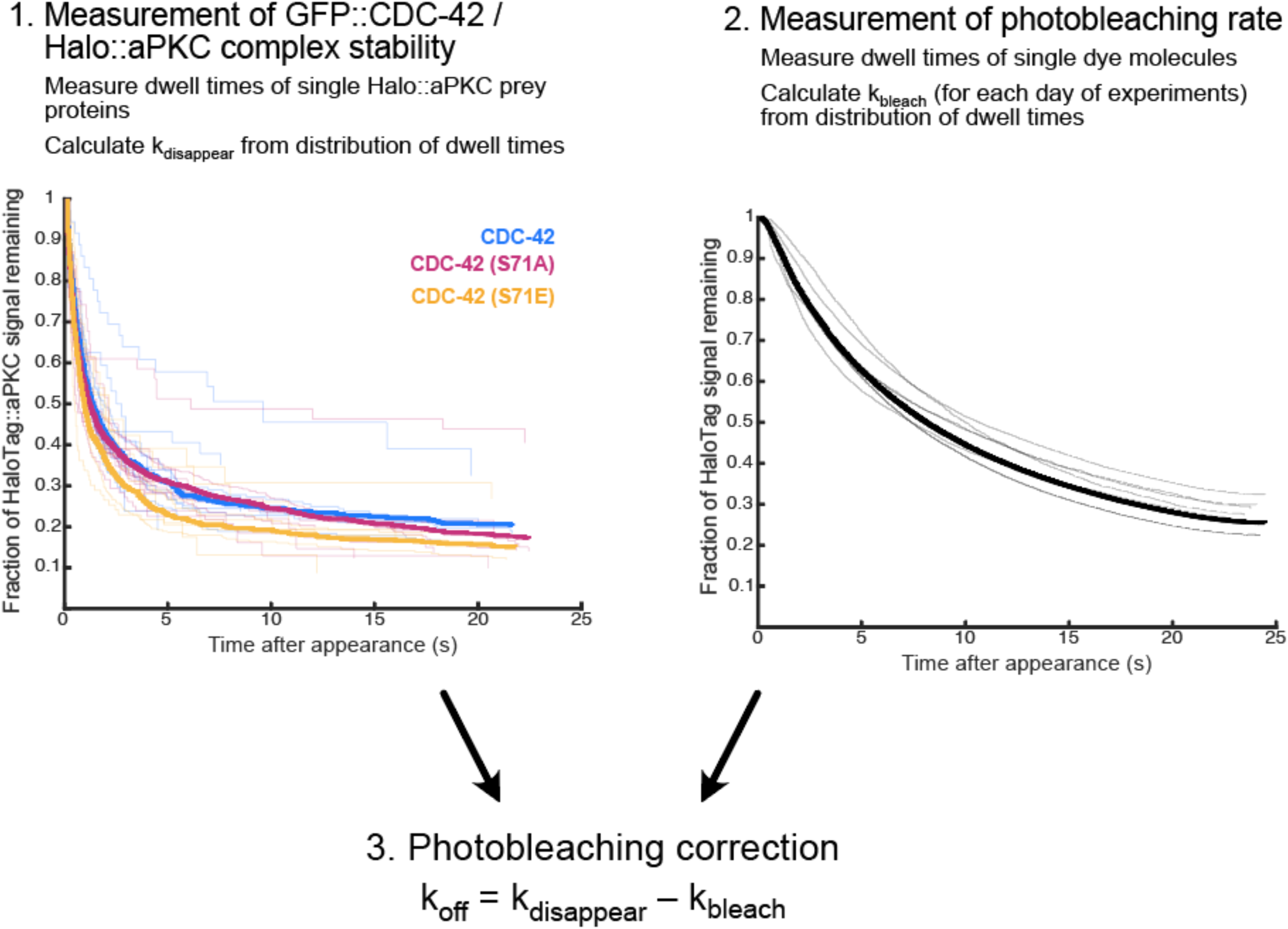
Calculation of koff for the CDC-42/aPKC interaction from HaloTag::aPKC signal disappearance. For each identified GFP::CDC-42/HaloTag::aPKC complex, we measure the survival time (i.e., the dwell time) of the HaloTag::aPKC prey protein signal (see Fig. 2E for an illustration of the approach). Each dwell time is the interval between GFP::CDC-42/HaloTag::aPKC complex capture and HaloTag::aPKC signal disappearance. The distribution of dwell times for the population is represented as a Kaplan-Meier survival curve (step 1), which illustrates the probability of the HaloTag::aPKC signal remaining present as a function of time since capture. In the plot, thin curves represent single-embryo experiments, and the bold curve represents the sum of all data (obtained by pooling all single-molecule observations regardless of which embryo they originated from). We use the distribution of dwell times to calculate the apparent rate constant kdisappear for each single-embryo experiment. Since each HaloTag::aPKC molecule can disappear due to either unbinding or photobleaching, we separately measured the photobleaching rate of the JF646 HaloTag ligand using a YFP::HaloTag transgenic control strain (step 2). The photobleaching rate constant kbleach is calculated from these data. There is some variability in the photobleaching rate over time, due to fluctuations in laser power and/or microscope alignment, and so each experimental measurement is corrected separately for photobleaching using a paired control measurement captured the same week (step 3). The corrected photobleaching rates are pooled across multiple embryos to obtain a maximum probability estimate of koff, and its 95% credible interval, for each experimental condition. See Deutz et al. 2023 and Methods for details of this calculation.

**Figure S5.**
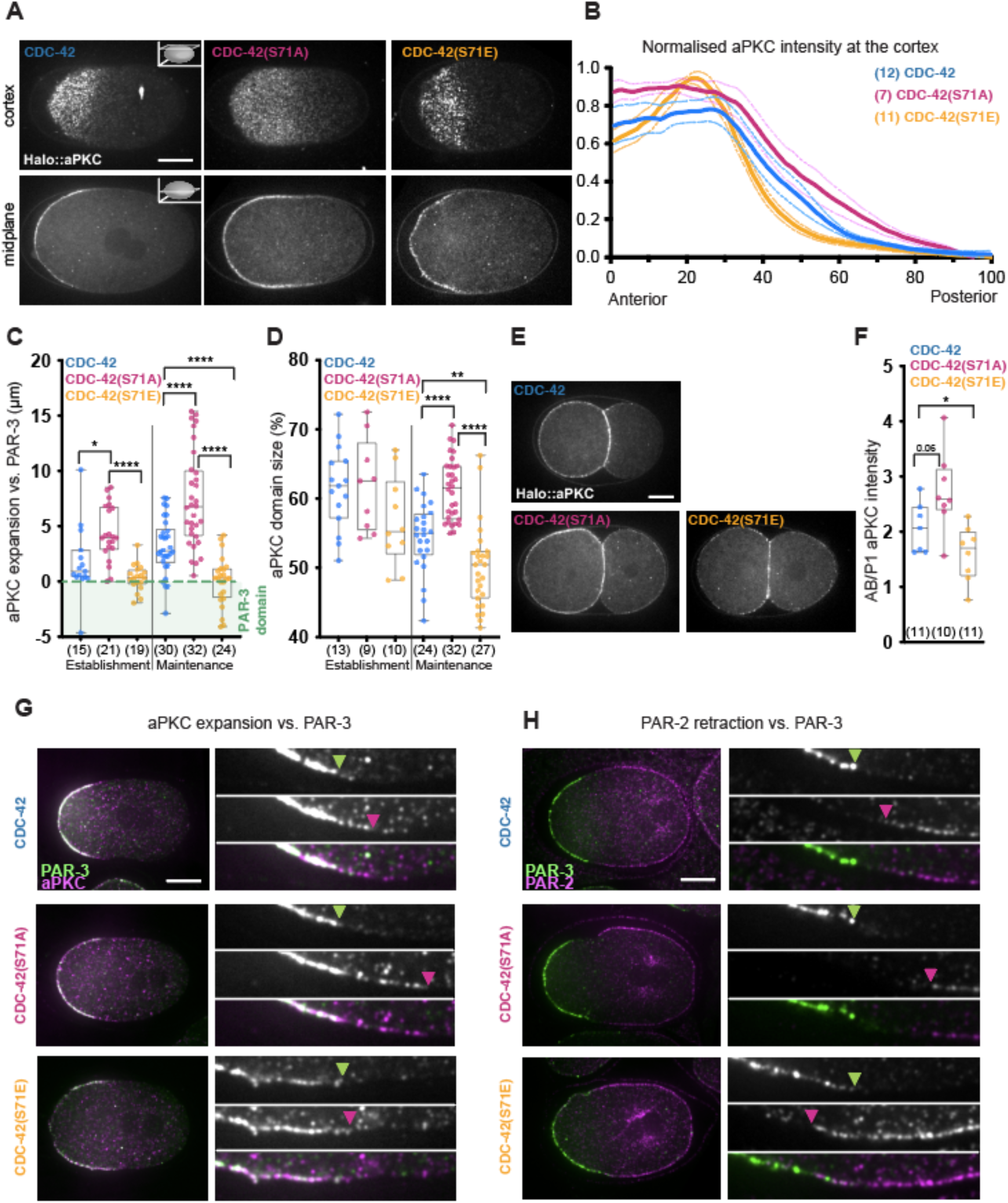
Mimicking or impeding CDC-42 phosphorylation on S71 leads *in vivo* to aPKC domain alteration that resemble those observed in immunostained zygotes. **A.** Representative cortical and midplane confocal images of HaloTag::aPKC in CDC-42, CDC-42(S71A) and CDC-42(S71E) live zygotes (endogenous CDC-42 is depleted in all CDC-42 reporters). **B.** Plot profile of normalised aPKC intensity (mean ± SEM) extracted from cortical sections of polarity maintenance zygotes (anterior to posterior, see methods), showing a slower decline of aPKC levels in the CDC-42(S71A) mutant and a faster one in the CDC-42(S71E) when compared to the control. **C.** Quantification of aPKC expansion vs. PAR-3 (midplane images) in immunostained polarity establishment and maintenance stage zygotes. The maintenance stage data is also shown in Fig. 3C. **D.** aPKC domain size measurements from HaloTag::aPKC live zygotes (midplane sections) during polarity establishment and maintenance. **E.** Two-cell stage representative midplane confocal images of Halo-labelled aPKC in CDC-42, CDC-42(S71A) and CDC-42(S71E) zygotes (endogenous CDC-42 is depleted in all CDC-42 reporters). **F.** Quantification of aPKC membrane enrichment in the anterior daughter cell (AB) vs. the posterior one (P1). **G.** and **H.** Representative midplane confocal images of co-immunostained PAR-3/aPKC (G) and PAR-3/PAR-2 (H) polarity maintenance stage CDC-42, CDC-42(S71A) and CDC-42(S71E) zygotes (endogenous CDC-42 is depleted in all CDC-42 reporters). Insets are zoomed regions at the end of the PAR-3 domain showing the corresponding aPKC or PAR-2 localisation. All box plots show the median ± IQR and all data points. Unpaired, two-tail Student’s T test *p<0.05 **p<0.01, ****p<0.0001. Scale bar: 10 µm.

**Figure S6.**
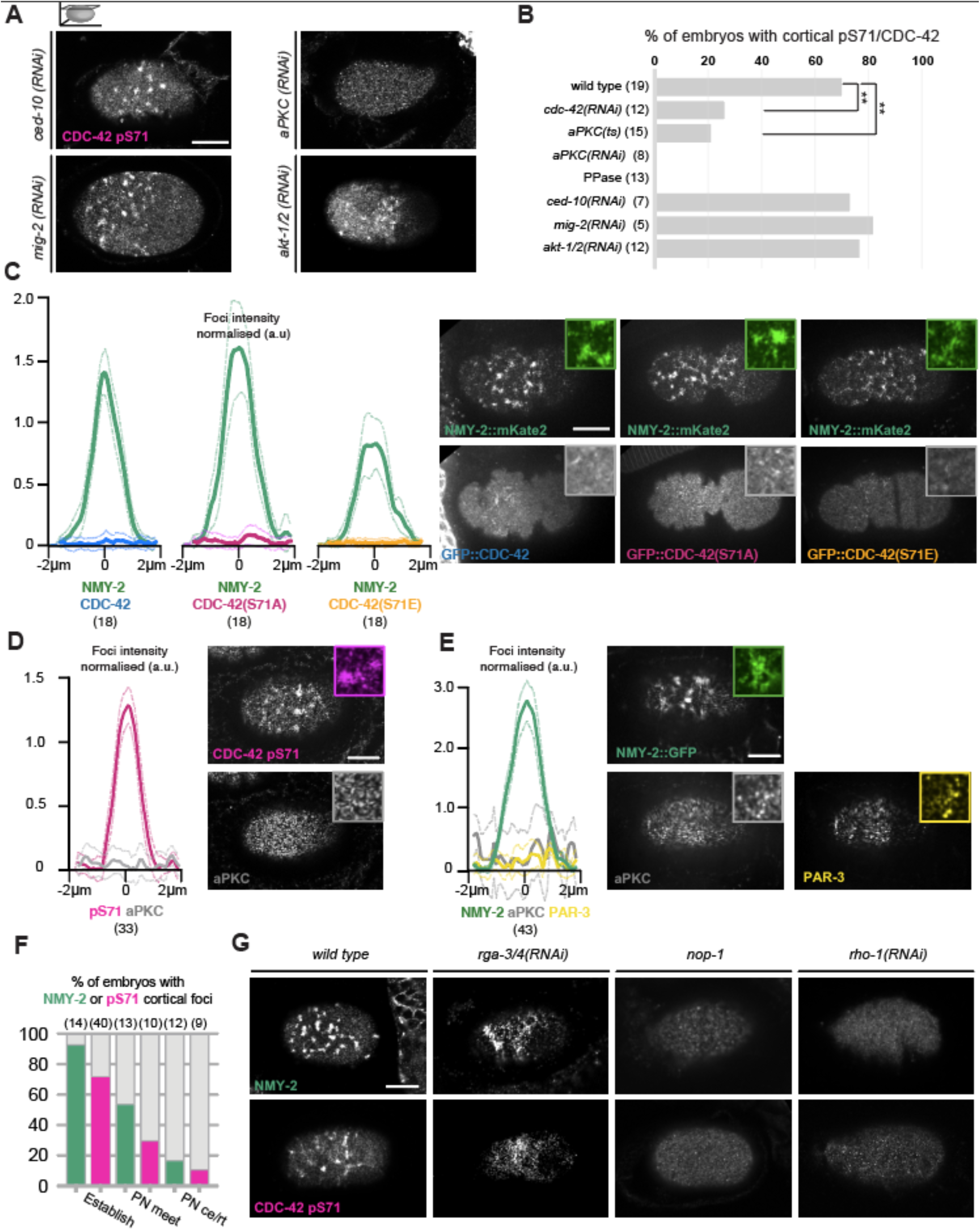
CDC-42 pS71 cortical foci do not depend on Rac1 small GTPases or Akt1 kinase but do depend on the actomyosin organisation. aPARs and CDC-42 variants are not observed at cortical foci. **A.** Immunofluorescent detection of CDC-42 phosphorylation on serine 71 in polarity establishment zygotes upon depletion of Rac1 homologues (CED-10 and MIG-1) and depletion of aPKC or AKT-1 and AKT-2 kinases (*akt-1/2(RNAi)*). **B.** Quantification showing the percentage of zygotes with CDC-42 pS71 cortical foci inferred from 2D intensity correlation analysis (see methods). For ease of comparison the graph contains data presented in main Fig. 4B. **C.** Plot profiles of NMY-2 and CDC-42 intensities (mean ± SEM) across cortical foci in the different CDC-42 variant strains (endogenous CDC-42 is depleted in all CDC-42 reporters). Representative confocal cortical images and insets containing zoomed in cortical regions of the CDC-42 variant strains at polarity establishment. **D.** Plot profiles of CDC-42 pS71 and aPKC intensities (mean ± SEM) across cortical foci in wild-type zygotes. Representative confocal cortical images of CDC-42 pS71 and aPKC co-immunostained wild-type zygotes during polarity establishment. **E.** Plot profiles of NMY-2, aPKC and PAR-3 intensities (mean ± SEM) across cortical foci from wild-type zygotes. Representative confocal cortical images of aPKC and PAR-3 co-immunostained in NMY-2::GFP zygotes during polarity establishment. **F.** Quantification of zygotes with NMY-2 or CDC-42 pS71 foci during polarity establishment and at the later polarity maintenance stages of pronuclei meet and pronuclei centration/rotation. The number of zygotes with NMY-2 and pS71 foci drops similarly over-time. **G.** Representative cortical confocal images of wild-type, *nop-1*, *rga-3/4* (*rga-3 and rga-4)* (*RNAi*) or *rho-1(RNAi)* depleted embryos at polarity establishment, stained for NMY-2 or CDC-42 pS71. We observed that hyper activation of the RHO-1 pathway, via simultaneous downregulation of the GAPs, RGA-3 and RGA-4, led to expanded and disorganised CDC-42 pS71 foci similar to those reported for NMY-2 (Michaux et al., 2018; Schmutz et al., 2007; Schonegg et al., 2007; Tse et al., 2012). In *nop-1* mutant or *rho-1* partial depletion (10% RNAi, see methods) where NMY-2 foci are not observed during polarity establishment (Fievet et al., 2013; Tse et al., 2012), we also did not detect CDC-42 pS71 foci. Chi-square test **p<0.01. Scale bar: 10 µm.

**Figure S7.**
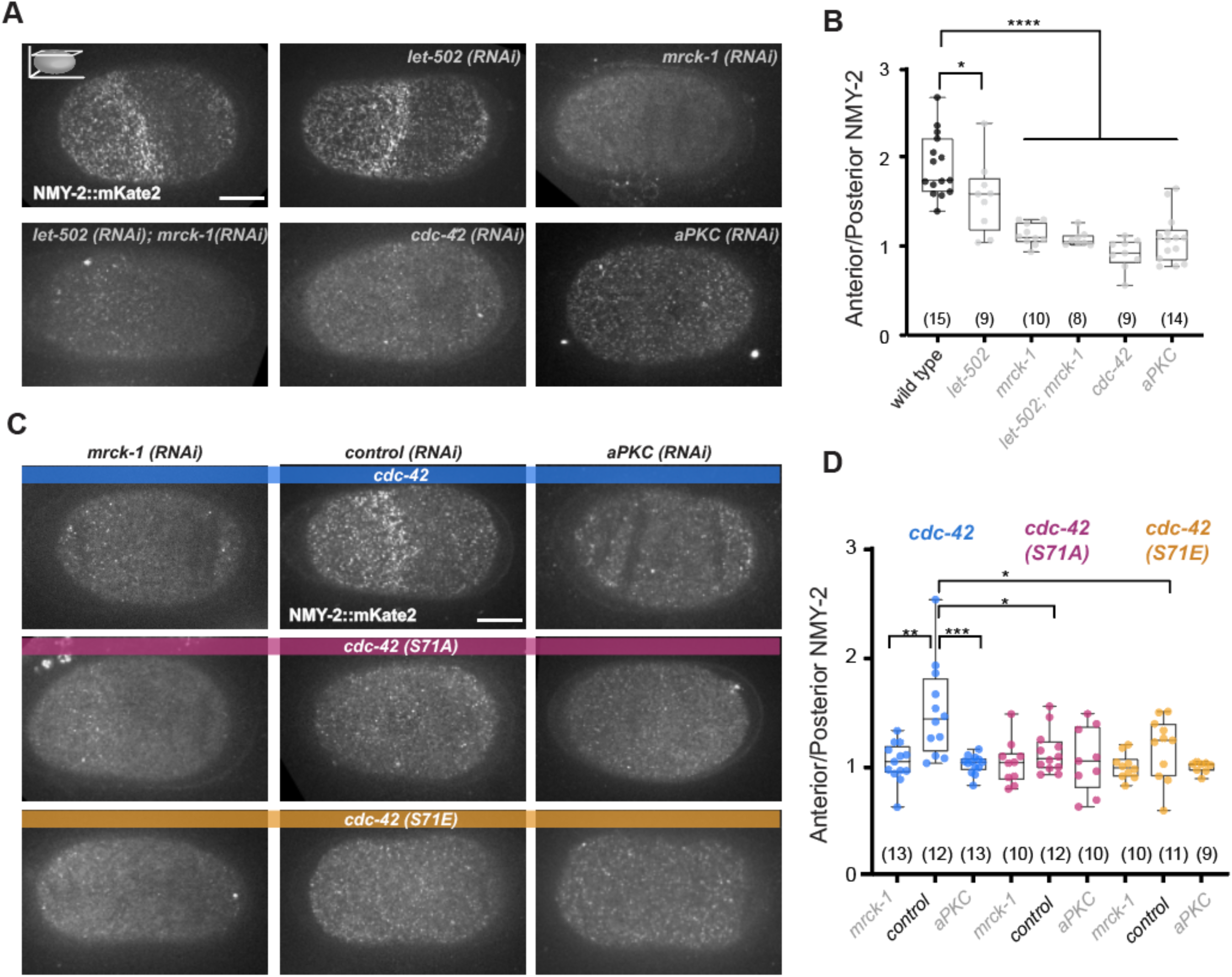
Turnover of CDC-42 phosphorylation is required for the formation of the anterior NMY-2 cap at polarity maintenance. **A.** Representative cortical confocal images of NMY-2 in wild-type, *let-502*, *mrck-1*, double *let-502/mrck-1* (1:1 RNAi bacteria mix, see methods for more details), *cdc-42* or *aPKC (RNAi)* depleted zygotes at polarity maintenance (prior to cleavage furrow). **B.** The presence of an anterior NMY-2 cap was determined by plotting NMY-2 CV at the anterior half of the zygote normalised by its CV at the posterior (a value of 1 indicating no NMY-2 cap presence). **C.** Cortical confocal images of NMY-2 in polarity maintenance CDC-42, CDC-42(S71A) and CDC-42(S71E) zygotes in control, *mrck-1(RNAi)* or *aPKC (RNAi)* conditions. Here endogenous CDC-42 is depleted by 50% RNAi of its 3’ UTR (50% dilution of RNAi bacteria) to be able to combine with 50% control (*RNAi)*, *mrck-1 (RNAi)* or *aPKC (RNAi)* (see methods). **D.** Plot analysing the anterior enrichment of NMY-2 (NMY-2 cap presence) as done in (B) for the conditions shown in C. All box plots show median ± IQR and all data points. Unpaired, two-tail Student’s T test *p<0.05, **p<0.01, ***p<0.001, ****p<0.0001, ns not significant. Scale bar: 10 µm.

## Notes

### Competing Interest Statement

The authors have declared no competing interest.

### Summary of Updates

We have edited the previous manuscript and added new data

## Bibliography

1. Campanale, J.P., Sun, T.Y., and Montell, D.J. (2017). Development and dynamics of cell polarity at a glance. J Cell Sci 130, 1201–1207. 10.1242/jcs.188599.

2. Goldstein, B., and Macara, I.G. (2007). The PAR Proteins: Fundamental Players in Animal Cell Polarization. Dev Cell 13, 609–622. 10.1016/j.devcel.2007.10.007.

3. St Johnston, D., and Ahringer, J. (2010). Cell polarity in eggs and epithelia: Parallels and diversity. Cell 141, 757–774. 10.1016/j.cell.2010.05.011.

4. Kemphues, K.J., Priess, J.R., Morton, D.G., and Cheng, N.S. (1988). Identification of genes required for cytoplasmic localization in early C. elegans embryos. Cell 52, 311– 320.

5. Watts, J.L., Etemad-Moghadam, B., Guo, S., Boyd, L., Draper, B.W., Mello, C.C., Priess, J.R., and Kemphues, K.J. (1996). par-6, a gene involved in the establishment of asymmetry in early C. elegans embryos, mediates the asymmetric localization of PAR-3. Development 122, 3133–3140.

6. Tabuse, Y., Izumi, Y., Piano, F., Kemphues, K.J., Miwa, J., and Ohno, S. (1998). Atypical protein kinase C cooperates with PAR-3 to establish embryonic polarity in Caenorhabditis elegans. Development 125, 3607–3614.

7. Rodriguez-Boulan, E., and Macara, I.G. (2014). Organization and execution of the epithelial polarity programme. Nat Rev Mol Cell Biol 15, 225–242. 10.1038/nrm3775.

8. Riga, A., Castiglioni, V.G., and Boxem, M. (2020). New insights into apical-basal polarization in epithelia. Curr Opin Cell Biol 62, 1–8. 10.1016/j.ceb.2019.07.017.

9. Mayor, R., and Etienne-Manneville, S. (2016). The front and rear of collective cell migration. Nat Rev Mol Cell Biol 17, 97–109. 10.1038/nrm.2015.14.

10. Petrie, R.J., Doyle, A.D., and Yamada, K.M. (2009). Random versus directionally persistent cell migration. Nat Rev Mol Cell Biol 10, 538–549. 10.1038/nrm2729.

11. Homem, C.C.F., and Knoblich, J.A. (2012). Drosophila neuroblasts: A model for stem cell biology. Development (Cambridge) 139, 4297–4310. 10.1242/dev.080515.

12. Gallaud, E., Pham, T., and Cabernard, C. (2017). Drosophila melanogaster Neuroblasts: A Model for Asymmetric Stem Cell Divisions. In, pp. 183–210. 10.1007/978-3-319-53150-2_8.

13. Hong, Y. (2018). aPKC: the Kinase that Phosphorylates Cell Polarity. F1000Res *7*, 903. 10.12688/f1000research.14427.1.

14. Achilleos, A., Wehman, A.M., and Nance, J. (2010). PAR-3 mediates the initial clustering and apical localization of junction and polarity proteins during C. elegans intestinal epithelial cell polarization. Development 137, 1833–1842.

15. David, D.J.V., Wang, Q., Feng, J.J., and Harris, T.J.C. (2013). Bazooka inhibits aPKC to limit antagonism of actomyosin networks during amnioserosa apical constriction. Development (Cambridge) 140, 4719–4729. 10.1242/dev.098491.

16. Graybill, C., Wee, B., Atwood, S.X., and Prehoda, K.E. (2012). Partitioning-defective protein 6 (Par-6) activates atypical protein kinase C (aPKC) by pseudosubstrate displacement. J Biol Chem 287, 21003–21011.

17. Lin, D., Edwards, A.S., Fawcett, J.P., Mbamalu, G., Scott, J.D., and Pawson, T. (2000). A mammalian PAR-3-PAR-6 complex implicated in Cdc42/Rac1 and aPKC signalling and cell polarity. Nat Cell Biol 2, 540–547.

18. McCaffrey, L.M., and Macara, I.G. (2009). The Par3/aPKC interaction is essential for end bud remodeling and progenitor differentiation during mammary gland morphogenesis. Genes Dev 23, 1450–1460.

19. Wirtz-Peitz, F., Nishimura, T., and Knoblich, J.A. (2008). Linking Cell Cycle to Asymmetric Division: Aurora-A Phosphorylates the Par Complex to Regulate Numb Localization. Cell 135, 161–173.

20. Soriano, E. V, Ivanova, M.E., Fletcher, G., Riou, P., Knowles, P.P., Barnouin, K., Purkiss, A., Kostelecky, B., Saiu, P., Linch, M., et al. (2016). aPKC Inhibition by Par3 CR3 Flanking Regions Controls Substrate Access and Underpins Apical-Junctional Polarization. Dev Cell 38, 384–398.

21. Rodriguez, J., Peglion, F., Martin, J., Hubatsch, L., Reich, J., Hirani, N., Gubieda, A.G., Roffey, J., Fernandes, A.R., St Johnston, D., et al. (2017). aPKC Cycles between Functionally Distinct PAR Protein Assemblies to Drive Cell Polarity. Dev Cell 42, 400–415.e9. 10.1016/j.devcel.2017.07.007.

22. Atwood, S.X., Chabu, C., Penkert, R.R., Doe, C.Q., and Prehoda, K.E. (2007). Cdc42 acts downstream of Bazooka to regulate neuroblast polarity through Par-6–aPKC. J Cell Sci 120, 3200–3206. 10.1242/jcs.014902.

23. Yamanaka, T., Horikoshi, Y., Suzuki, A., Sugiyama, Y., Kitamura, K., Maniwa, R., Nagai, Y., Yamashita, A., Hirose, T., Ishikawa, H., et al. (2001). PAR-6 regulates aPKC activity in a novel way and mediates cell-cell contact-induces formation of the epithelial junctional complex. Genes to Cells 6, 721–731. 10.1046/j.1365-2443.2001.00453.x.

24. Holly, R.W., and Prehoda, K.E. (2019). Phosphorylation of Par-3 by Atypical Protein Kinase C and Competition between Its Substrates. Dev Cell 49, 678–679. 10.1016/j.devcel.2019.05.002.

25. Qiu, R.G., Abo, A., and Steven Martin, G. (2000). A human homolog of the C. elegans polarity determinant Par-6 links Rac and Cdc42 to PKCzeta signaling and cell transformation. Curr Biol 10, 697–707.

26. Martin-Belmonte, F., Gassama, A., Datta, A., Yu, W., Rescher, U., Gerke, V., and Mostov, K. (2007). PTEN-mediated apical segregation of phosphoinositides controls epithelial morphogenesis through Cdc42. Cell 128, 383–397.

27. Joberty, G., Petersen, C., Gao, L., and Macara, I.G. (2000). The cell-polarity protein Par6 links Par3 and atypical protein kinase C to Cdc42. Nat Cell Biol 2.

28. Li, J., Kim, H., Aceto, D.G., Hung, J., Aono, S., and Kemphues, K.J. (2010). Binding to PKC-3, but not to PAR-3 or to a conventional PDZ domain ligand, is required for PAR-6 function in C. elegans. Dev Biol 340, 88–98.

29. Renschler, F.A., Bruekner, S.R., Salomon, P.L., Mukherjee, A., Kullmann, L., Schütz-Stoffregen, M.C., Henzler, C., Pawson, T., Krahn, M.P., and Wiesner, S. (2018). Structural basis for the interaction between the cell polarity proteins Par3 and Par6. Sci Signal 11. 10.1126/scisignal.aam9899.

30. Vargas, E., and Prehoda, K.E. (2023). Negative cooperativity underlies dynamic assembly of the Par complex regulators Cdc42 and Par-3. Journal of Biological Chemistry 299, 102749. 10.1016/j.jbc.2022.102749.

31. Beers, M., and Kemphues, K. (2006). Depletion of the co-chaperone CDC-37 reveals two modes of PAR-6 cortical association in C. elegans embryos. Development 133, 3745–3754.

32. Johnson, J.L., Erickson, J.W., and Cerione, R.A. (2012). C-terminal Di-arginine motif of Cdc42 protein is essential for binding to phosphatidylinositol 4,5-bisphosphate-containing membranes and inducing cellular transformation. Journal of Biological Chemistry 287, 5764–5774. 10.1074/jbc.M111.336487.

33. Krahn, M.P., Klopfenstein, D.R., Fischer, N., and Wodarz, A. (2010). Membrane targeting of Bazooka/PAR-3 is mediated by direct binding to phosphoinositide lipids. Curr Biol 20, 636–642.

34. Roberts, P.J., Mitin, N., Keller, P.J., Chenette, E.J., Madigan, J.P., Currin, R.O., Cox, A.D., Wilson, O., Kirschmeier, P., and Der, C.J. (2008). Rho family GTPase modification and dependence on CAAX motif-signaled posttranslational modification. Journal of Biological Chemistry 283, 25150–25163. 10.1074/jbc.M800882200.

35. Sarıkaya, S., and Dickinson, D.J. (2021). Rapid extraction and kinetic analysis of protein complexes from single cells. Biophys J 120, 5018–5031. 10.1016/j.bpj.2021.10.011.

36. Deutz, L.N., Sarıkaya, S., and Dickinson, D.J. (2023). Membrane extraction in native lipid nanodiscs reveals dynamic regulation of Cdc42 complexes during cell polarization. 1–35.

37. Holly, R.W., Jones, K., and Prehoda, K.E. (2020). A Conserved PDZ-Binding Motif in aPKC Interacts with Par-3 and Mediates Cortical Polarity. Current Biology 30, 893–898.e5. 10.1016/j.cub.2019.12.055.

38. Penkert, R.R., Vargas, E., and Prehoda, K.E. (2022). Energetic determinants of animal cell polarity regulator Par-3 interaction with the Par complex. Journal of Biological Chemistry 298, 102223. 10.1016/j.jbc.2022.102223.

39. Dong, W., Lu, J., Zhang, X., Wu, Y., Lettieri, K., Hammond, G.R., and Hong, Y. (2020). A polybasic domain in aPKC mediates Par6-dependent control of membrane targeting and kinase activity. 219.

40. Jones, K.A., Drummond, M.L., and Prehoda, K.E. (2022). Cooperative regulation of C1-domain membrane recruitment polarizes atypical Protein Kinase C.

41. Dickinson, D.J., Schwager, F., Pintard, L., Gotta, M., and Goldstein, B. (2017). A Single-Cell Biochemistry Approach Reveals PAR Complex Dynamics during Cell Polarization. Dev Cell 42, 416–434.e11. 10.1016/j.devcel.2017.07.024.

42. Lee, C.Y., Andersen, R.O., Cabernard, C., Manning, L., Tran, K.D., Lanskey, M.J., Bashirullah, A., and Doe, C.Q. (2006). Drosophila Aurora-A kinase inhibits neuroblast self-renewal by regulating aPKC/Numb cortical polarity and spindle orientation. Genes Dev 20, 3464–3474. 10.1101/gad.1489406.

43. Reich, J.D., Hubatsch, L., Illukkumbura, R., Peglion, F., Bland, T., Hirani, N., and Goehring, N.W. (2019). Regulated Activation of the PAR Polarity Network Ensures a Timely and Specific Response to Spatial Cues. Current Biology 29, 1911–1923.e5. 10.1016/j.cub.2019.04.058.

44. Benton, R., and St Johnston, D. (2003). Drosophila PAR-1 and 14-3-3 inhibit Bazooka/PAR-3 to establish complementary cortical domains in polarized cells. Cell 115, 691–704.

45. Chen, Y.M., Wang, Q.J., Hu, H.S., Yu, P.C., Zhu, J., Drewes, G., Piwnica-Worms, H., and Luo, Z.G. (2006). Microtubule affinity-regulating kinase 2 functions downstream of the PAR-3/PAR-6/atypical PKC complex in regulating hippocampal neuronal polarity. Proc Natl Acad Sci U S A 103, 8534–8539.

46. Hurov, J.B., Watkins, J.L., and Piwnica-Worms, H. (2004). Atypical PKC phosphorylates PAR-1 kinases to regulate localization and activity. Curr Biol 14, 736–741.

47. Motegi, F., Zonies, S., Hao, Y., Cuenca, A.A., Griffin, E., and Seydoux, G. (2011). Microtubules induce self-organization of polarized PAR domains in Caenorhabditis elegans zygotes. Nat Cell Biol.

48. Suzuki, A., Hirata, M., Kamimura, K., Maniwa, R., Yamanaka, T., Mizuno, K., Kishikawa, M., Hirose, H., Amano, Y., Izumi, N., et al. (2004). aPKC acts upstream of PAR-1b in both the establishment and maintenance of mammalian epithelial polarity. Curr Biol 14, 1425–1435.

49. Hannaford, M., Loyer, N., Tonelli, F., Zoltner, M., and Januschke, J. (2019). A chemical-genetics approach to study the role of atypical protein kinase C in Drosophila. Development (Cambridge) 146. 10.1242/dev.170589.

50. Cheeks, R., Canman, J., Gabriel, W., Meyer, N., Strome, S., and Goldstein, B. (2004). C. elegans PAR Proteins Function by Mobilizing and Stabilizing Asymmetrically Localized Protein Complexes. Current Biology 14, 851–862.

51. Munro, E., Nance, J., and Priess, J.R. (2004). Cortical flows powered by asymmetrical contraction transport PAR proteins to establish and maintain anterior-posterior polarity in the early C. elegans embryo. Dev Cell. 10.1016/j.devcel.2004.08.001.

52. Motegi, F., and Sugimoto, A. (2006). Sequential functioning of the ECT-2 RhoGEF, RHO-1 and CDC-42 establishes cell polarity in Caenorhabditis elegans embryos. Nat Cell Biol *8*, 978–985. 10.1038/ncb1459.

53. Schonegg, S., and Hyman, A.A. (2006). CDC-42 and RHO-1 coordinate acto-myosin contractility and PAR protein localization during polarity establishment in C. elegans embryos. Development 133, 3507–3516. 10.1242/dev.02527.

54. Nishikawa, M., Naganathan, S.R., Jülicher, F., and Grill, S.W. (2017). Controlling contractile instabilities in the actomyosin cortex. Elife 6, 1–21. 10.7554/eLife.19595.

55. Michaux, J.B., Robin, F.B., McFadden, W.M., and Munro, E.M. (2018). Excitable RhoA dynamics drive pulsed contractions in the early C. elegans embryo. J Cell Biol 217, 4230–4252. 10.1083/jcb.201806161.

56. Naganathan, S.R., Fürthauer, S., Rodriguez, J., Fievet, B.T., Jülicher, F., Ahringer, J., Cannistraci, C.V., and Grill, S.W. (2018). Morphogenetic degeneracies in the actomyosin cortex. Elife 7, 1–21. 10.7554/eLife.37677.

57. Schmutz, C., Stevens, J., and Spang, A. (2007). Functions of the novel RhoGAP proteins RGA-3 and RGA-4 in the germ line and in the early embryo of C. elegans. Development 134, 3495–3505. 10.1242/dev.000802.

58. Schonegg, S., Constantinescu, A.T., Hoege, C., and Hyman, A.A. (2007). The Rho GTPase-activating proteins RGA-3 and RGA-4 are required to set the initial size of PAR domains in Caenorhabditis elegans one-cell embryos. Proc Natl Acad Sci U S A 104, 14976–14981. 10.1073/pnas.0706941104.

59. Boyd, L., Guo, S., Levitan, D., Stinchcomb, D.T., and Kemphues, K.J. (1996). PAR-2 is asymmetrically distributed and promotes association of P granules and PAR-1 with the cortex in C. elegans embryos. Development 122, 3075–3084.

60. Hao, Y., Boyd, L., and Seydoux, G. (2006). Stabilization of cell polarity by the C. elegans RING protein PAR-2. Dev Cell 10, 199–208.

61. Hoege, C., Constantinescu, A.-T., Schwager, A., Goehring, N.W., Kumar, P., and Hyman, A.A. (2010). LGL can partition the cortex of one-cell Caenorhabditis elegans embryos into two domains. Curr Biol 20, 1296–1303.

62. Sailer, A., Anneken, A., Li, Y., Lee, S., and Munro, E. (2015). Dynamic Opposition of Clustered Proteins Stabilizes Cortical Polarity in the C. elegans Zygote. Dev Cell 35, 131–142.

63. Beatty, A., Morton, D., and Kemphues, K. (2010). The C. elegans homolog of Drosophila lethal giant larvae functions redundantly with PAR-2 to maintain polarity in the early embryo. Development 137, 3995–4004. 10.1242/dev.056028.

64. Kumfer, K.T., Cook, S.J., Squirrell, J.M., Eliceiri, K.W., Peel, N., O’Connell, K.F., and White, J.G. (2010). CGEF-1 and CHIN-1 Regulate CDC-42 Activity during Asymmetric Division in the Caenorhabditis elegans Embryo. Mol Biol Cell 21, 266–277. 10.1091/mbc.E09.

65. Liu, J., Maduzia, L.L., Shirayama, M., and Mello, C.C. (2010). NMY-2 maintains cellular asymmetry and cell boundaries, and promotes a SRC-dependent asymmetric cell division. Dev Biol 339, 366–373. 10.1016/j.ydbio.2009.12.041.

66. Aceto, D., Beers, M., and Kemphues, K.J. (2006). Interaction of PAR-6 with CDC-42 is required for maintenance but not establishment of PAR asymmetry in C. elegans. Dev Biol 299, 386–397.

67. Hung, T.J., and Kemphues, K.J. (1999). PAR-6 is a conserved PDZ domain-containing protein that colocalizes with PAR-3 in Caenorhabditis elegans embryos. Development 126, 127–135.

68. Chang, Y., and Dickinson, D.J. (2022). A particle size threshold governs diffusion and segregation of PAR-3 during cell polarization. Cell Rep 39, 110652. 10.1016/j.celrep.2022.110652.

69. Goehring, N.W., Trong, P.K., Bois, J.S., Chowdhury, D., Nicola, E.M., Hyman, A.A., and Grill, S.W. (2011). Polarization of PAR proteins by advective triggering of a pattern-forming system. Science (1979) 334, 1137–1141. 10.1126/science.1208619.

70. Zmurchok, C., and Holmes, W.R. (2021). Biophysical Models of PAR Cluster Transport by Cortical Flow in C . elegans Early Embryogenesis. 1–26.

71. Oon, C.H., and Prehoda, K.E. (2019). Asymmetric recruitment and actin-dependent cortical flows drive the neuroblast polarity cycle. Elife 8, 1–15. 10.7554/eLife.45815.

72. Wang, S.C., Low, T.Y.F., Nishimura, Y., Gole, L., Yu, W., and Motegi, F. (2017). Cortical forces and CDC-42 control clustering of PAR proteins for Caenorhabditis elegans embryonic polarization. Nat Cell Biol 19, 988–995. 10.1038/ncb3577.

73. Illukkumbura, R., Hirani, N., Borrego-Pinto, J., Bland, T., Ng, K.B., Hubatsch, L., McQuade, J., Endres, R.G., and Goehring, N.W. (2023). Design principles for selective polarization of PAR proteins by cortical flows. J Cell Biol 222. 10.1083/jcb.202209111.

74. Wang, C., Xu, H., Lin, S., Deng, W., Zhou, J., Zhang, Y., Shi, Y., Peng, D., and Xue, Y. (2020). GPS 5.0: An Update on the Prediction of Kinase-specific Phosphorylation Sites in Proteins. Genomics Proteomics Bioinformatics 18, 72–80. 10.1016/j.gpb.2020.01.001.

75. Schoentaube, J., Olling, A., Tatge, H., Just, I., and Gerhard, R. (2009). Serine-71 phosphorylation of Rac1/Cdc42 diminishes the pathogenic effect of Clostridium difficile toxin A. Cell Microbiol 11, 1816–1826. 10.1111/j.1462-5822.2009.01373.x.

76. Schwarz, J., Proff, J., Hävemeier, A., Ladwein, M., Rottner, K., Barlag, B., Pich, A., Tatge, H., Just, I., and Gerhard, R. (2012). Serine-71 phosphorylation of Rac1 modulates downstream signaling. PLoS One 7, e44358.

77. Pothula, S., Bazan, H.E.P., and Chandrasekher, G. (2013). Regulation of cdc42 expression and signaling is critical for promoting corneal epithelial wound healing. Invest Ophthalmol Vis Sci 54, 5343–5352. 10.1167/iovs.13-11955.

78. Gunaratne, A., Thai, B.L., and Di Guglielmo, G.M. (2013). Atypical Protein Kinase C Phosphorylates Par6 and Facilitates Transforming Growth Factor β-Induced Epithelial-to-Mesenchymal Transition. Mol Cell Biol 33, 874–886. 10.1128/mcb.00837-12.

79. Tokunaga, M., Imamoto, N., and Sakata-Sogawa, K. (2008). Highly inclined thin illumination enables clear single-molecule imaging in cells. Nat Methods 5, 159–161. 10.1038/NMETH.1171.

80. Gubieda, A.G., Packer, J.R., Squires, I., Martin, J., and Rodriguez, J. (2020). Going with the flow: insights from Caenorhabditis elegans zygote polarization. Philos Trans R Soc Lond B Biol Sci 375, 20190555. 10.1098/rstb.2019.0555.

81. Illukkumbura, R., Bland, T., and Goehring, N.W. (2020). Patterning and polarization of cells by intracellular flows. Curr Opin Cell Biol 62, 123–134. 10.1016/j.ceb.2019.10.005.

82. Robin, F.B., Mcfadden, W.M., Yao, B., and Munro, E.M. (2014). Single-molecule analysis of cell surface dynamics in Caenorhabditis elegans embryos. Nat Methods 11, 677–682.

83. Karslake, J.D., Donarski, E.D., Shelby, S.A., Demey, L.M., DiRita, V.J., Veatch, S.L., and Biteen, J.S. (2021). SMAUG: Analyzing single-molecule tracks with nonparametric Bayesian statistics. Methods 193, 16–26. 10.1016/j.ymeth.2020.03.008.

84. Etienne-Manneville, S. (2004). Cdc42--the centre of polarity. J Cell Sci 117, 1291–1300.

85. Sit, S.-T., and Manser, E. (2011). Rho GTPases and their role in organizing the actin cytoskeleton. J Cell Sci 124, 679–683.

86. Kwon, T., Kwon, D.Y., Chun, J., Kim, J.H., and Kang, S.S. (2000). Akt protein kinase inhibits Rac1-GTP binding through phosphorylation at serine 71 of Rac1. J Biol Chem 275, 423–428.

87. Fievet, B.T., Rodriguez, J., Naganathan, S., Lee, C., Zeiser, E., Ishidate, T., Shirayama, M., Grill, S., and Ahringer, J. (2013). Systematic genetic interaction screens uncover cell polarity regulators and functional redundancy. Nat Cell Biol 15, 103–112. 10.1038/ncb2639.

88. Mayer, M., Depken, M., Bois, J.S., Jülicher, F., and Grill, S.W. (2010). Anisotropies in cortical tension reveal the physical basis of polarizing cortical flows. Nature 467, 617– 621. 10.1038/nature09376.

89. Reymann, A.C., Staniscia, F., Erzberger, A., Salbreux, G., and Grill, S.W. (2016). Cortical flow aligns actin filaments to form a furrow. Elife 5, 1–25. 10.7554/eLife.17807.

90. Garrard, S.M., Capaldo, C.T., Gao, L., Rosen, M.K., Macara, I.G., and Tomchick, D.R. (2003). Structure of Cdc42 in a complex with the GTPase-binding domain of the cell polarity protein, Par6. EMBO J 22, 1125–1133.

91. Peterson, F.C., Penkert, R.R., Volkman, B.F., and Prehoda, K.E. (2004). Cdc42 regulates the Par-6 PDZ domain through an allosteric CRIB-PDZ transition. Mol Cell 13, 665– 676.

92. Hakoshima, T., Shimizu, T., and Maesaki, R. (2003). Structural Basis of the Rho GTPase Signaling. J Biochem 134, 327–331. 10.1093/jb/mvg149.

93. Nagai-Tamai, Y., Mizuno, K., Hirose, T., Suzuki, A., and Ohno, S. (2002). Regulated protein-protein interaction between aPKC and PAR-3 plays an essential role in the polarization of epithelial cells. Genes to Cells 7, 1161–1171. 10.1046/j.1365-2443.2002.00590.x.

94. Morais-de-Sá, E., Mirouse, V., and St Johnston, D. (2010). aPKC phosphorylation of Bazooka defines the apical/lateral border in Drosophila epithelial cells. Cell 141, 509– 523.

95. Walther, R.F., and Pichaud, F. (2010). Crumbs/DaPKC-dependent apical exclusion of Bazooka promotes photoreceptor polarity remodeling. Curr Biol 20, 1065–1074.

96. Betschinger, J., Mechtler, K., and Knoblich, J.A. (2003). The Par complex directs asymmetric cell division by phosphorylating the cytoskeletal protein Lgl. Nature 422, 326–330.

97. Dong, W., Zhang, X., Liu, W., Chen, Y. jiun, Huang, J., Austin, E., Celotto, A.M., Jiang, W.Z., Palladino, M.J., Jiang, Y., et al. (2015). A conserved polybasic domain mediates plasma membrane targeting of Lgl and its regulation by hypoxia. Journal of Cell Biology 211, 273–286. 10.1083/jcb.201503067.

98. Bailey, M.J., and Prehoda, K.E. (2015). Establishment of Par-Polarized Cortical Domains via Phosphoregulated Membrane Motifs. Dev Cell 35, 199–210.

99. Brzeska, H., Guag, J., Remmert, K., Chacko, S., and Korn, E.D. (2010). An experimentally based computer search identifies unstructured membrane-binding sites in proteins: Application to class I myosins, PAKs, and CARMIL. Journal of Biological Chemistry 285, 5738–5747. 10.1074/jbc.M109.066910.

100. Ziman, M., Preuss, D., Mulholland, J., O’Brien, J.M., Botstein, D., and Johnson, D.I. (1993). Subcellular localization of Cdc42p, a Saccharomyces cerevisiae GTP-binding protein involved in the control of cell polarity. Mol Biol Cell 4, 1307–1316. 10.1091/mbc.4.12.1307.

101. Wu, W.J., Erickson, J.W., Lin, R., and Cerione, R.A. (2000). The γ-subunit of the coatomer complex binds Cdc42 to mediate transformation. Nature 405, 800–804. 10.1038/35015585.

102. Hall, A. (1998). Rho GTpases and the actin cytoskeleton. Science (1979) 279, 509–514. 10.1126/science.279.5350.509.

103. Hannaford, M.R., Ramat, A., Loyer, N., and Januschke, J. (2018). aPKC-mediated displacement and actomyosin-mediated retention polarize Miranda in Drosophila neuroblasts. Elife 7, 1–22. 10.7554/eLife.29939.

104. Gross, P., Kumar, K.V., Goehring, N.W., Bois, J.S., Hoege, C., Jülicher, F., and Grill, S.W. (2019). Guiding self-organized pattern formation in cell polarity establishment. Nat Phys 15, 293–300. 10.1038/s41567-018-0358-7.

105. Truebestein, L., Antonioli, S., Waltenberger, E., Gehin, C., Gavin, A.C., and Leonard, T.A. (2023). Structure and regulation of the myotonic dystrophy kinase-related Cdc42-binding kinase. Structure 31, 435–446.e4. 10.1016/j.str.2023.02.002.

106. Holland, A.J., Lan, W., Niessen, S., Hoover, H., and Cleveland, D.W. (2010). Polo-like kinase 4 kinase activity limits centrosome overduplication by autoregulating its own stability. Journal of Cell Biology 188, 191–198. 10.1083/jcb.200911102.

107. Howell, A.S., Jin, M., Wu, C.F., Zyla, T.R., Elston, T.C., and Lew, D.J. (2012). Negative feedback enhances robustness in the yeast polarity establishment circuit. Cell 149, 322–333. 10.1016/j.cell.2012.03.012.

108. Bement, W.M., and Von Dassow, G. (2015). Activator-inhibitor coupling between Rho signaling and actin assembly make the cell cortex an excitable medium. Nat Cell Biol 17, 1471–1483. 10.1038/ncb3251.Activator-inhibitor.

109. Brenner, S. (1974). The Genetics Of Caenorhabditis Elegans. Genetics 77, 71–94. 10.1093/genetics/77.1.71.

110. Zeiser, E., Frøkjær-Jensen, C., Jorgensen, E., and Ahringer, J. (2011). MosSCI and gateway compatible plasmid toolkit for constitutive and inducible expression of transgenes in the c. elegans germline. PLoS One 6, 3–8. 10.1371/journal.pone.0020082.

111. Frøkjær-Jensen, C., Davis, M.W., Ailion, M., and Jorgensen, E.M. (2012). Improved Mos1-mediated transgenesis in C. elegans. Nat Methods 9, 117–118.

112. Kamath, R.S., Fraser, A.G., Dong, Y., Poulin, G., Durbin, R., Gotta, M., Kanapin, A., Le Bot, N., Moreno, S., Sohrmann, M., et al. (2003). Systematic functional analysis of the Caenorhabditis elegans genome using RNAi. Nature 421, 231–237.

113. Andrews, R., and Ahringer, J. (2007). Asymmetry of early endosome distribution in C. elegans embryos. PLoS One 2, e493.

114. Dong, Y., Bogdanova, A., Habermann, B., Zachariae, W., and Ahringer, J. (2007). Identification of the C. elegans anaphase promoting complex subunit Cdc26 by phenotypic profiling and functional rescue in yeast. BMC Dev Biol 7, 19.

115. Grimm, J.B., English, B.P., Chen, J., Slaughter, J.P., Zhang, Z., Revyakin, A., Patel, R., Macklin, J.J., Normanno, D., Singer, R.H., et al. (2015). A general method to improve fluorophores for live-cell and single-molecule microscopy. Nat Methods 12, 244–250. 10.1038/nmeth.3256.

116. Schindelin, J., Arganda-Carrera, I., Frise, E., Verena, K., Mark, L., Tobias, P., Stephan, P., Curtis, R., Stephan, S., Benjamin, S., et al. (2009). Fiji - an Open platform for biological image analysis. Nat Methods 9. 10.1038/nmeth.2019.Fiji.

117. Wollman, A.J.M., Shashkova, S., Hedlund, E.G., Friemann, R., Hohmann, S., and Leake, M.C. (2017). Transcription factor clusters regulate genes in eukaryotic cells. Elife 6, 1–36. 10.7554/eLife.27451.

118. Kusumi, A., Sako, Y., and Yamamoto, M. (1993). Confined Lateral Diffusion of Membrane Receptors. Biophys J 65, 2021–2040.

119. Syeda, A.H., Wollman, A.J.M., Hargreaves, A.L., Howard, J.A.L., Bruning, J.G., McGlynn, P., and Leake, M.C. (2019). Single-molecule live cell imaging of Rep reveals the dynamic interplay between an accessory replicative helicase and the replisome. Nucleic Acids Res 47, 6287–6298. 10.1093/nar/gkz298.

120. Kinz-Thompson, C.D., Bailey, N.A., and Gonzalez, R.L. (2016). Precisely and Accurately Inferring Single-Molecule Rate Constants. Methods Enzymol 581, 187–225. 10.1016/bs.mie.2016.08.021.

