## Supplementary material for "Atypical Protein Kinase C Promotes its own Asymmetric Localisation by Phosphorylating Cdc42 in the *C. elegans* zygote": Key Resource table

| REAGENT or RESOURCE | SOURCE | IDENTIFIER |
|---|---|---|
| **Antibodies** | | |
| Rat Anti-PKC-3 | (Tabuse et al., 1998) | N/A |
| Rabbit Anti-PAR-2 | (Dong et al., 2007) | N/A |
| Mouse Anti-PAR-3 | Developmental Studies Hybridoma Bank | P4A1; RRID: AB_528424 |
| Rabbit Anti-NMY-2 | Gift from Julie Ahringer | N/A |
| Mouse Anti-CDC-42(total) | Santa Cruz | sc-390210 |
| Rabbit Anti-pS71-CDC42/RAC1 | Invitrogen (Lot: 1186800B) | 44214G |
| Mouse Anti-HSP-60 | Developmental Studies Hybridoma Bank | HSP60, RRID: AB_10572252 |
| Mouse Anti-GFP | Roche | #11814460001 |
| Anti-rabbit-Alexa488/594/647 | Molecular Probes | RRID: AB_2576217 / RRID: AB_2534095 / RRID: AB_2535813 |
| Anti-mouse-Alexa488/594/647 | Molecular Probes | RRID: AB_138404 / RRID: AB_141672 / RRID: AB_141725 |
| Anti-rat-Alexa488/594/647 | Molecular Probes | RRID: AB_141373 / RRID: AB_141374 / RRID: AB_141778 |
| Anti-mouse-HRP | DAKO | P0447; RRID: AB_2617137 |
| Thiophosphate Ester Specific RabMAb (ab92570) | Abcam | ab92570; RRID:AB_10562142 |
| Anti-rabbit-HRP (ab136636) | Abcam | ab136636 |
| **Bacteria and Virus Strains** | | |
| E. coli: OP50: E. coli B, uracil auxotroph | CGC | WB Strain: OP50 |
| E. coli: HT115(DE3): F-, mcrA, mcrB, IN(rrnD-rrnE)1, rnc14::Tn10(DE3 lysogen: lavUV5 promoter-T7 polymerase). E. | CGC | WB Strain: HT115(DE3) |
| E. coli: HT115(DE3)- RNAi library | (Kamath and Ahringer, 2003) | N/A |
| XL10-Gold ultracompetent cells (#200314) | Agilent technologies | Cat#200314 |
| E. coli : DH5a Electrocompetent cells | Gift from Julie Ahringer | N/A |
| NEB Express Competent E coli (#C2523H) | NEB | #C2523H |

| | | |
|---|---|---|
| **Chemical, Peptides and Recombinant Protiens** | | |
| aPKC iota - Par-6 complex | this paper (produced by Shona Ellison in the Mc Donald and Goehring labs, Crick Institute) | N/A |
| PKC ζ Protein | Merk Millipore | Cat#14-525 |
| Cdc42 | Cytoskeleton | Cat#CD01 |
| MBP-CDC-42 | this paper | N/A |
| MBP-CDC-42(S71A) | this paper | N/A |
| Gateway BP Clonease II Enzyme mix | ThermoFisher | Cat#11789020 |
| Gateway™ LR Clonase™ II Enzyme mix | ThermoFisher | Cat#11791020 |
| ATP-gamma-S | Abcam | Cat# ab138911 |
| p-nitrobenzyl mesylate | Abcam | Cat#ab138910 |
| DIMBA | Cube Biotech | Cat# 18014 |
| JFX549 HaloTag | Gift from Luke Lavis (Janelia Farm) | Cat# 18014 |
| **Critical Commercial Assays** | | |
| QuickChange II XL Site directed mutagenesis kit | Agilent technologies | Cat#200521 |
| **Deposited Data** | | |
| **Experimental Models: Organisms/Strains** | | |
| C. elegans: N2 (Bristol) | CGC | WB Strain: N2 |
| C. elegans: EG6701 [ttTi4348 I; unc-119(ed3) III; oxEx1580.] | C. Frokjaer-Jensen / CGC | EG6701 |
| C. elegans: JRL1685 [gfp::cdc-42]: jraSi1[Pmex-5::gfp::cdc-42::tbb-2 3'UTR;cb-unc-119 (+)] I; unc-119(ed3) III | this paper | JRL1685 |
| C. elegans: JRL1686 [gfp::cdc-42(S71A)]: jraSi2[Pmex-5::gfp::cdc-42(S71A)::tbb-2 3'UTR;cb-unc-119 (+)] I; unc-119(ed3) III | this paper | JRL1686 |
| C. elegans: JRL1687 [gfp::cdc-42(S71E)]: jraSi3[Pmex-5::gfp::cdc-42(S71E)::tbb-2 3'UTR;cb-unc-119 (+)] I; unc-119(ed3) III | this paper | JRL1687 |
| C. elegans: JRL0016 [gfp::cdc-42; PH-GBP]: jraSi1 [Pmex-5::gfp::cdc-42::tbb-2 3'UTR;cb-unc-119 (+)] I; unc-119(ed3) III; crkEx1[pNG19: mex-5p::PH(PLC1D1)::GBP::mKate::nmy- 2UTR + unc-119(+)]; him-5 (e1490) V. | this paper | JRL0016 |
| C. elegans: JRL0031 [gfp::cdc-42(S71A); PH-GBP]: jraSi2 [Pmex-5::gfp::cdc-42(S71A)::tbb-2 3'UTR;cb-unc-119 (+)] I; unc-119(ed3) III; crkEx1[pNG19: mex-5p::PH(PLC1D1)::GBP::mKate::nmy- 2UTR + unc-119(+)]; him-5 (e1490) V. | this paper | JRL0031 |
| C. elegans: JRL0015 [gfp::cdc-42(S71E); PH-GBP]: jraSi3 [Pmex-5::gfp::cdc-42(S71E)::tbb-2 3'UTR;cb-unc-119 (+)] I; unc-119(ed3) III; crkEx1[pNG19: mex-5p::PH(PLC1D1)::GBP::mKate::nmy- 2UTR + unc-119(+)]; him-5 (e1490) V. | this paper | JRL0015 |
| C. elegans: UTX218 [gfp::cdc-42; Halo::aPKC]: jraSi1[Pmex-5::gfp::cdc-42::tbb-2 3'UTR;cb-unc- | this paper | UTX218 |

| | | |
|---|---|---|
| 119 (+)] I; pkc-3(cp328[HaloTag-GLO^PKC-3]) II; unc-119(ed3) III | | |
| C. elegans: UTX219 [gfp::cdc-42(S71A); Halo::aPKC]: jraSi2[Pmex-5::gfp::cdc-42(S71A)::tbb-2 3'UTR;cb-unc-119 (+)] I; pkc-3(cp328[HaloTag-GLO^PKC-3]) II; unc-119(ed3) III | this paper | UTX219 |
| C. elegans: UTX220 [gfp::cdc-42(S71E); Halo::aPKC]: jraSi3[Pmex-5::gfp::cdc-42(S71E)::tbb-2 3'UTR;cb-unc-119 (+)] I; pkc-3(cp328[HaloTag-GLO^PKC-3]) II; unc-119(ed3) III | this paper | UTX220 |
| *C. elegans:* JRL0034[gfp::cdc-42; NMY-2::mKate2]: jraSi1[Pmex-5::gfp::cdc-42::tbb-2 3'UTR;cb-unc-119 (+)] I; nmy-2(cp52[nmy-2::mkate2 + LoxP unc-119(+) LoxP]) I; unc-119(ed3) III | this paper | JRL0034 |
| *C. elegans:* JRL0032[gfp::cdc-42(S71A); NMY-2::mKate2]: jraSi2 [Pmex-5::gfp::cdc-42(S71A)::tbb-2 3'UTR;cb-unc-119 (+)] I; nmy-2(cp52[nmy-2::mkate2 + LoxP unc-119(+) LoxP]) I; unc-119(ed3) III | this paper | JRL0032 |
| *C. elegans:* JRL0033[gfp::cdc-42(S71E); NMY-2::mKate2]: jraSi3 [Pmex-5::gfp::cdc-42(S71E)::tbb-2 3'UTR;cb-unc-119 (+)] I; nmy-2(cp52[nmy-2::mkate2 + LoxP unc-119(+) LoxP]) I; unc-119(ed3) III | this paper | JRL0033 |
| C. elegans: NWG0047[PH-GBP]: unc-119(ed3) III; crkEx1[pNG19: mex-5p::PH(PLC1D1)::GBP::mKate::nmy- 2UTR + unc-119(+)]; him-5 (e1490) V. | (Rodriguez et al., 2017) | NWG0047 |
| *C. elegans: WM150[pkc-3(ts)]: pkc-3(ne4246) II* | (Fievet et al., 2012) | WM150 |
| C. elegans: WH423[mCherry::cdc-42(Q61L)]: Ppie-1::mcherry::cdc-42(Q61L) | (Kumfer et al., 2010) | WH423 |
| C. elegans: JA1646 [pkc-3(ts); mCherry::cdc-42(Q61L)]: pkc-3(ne4246)II; Ppie-1::mcherry::cdc-42(Q61L) | this paper | JA1646 |
| C. elegans: LP229 [NMY-2::mKate2]: nmy-2(cp52[nmy-2::mkate2 + LoxP unc-119(+) LoxP]) I; unc-119(ed3) III | (Dickinson et al 2017) | LP229 |
| C. elegans: JJ1473[NMY-2::GFP]: zuIs45 [nmy-2::NMY-2::GFP + unc-119(+)] V | (Nance et al., 2003) | JJ1473 |
| C. elegans: JA1641[pkc-3(ts); NMY-2::GFP]: pkc-3(ne4246)II; zuIs45 [nmy-2::NMY-2::GFP + unc-119(+)] V | this paper | JA1641 |
| C. elegans: JJ1579 [PAR-6 ::GFP]: unc-119(ed3) III; zuIs77 [par-6p::PAR-6::GFP; unc-119(+)] | Jim Priess | JJ1579 |
| C. elegans: KK725[nop-1]; nop-1(it142) III. | Ken Kemphues / CGC | KK725 |
| **Oligonucleotides** | | |

| | | |
|---|---|---|
| cdc-42(attb1_genomic) fwd: ggggacaagtttgtacaaaaaagcaggctcgatgcagacgatcaagtgc | IDT DNA | N/A |
| cdc-42(attb2_genomic) rev: cggggaccactttgtacaagaaagctgggtctagagaatattgcacttcttcttct | IDT DNA | N/A |
| site directed mutagenesis cdc-42 (S71A) fwd: gatcgattaaggcctctagcctatccacagaccgacgtg | IDT DNA | N/A |
| site directed mutagenesis cdc-42 (S71A) rev: cacgtcggtctgtggataggctagaggccttaatcgatc | IDT DNA | N/A |
| site directed mutagenesis cdc-42 (S71E) fwd: cgatcgattaaggcctctagagtatccacagaccg | IDT DNA | N/A |
| site directed mutagenesis cdc-42 (S71E) rev: cacgtcggtctgtggatactctagaggccttaatcgatcg | IDT DNA | N/A |
| Fwd_BglII_3'utr_CDC42:gaagatctgaacgtcttccttgtctccatgt | IDT DNA | N/A |
| Rev_KpnI_3'utr_CDC42: ggggtaccacgtaacggtgtatccggac | IDT DNA | N/A |
| **Recombinant DNA** | | |
| Plasmid: L4440 | Addgene | plasmid#1654 |
| Plasmid: pDONR221 | ThermoFisher | Cat#12536017 |
| pDONR221-CDC-42 | this paper | N/A |
| pDONR221-CDC-42(S71A) | this paper | N/A |
| pDONR221-CDC-42(S71E) | this paper | N/A |
| pCFJ210 | Addgene | plasmid#30538 |
| pJA245 | Addgene | plasmid#21506 |
| pCM1.36 | Addgene | plasmid#17249 |
| pCFJ103 | (Frøkjær-Jensen et al, 2012) | N/A |
| pCFJ04 | pMYO-3_mCherry-unc-54 | N/A |
| pCFJ90 | MYO-2_mCherry-unc-54 | N/A |
| pMAL-c5X | NEB | N/A |
| pMAL-c5X-CDC-42 | this paper | N/A |
| pMAL-c5X-CDC-42(S71A) | this paper | N/A |
| RNAi clone cdc-42 3'UTR, *cdc42(RNAi)* | this paper | N/A |
| Ahringer Feeding RNAi: *rho-1(RNAi)* | Source Bioscience | WB Clone: sjj2_Y51H4A.3 |
| Ahringer Feeding RNAi: *ced-10(RNAi)* | Source Bioscience | WB Clone: sjj_C09G12.8a |
| Ahringer Feeding RNAi: *mig-2(RNAi)* | Source Bioscience | WB Clone: sjj2_C35C5.4 |
| Ahringer Feeding RNAi: *pkc-3(RNAi)* | Source Bioscience | WB Clone: sjj_F09E5.1 |
| Ahringer Feeding RNAi: *akt-1/2(RNAi)( targets akt-1 and akt-2)* | Source Bioscience | WB Clone: sjj_C12D8.10 |
| Ahringer Feeding RNAi: rga-3*/4(RNAi)( targets rga-3 and rga-4)* | Source Bioscience | WB Clone: sjj_K09H11.3 |
| Ahringer Feeding RNAi: *mlc-5(RNAi)* | Source Bioscience | WB Clone: sjj_T12D8.6 |

| | | |
|---|---|---|
| Ahringer Feeding RNAi: *ect-2(RNAi)* | Source Bioscience | WB Clone: sjj_T19E10.1 |
| Ahringer Feeding RNAi: *let-502(RNAi)* | Source Bioscience | WB Clone: sjj_C10H11.8 |
| Ahringer Feeding RNAi: *mrck-1(RNAi)* | Source Bioscience | WB Clone: sjj_K08B12.5 |
| Ahringer Feeding RNAi: *par-6(RNAi)* | Source Bioscience | WB Clone: sjj_T26E3.3 |
| Feeding RNAi: control (ctl) | Julie Ahringer | N/A |
| **Software and Algoithms** | | |
| Matlab | Mathworks | R2017a |
| Live single molecule tracking | https://github.com/awollman/single-molecule-tools | |
| SiMPull Analysis | https://github.com/dickinson-lab/SiMPull-Analysis-Software/, | |
| Cortical flow Analysis | https://github.com/sundar07/CorticalFlow-Analysis | |
| Fiji (imageJ) | http://fiji.sc/ | N/A |
